# Task structure flexibly shifts working memory formats across cortical circuits

**DOI:** 10.1101/2025.11.28.691186

**Authors:** Jingjie Li, Ariel Ziqian Xu, Chaofei Bao, Albert Albesa-González, Liujunli Li, Claudia Clopath, Jeffrey C. Erlich

**Affiliations:** Sainsbury Wellcome Centre, University College London, London, UK; NYU-ECNU Institute of Brain and Cognitive Science at NYU Shanghai, China; NYU Shanghai, Shanghai, China; Shanghai Key Laboratory of Brain Functional Genomics (Ministry of Education), East China Normal University, Shanghai, China; Department of Bioengineering, Imperial College London, London, UK

**Keywords:** working memory, rat, cognitive control, frontal cortex, auditory cortex, electrophysiology, optogenetics, computational modeling

## Abstract

Short-term memory in delayed-response tasks is widely attributed to choice-predictive activity in frontal cortex, maintained by discrete attractor dynamics in thalamocortical circuits. Here we ask whether this mechanism extends to tasks where cues specify goals rather than actions. We trained rats on two auditory delayed-response tasks: an egocentric task, where cues mapped to movement directions, and an allocentric task, where cues specified world-centered locations. Using electrophysiology and optogenetics, we found that the egocentric task engaged early sensorimotor transformation and prospective motor planning in secondary motor cortex (M2). The allocentric task, however, relied on a delayed sensorimotor transformation: auditory cortex carried a retrospective sensory trace, whereas M2 neither encoded choice nor was required for its maintenance during the delay. Simplifying the allocentric task to a single start position rapidly reinstated early transformation and M2 dependence within the same animals. These results demonstrate that animals flexibly select the timing of sensorimotor transformation, and that this choice determines the format and cortical locus of working memory.

## Introduction

Short-term memory, the ability to maintain and act on information across a brief delay, is a foundation of adaptive behavior. The delayed-response task, in which a sensory cue instructs a future action separated by an imposed delay, has proven remarkably productive for uncovering the neural mechanisms that support this capacity. Across primates, rodents, and humans, a consistent picture has emerged: sensory cues are rapidly transformed into prospective motor plans that are maintained in frontal and premotor cortex through persistent delay-period activity^1–7^, supported by recurrent dynamics in thalamocortical circuits^8^ and causally required for accurate performance^4,5,9–11^. The success of this framework represents one of the most complete mechanistic accounts in systems neuroscience, linking computation, dynamics, and behavior^12,13^.

This success raises a natural next question: how far does the frontal motor-planning mechanism extend? In most delayed-response paradigms, subjects maintain a fixed posture and choose among a small set of stereotyped movements, creating conditions where each cue maps onto one of a few possible actions. Early sensorimotor transformation and prospective motor maintenance is a natural and efficient strategy under these conditions. But many real-world situations require remembering information that does not map straightforwardly onto a single action; instead, the same instruction demands different movements depending on context. Studying such tasks offers an opportunity to discover whether frontal motor planning is the brain’s universal solution to bridging delays^3^, or one strategy among several that can be flexibly deployed^14^.

To explore this, we designed two parallel auditory delayed-response tasks using a multi-port response wall^15^, which allowed us to vary the start position of the animal on each trial. In the egocentric (Ego) task, auditory cues specified self-centered movement directions, creating a fixed two-alternative mapping regardless of start location. In the allocentric (Allo) task, the same cues specified world-centered goal locations. Because trials began from multiple positions, a single Allo cue was associated with many possible movement vectors, while there were only two possible auditory cues across start positions. Critically, trials initiated from the center port were physically identical across the two tasks—same sounds, same movements, same contingencies—so any differences on these center-start trials must reflect the animals’ internal strategy.

This design allowed us to evaluate three candidate working-memory formats for the tasks, each with distinct neural predictions. First, animals could perform an early sensorimotor transformation and maintain a prospective motor plan in M2 (H1), as observed in standard delayed-response tasks^1,4,5^. Second, they could compute and maintain a worldcentered goal representation in M2 (H2), consistent with previous finding of allocentric representations in frontal cortex^15–17^. Third, motivated by evidence that behavioral strategy can determine whether short-term memory resides in frontal or sensory cortex^14,18^, we hypothesized that animals could defer the sensorimotor transformation entirely and maintain a retrospective memory of the auditory cue in posterior cortex (H3). H1 and H2 both predict choice-selective delay activity in M2 and causal dependence on M2 during the delay; H3 predicts a strategy that is independent of M2 during the delay. Based on the literature we expected the Ego task to be mediated by self-centered motor WM (H1). For the Allo task, we were less confident, but favored H1 or H2.

Using electrophysiology and optogenetic silencing, we found that the Ego task engaged frontal motor planning as expected: the frontal orienting field (FOF) in M2 exhibited robust, stable, choice-predictive delay activity in an egocentric reference frame, and FOF silencing impaired performance. In the Allo task, by contrast, FOF showed minimal choice-selective delay activity on center-start trials until just before the go cue, and delay-period FOF silencing had no significant effect on accuracy: arguing against both H1 and H2. Instead, auditory cortex carried a sustained sensory memory signal that covaried with task performance, consistent with H3.

Why would animals adopt a sensory memory strategy when frontal motor planning is available? The Ego and Allo tasks differ in several aspects. In Ego each sound maps to a unique action (in a self-centered frame), so animals can learn a direct cue-response map. In Allo, animals must combine the auditory cue with their start position to generate the correct action, so constructing and maintaining a precise motor plan is more demanding (i.e., combining or binding two operands, planning movements to the same side, planning movements with similar angles but different lengths) than retaining the auditory cue and computing the movement later. We formalized this intuition in a dynamical model that selects working-memory format considerate of cognitive costs. The model demonstrated that restricting the Allo task to a single start position—thereby simplifying motor planning to a binary choice—should rapidly shift animals back to a frontal strategy. We confirmed this within subject: animals with months of Allo training switched to the reduced task and within days showed faster movements, emergent M2 delay activity, and renewed sensitivity to M2 silencing. Together, these results reveal that the timing of sensory-motor transformation, and as such, working-memory format, is not a fixed property of a brain region or a training history, but a flexible cognitive strategy that animals dynamically adapt to the demands of the task at hand.

## Results

### Two auditory memory-guided tasks in different reference frames

We separately trained two groups of rats on either the Ego or the Allo task (details of subjects can be found in Table S2). The two tasks shared the same trial structure. Trials started with the illumination of a randomly selected start port. Rats poked their nose into the start port and remain fixated there for 1.5 s. During the fixation period, rats heard one of two 0.75 s sound cues (30 or 68 Hz click train) followed by another 0.75 s silent delay period. An auditory ‘go’ cue signaled the end of fixation, after which rats reported their choice. If they poked into the correct target port (described below) they could collect a reward in the reward port (Fig. 1a).

**Figure 1.**
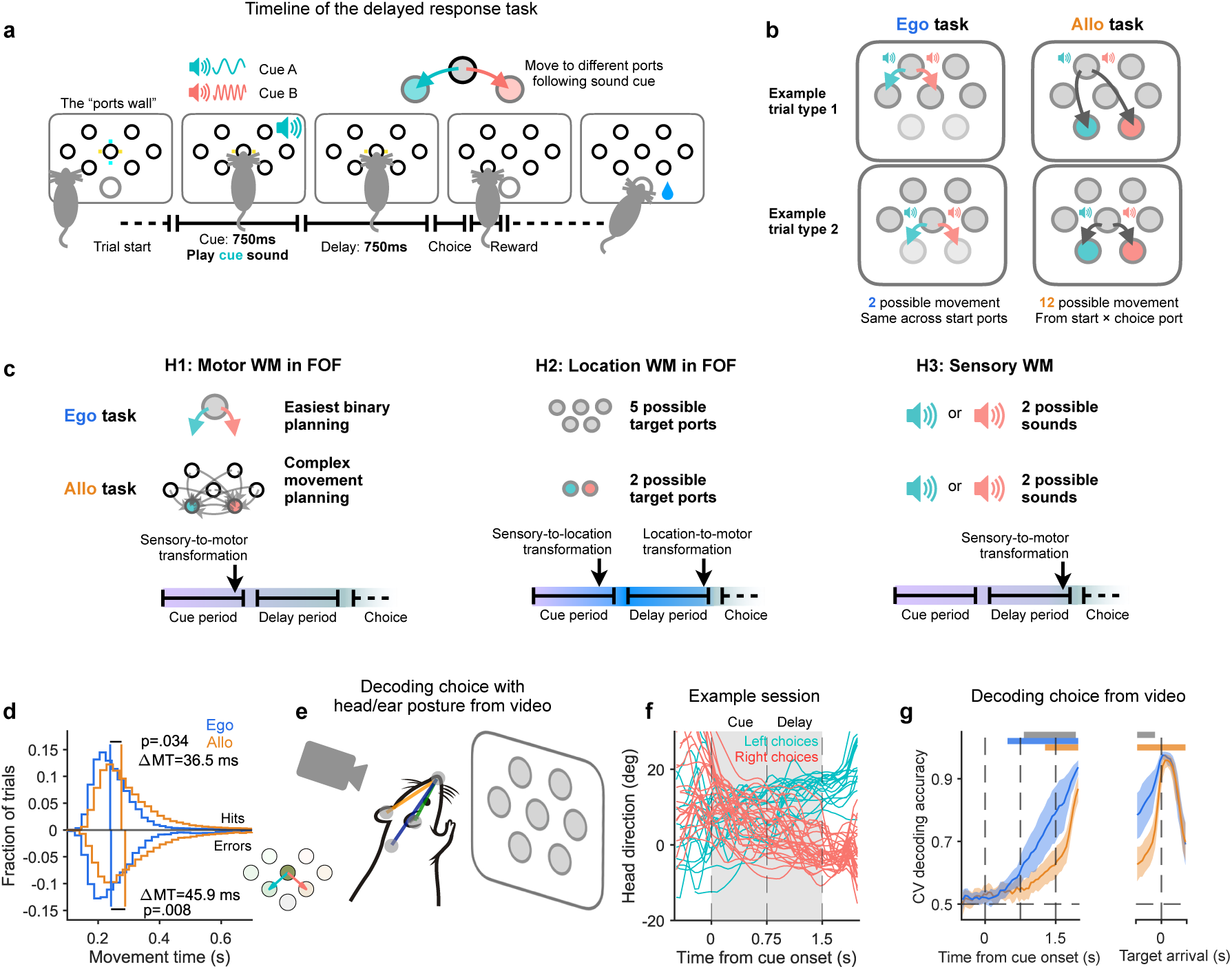
Task design and behavior quantification **a.** Schematic of the task. **b.** Mapping the sounds to the ports in the two tasks. Ego task: sounds map to movement directions. Allo task: sounds map to port locations. **c.** Given the structure of our task and prior literature, we considered three possible working-memory formats associated with representational transformations at different times in the trial (Color represent formats across time: purple for cue, gray for motor, and blue for location format) H1: A self-centered movement-vector format in the FOF. Motor planning is simple in the Ego task, but more complex in the Allo task. H2: A world-centered location-based format in the FOF ^15^ or elsewhere. This format has high WM load in the Ego task, and low load for Allo. Our task design allowed animals to adopt any of these three formats, enabling us to test whether and how representational demands steer the choice among them. H3: A sensory working-memory format in posterior cortex ^14^. This format has low WM load in both tasks, but sensory WM is generally less robust than motor WM ^19^. **d.** Histograms of movement times (MT) for the Ego and Allo tasks, for center-start trials. Correct (Top) and error (Bottom) trials. Vertical line: median of MT. **e.** Example key points and vector angles feature for video decoding analysis. **f.** Head direction trajectories from one example session. Thin lines are individual trials. The gray box highlights the fixation period. **g.** Cross-validated (CV) decoding accuracy for choice from posture features across time in a trial. Shade: 95% CI of the mean across subjects. The blue and orange thick bars at the top indicate above-chance accuracy for more than half of the subjects for the Ego and Allo task (*p <* .05, permutation test with FDR correction). The thick grey bar at the top indicates the time when the decoding accuracy was significantly different on the Ego versus Allo task (*p <* .05, two-sample *t*-test with FDR correction).

The correct target port was defined based on the task and sound cue. For the Ego task, sound A indicated that the target was to the left and sound B indicated that the target was to the right of the start port (Fig. 1b-left, Fig. S1a-left). For the Allo task, sound A indicated that the target was the bottom left port and sound B indicated that the target was the bottom right port, regardless of start position (Fig. 1b-right, Fig. S1a-right). Rats performed both tasks significantly above chance (Allo task: 69 ± 1% correct, n=21 rats, 2060 sessions, 423,115 trials. Ego task: 76 ± 2%, n=11 rats, 558 sessions, 96,523 trials). Chance performance in the task could be considered to be 1/6, since there are 6 ports available on the port wall beside the start port. However, given a task and start port, there were only two possible target ports (associated with the two sounds). An analysis of error trials showed that rats understood the structure of the tasks: most of the errors were made to the port corresponding to the unheard sound cue (Ego task: 94.9%, Allo task: 94.7%, see Fig. S1b). Performance on the Ego task was significantly higher than the Allo task (*t*(30) = 4.6973, *p* = 5.5 · 10*^−^*^5^, two-sampled *t*-test; Fig. S1g).

A key feature of our experimental design was that trials started from the center port were identical in the two tasks, since from that port the direction rule in the Ego task aligned with the location rule for the Allo task: sound A indicated that the target is down to the left and sound B indicated that the target is down to the right (Fig. 1b, trial type 2). By restricting analyses to trials starting from the center port, ‘center-start’ trials, all sensory and motor aspects were identical across the two tasks, so any behavioral (or neural) differences must reflect the task context (i.e., differences in strategy that animals are using to solve the two tasks). For the following analyses, we restricted the analysis of behavioral data to these center-start trials.

In delayed-response tasks, early movement preparation is associated with faster movements and small postural adjustments during the delay period^4,20,21^. In line with this, we found that the time from exiting the center port until reaching the chosen port (movement time, MT) was, on average, 36.5 ms shorter on the Ego task trials compared with the Allo task (Fig. 1d top; rank-sum test: *U* = 61.5, *n_Ego_* = 11, *n_Allo_* = 21, *p* = .034). From error trials, we observed a similar result: Δ*MT* = 45.9 ms (Fig. 1d bottom; rank-sum test: *U* = 48, *n_Ego_* = 11, *n_Allo_* = 21, *p* = .008) This ≈50 ms difference is consistent with the previous reports comparing movements with vs. without planning^4,22–25^, suggesting that animals in the Allo were less prepared for the movement than animals in the Ego task.

Movement planning is associated with postural adjustments^4,20,26,27^. We observed that rats tended to turn slightly toward their intended target during fixation. We quantified this by first estimating posture from video (Fig. 1e,f), and performing choice decoding on this estimated posture. We found that choice was more accurately decoded from posture during the delay period and early movement period in the Ego task than in the Allo task (Fig. 1g, left panel). Note that as rats neared the target port we were able to decode their choice perfectly in both tasks, validating our decoding method (Fig. 1g, right panel). These task driven differences in MT and posture were robust across animals (Fig. S1c,d) and were not driven by differences in performance (Fig. S1e). Together, the posture and movement time results were consistent with the hypothesis that rats in the Ego task performed their sensory-motor transformation early and used a prospective motor plan to bridge the delay period (Fig. 1c:H1). In contrast, rats in the Allo task seemed to delay the sensory-motor transformation and used a retrospective memory (of the sound or location) to bridge the early delay period (Fig. 1c:H2 or H3).

### FOF encodes prospective choice in the Ego task but not the Allo task

In memory-guided 2AFC tasks, M2 activity encodes upcoming choice and causally contributes to task performance^4,9,12,28^. Using the poke wall, we previously showed that FOF neurons encode both self-centered movement plans and world-centered target positions^15^. Given the behavioral evidence for different mnemonic strategies between the two tasks, we wanted to investigate how the contribution of the FOF might differ between them. We recorded single-unit activity from the FOF in rats performing both Ego and Allo tasks (Ego task: n = 2887 neurons, 4 rats; Allo task: n = 3325 neurons, 8 rats).

We first explored the encoding of two key task variables in individual neurons on correct trials: choice and start position. In the Ego task, start position was not essential for solving the task, since the sounds mapped to ports relative to the start position. In the Allo task, rats must combine the start position information and the sound to compute a movement vector to the target. For each neuron, we fit firing rate as a function of choice and start-position. FOF neurons exhibited tuning for task variables to different degrees in the two tasks (Example neurons: Fig. 2a-c,e-g). We found that a significantly greater proportion of neurons showed selectivity for the upcoming choice in the Ego task compared to the Allo task (Ego task: 864/2589 = .33; Allo task: 374/3186=.12; *χ*^2^(1*, N* = 5775) = 396.90, *p <* .001; Fig. 2d,h). This finding was robust across subjects (Fig. S3). Ego-task neurons showed substantial tuning for both choice and start position (Fig. 2d). Allo-task neurons showed a strong tuning for start-position axis but much weaker tuning for choice (Fig. 2h). The lack of choice coding in the Allo task was surprising given the wide consensus about the role of FOF (and frontal cortex more generally) in delayed-response tasks.

**Figure 2.**
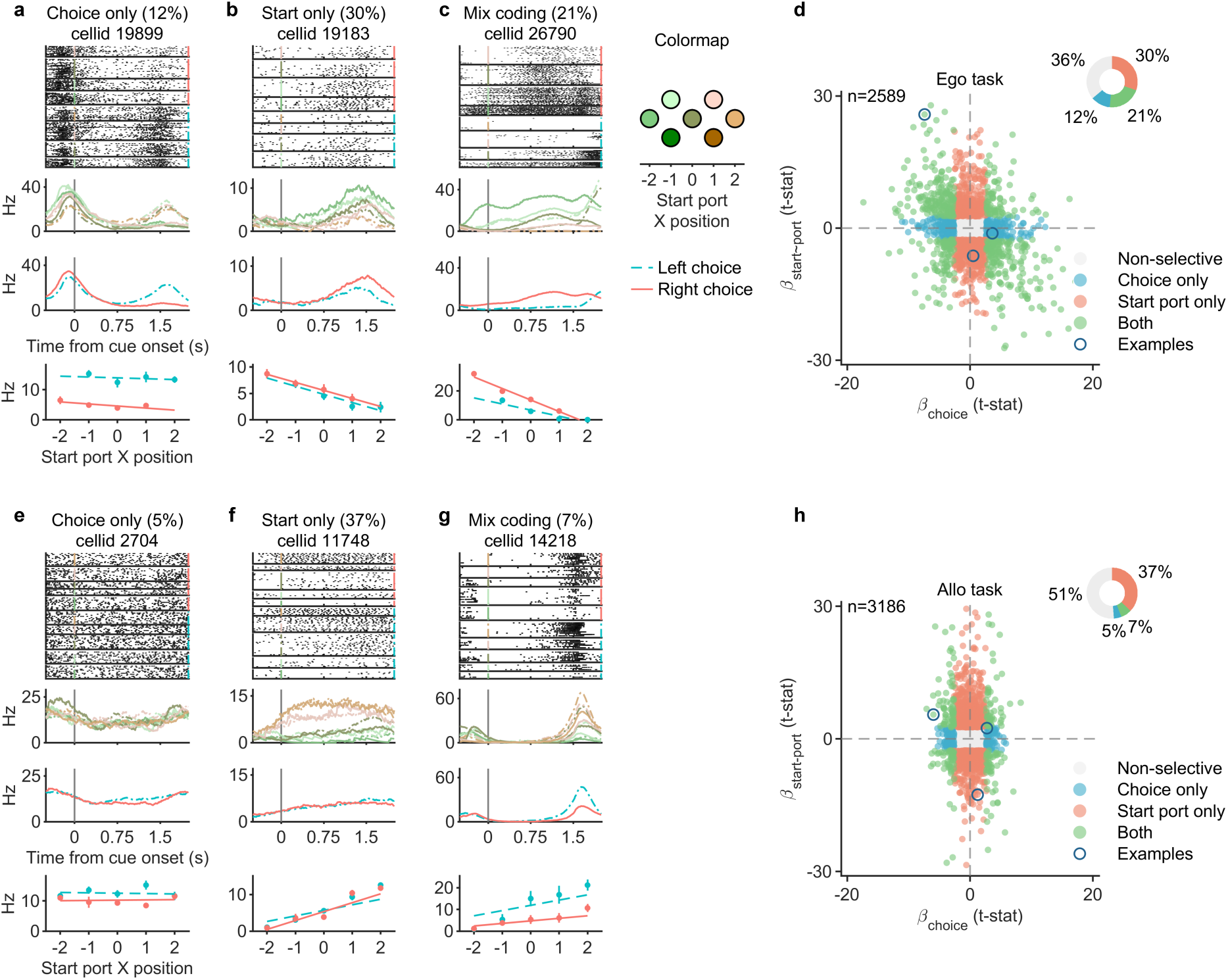
Single neuron quantification for choice and start port tuning **a-c.** Example neurons from the Ego task. The rasters and PSTHs were aligned to the sound cue onset. The rasters were sorted by different start ports (indicated by green-orange colors) and upcoming choices (red solid line for right choices, blue dashed line for left choices). The top PSTH was grouped by start port, the lower by upcoming choice. The bottom row (firing rate versus start port position) summarizes the relationship among the neural activity, start port position and choice. The lines are fits of a linear model (`Hz start_port*choice`) to the data (dots with error bars represent mean ± SEM). **d.** Distribution of coefficients of start-port (y) vs. choice (x). Black circled data show where the three example neurons shown in panel a-c are located in the scatter plot. Colors of the points indicate the significance of tuning (see legend). **e-g.** same as panel a-c, but from the Allo task **h.** same as panel d, but from the Allo task.

The single-cell selectivity analysis provided a useful first indication that the FOF plays distinct roles across the two tasks. To ensure that sensory and motor variables were fully matched, we focused subsequent analyses on center-start trials to examine the dynamics of choice coding. At the single neuron level, from the end of the cue period (0-0.75 s) until after the go signal, there was a significantly larger fraction of neurons with choice coding in Ego than in Allo animals (Fig. 3a).

**Figure 3.**
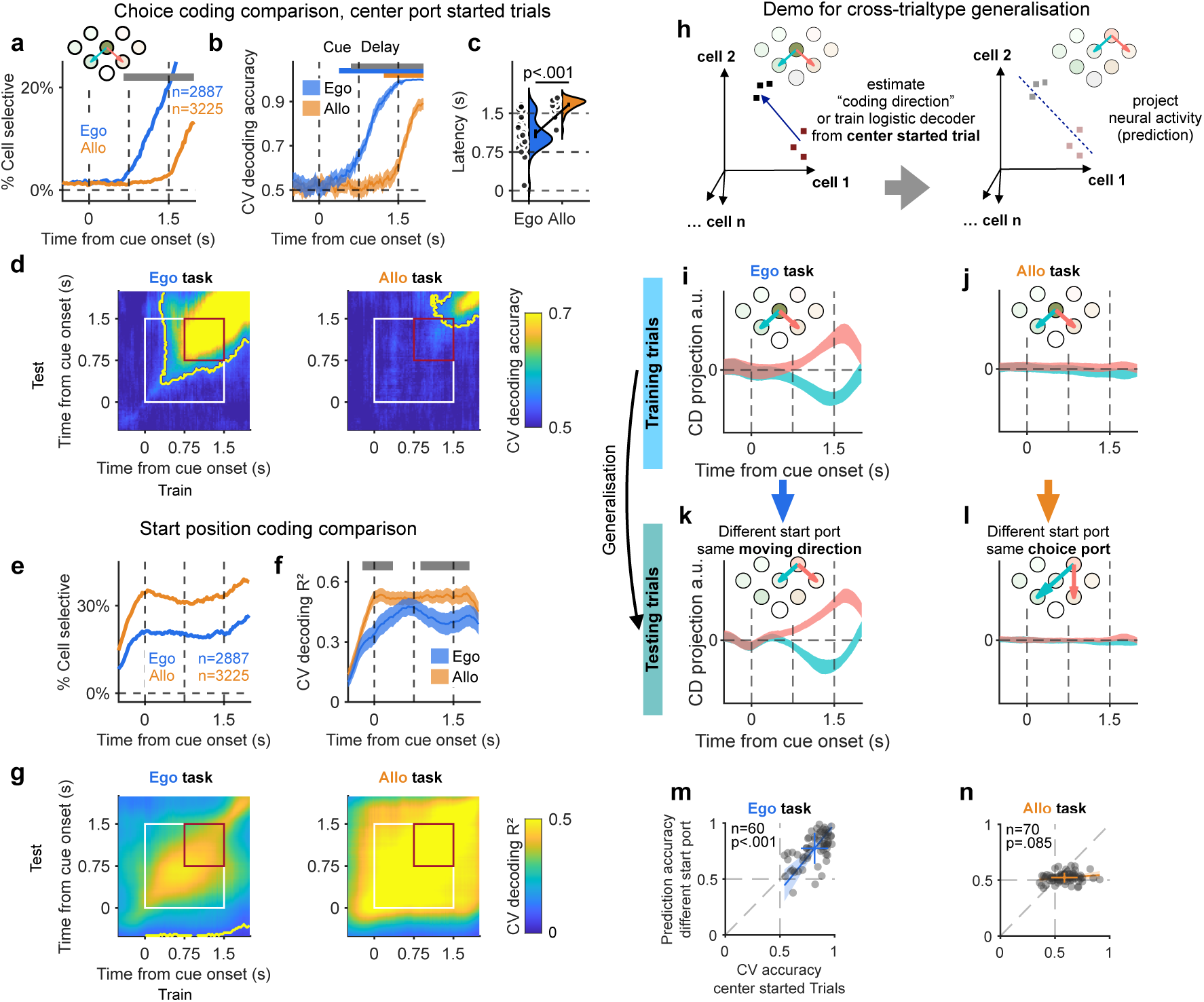
FOF population encoding of choice and start port information **a.** Development of choicedependent activity over the time course of the trial. Lines indicate % of cells at each time with significant left/right choice selectivity on center-start trials. Grey bar indicates when the % selective cells was significantly different between the Ego vs. Allo task (*p <* .01, Chi-Sq test, FDR Correction) **b.** Crossvalidated decoding accuracy for choice, across the trial, averaged over 100 pseudo populations. Shade is 99% CI The blue and orange thick bars at the top indicate above-chance accuracy for the Ego and Allo task. Grey bar indicates significant difference between Ego vs. Allo tasks (*p <* .05, permutation test with Bonferroni correction for number of time points). **c.** Distribution of choice decoding latency of each session. Dots are sessions. Error bars are 95% CI. **d.** Across-time cross-validated choice decoding accuracy averaged across 100 pseudopopulations. The decoders were trained at one time window (x-axis) and tested at another (y-axis). Contours, *p <* .01 (extreme pixel-based test). The white and red box highlight the fixation and delay period, respectively. **e.** As in (a) but for start-port selectivity over the trial. **f.** Cross-validated decoding *R*^2^ for start position, across the trial, averaged over 100 pseudo populations. Shade is 99% CI. Grey bar shows significant difference between Ego and Allo tasks (*p <* .05, permutation test with Bonferroni correction). **g.** Similar to **d** but for start port decoding. **h.** Schematic of single-trial choice coding and generalization. The coding direction (CD) for estimated from centerstart choices and then used to predict trials started from other ports. **i.** 95% CI of FOF neural activity projected to the CD (n=60 sessions) for left-out center-start trials in Ego. Blue/Red are left/right trials. **j.** same as **i** for the Allo task ((n=70 sessions). **k.** like **i**, but for trials other than center-start trials projected onto the center-start CD. **l.** as in **k**) but for the Allo task. **m.** Generalization of the choice decoder trained on center-start trials to other start positions in the Ego task. Each point represents one session: cross-validated accuracy on center-start trials (x-axis) versus out-of-sample accuracy at other start positions (y-axis). Cross: mean and SD; line and shading: TLS fit and 95% CI. **n.** same as **m** for the Allo task.

Visualizing the encoding of task variables as *proportion of selective neurons* can obscure a representation that is weak in each cell but robust in the population^29^. We employed pseudopopulation decoding, in which neurons recorded across sessions and animals are pooled and their trials combined into balanced pseudo-trials (Methods), to examine choice coding in population space. The decoding accuracy of the Ego task ramped up quickly over the trial, with a similar time-course as the rise in individually significant cells, and remained higher than the Allo task throughout the trial (Fig. 3b, p < .01, permutation test). Moreover, choice coding in the Allo task was significantly above chance only slightly before the go cue. To better understand the stability of choice coding across the trial, we trained decoders at one time in the trial and then tested them at other times (Fig. 3d). We found that decoding for choice in the Ego task was stable across the entire delay period (Fig. 3d, left panel, within red square), but absent in the Allo task (Fig. 3d, right panel, within red square).

The strong and stable choice coding in the Ego task is consistent with previous findings in delayed-response tasks in M2^15,30^. However, the absence of choice coding in the Allo task was surprising. This provided further evidence that animals in the Allo task were delaying the sensory-motor transformation until the end of the delay period. Because binary decoding tests for any separable neural signal between the two conditions, the absence of choice coding on center-start trials means no upcoming choice information is present in the FOF, regardless of the coordinate system or reference frame used to express it. This rules out both H1 (FOF motor WM) and H2 (FOF location WM) for the Allo task.

We also examined the dynamics of the start position coding in FOF at both the single cell and population level. At the single-cell level, throughout the trial, we observed more cells significantly encoding the start position in the Allo task than in the Ego task (Fig. 3e). Decoding performance for start position from the pseudopopulation in the Allo task was strong and stable for the entire fixation period. While decoding for the Ego task was strong, it was significantly worse than the Allo task during most of the early cue and delay period (Fig. 3f, p < .01, permutation test). We also found that, in both Ego and Allo tasks, start position at any time during fixation could be decoded from any other time during fixation, indicating a stable start-position code (Fig. 3g), consistent with previous results^15^.

One limitation of pseudopopulation analysis is that it may miss true encoding that depends on correlations between neurons, masked by large trial-by-trial fluctuations^31,32^ To address this, we performed moment-by-moment leave-one-trial-out decoding of start position and choice at the single-session level for both Ego and Allo. The results were consistent with both single-cell and pseudopopulation analyses: start position decoding was strong in both tasks but choice decoding was significantly stronger and earlier in the Ego task (Fig. S4). On average, the earliest time with above-chance single-trialdecoding in the Ego task was before the go cue (From cue onset: 1.10 ± 0.08 s, n=51 sessions), which was much earlier than the Allo task, which on average emerged after the go cue (1.67 ± 0.04 s, n=44 sessions, comparison: *p <* .001, permutation test, Fig. 3c). Taken together, the analysis of FOF activity on center-start trials showed clear differences between the two tasks. In the Ego task, FOF activity predicted upcoming choice, while in the Allo task there is a noticeable absence of choice activity until just before the go cue. Together with our demonstration of strong start-position coding in the Allo task (and the lack of correlation between start and choice coding in the Allo task or between choice coding and behavior performance, Fig. S4f-h), our results suggest an FOF-independent strategy for delay-period choice maintenance in the Allo task.

### In the Ego task, planning is in a self-centered frame that generalizes across start positions

For center-start trials in the Ego task, FOF persistently codes for choice during delay period. Does this code generalize to other start positions (Fig. 1c, H1)? To visualize choice encoding at the population level, we analyzed FOF activity in an activity space where each dimension corresponds to activity of one simultaneously recorded neuron. We estimated a ‘coding direction’^5,33^), *CD*, from the delay period of center-start trials along which activity maximally discriminated future choice (Fig. 3h, Methods). When left-out trials were projected to the coding direction, FOF population activity in the Ego task strongly separated by choice (Fig. 3i). However, in the Allo task, the same approach failed to show clear discrimination between choices (Fig. 3j). This recapitulates our earlier finding (Fig. 3b), but at the single-trial level, rather than in the pseudopopulation.

We then asked how the coding direction defined from center-start trials generalized to other start positions. In the Ego task, the coding direction generalized across start positions (Fig. 3k); Fig. S5a). This result confirmed our hypothesis: in the Ego task rats use an FOF dependent self-centered motor WM format that is robust to variation in start position. The coding direction also separated choices on single-option trials, in which animals started from the middle-left or middle-right port and only one response was available. As expected, in the Allo task, using the uninformative coding direction derived from center-start trials resulted in consistently poor decoding for other start positions (Fig. 3l; Fig. S5b).

In the Ego task, a decoder trained on center-start trials also decoded choice accurately at other start positions (Fig. 3m, *p*<.001, one-sided bootstrap test for TLS slope > 0.). This result confirmed that in the Ego task, the activity pattern in FOF was consistent for planning leftward/rightward movement across different start positions, i.e. planning in a self-centered motor-vector format. In the Allo task, choice decoding was weak on center-start trials and remained near chance at other start positions (Fig. 3n, p=.085, bootstrap). These results support a self-centered motor-vector representation in FOF during the Ego task.

### Silencing FOF during the delay impairs performance in the Ego task, but not the Allo task

The electrophysiological results imply that FOF supports a movement planning strategy for the Ego, and does not support the Allo task. To causally test this we expressed an inhibitory opsin, stGTACR2^34^, in pyramidal neurons bilaterally in FOF and implanted fibers for laser delivery (Fig. S11d,e). We interleaved FOF sessions with control sessions where we connected to the sham fiber (Method, laser light was delivered through a sham fiber above the skull; Fig. 4a left). In each session, we delivered laser on 25% of trials during either the cue or delay period (Fig. 4b).

**Figure 4.**
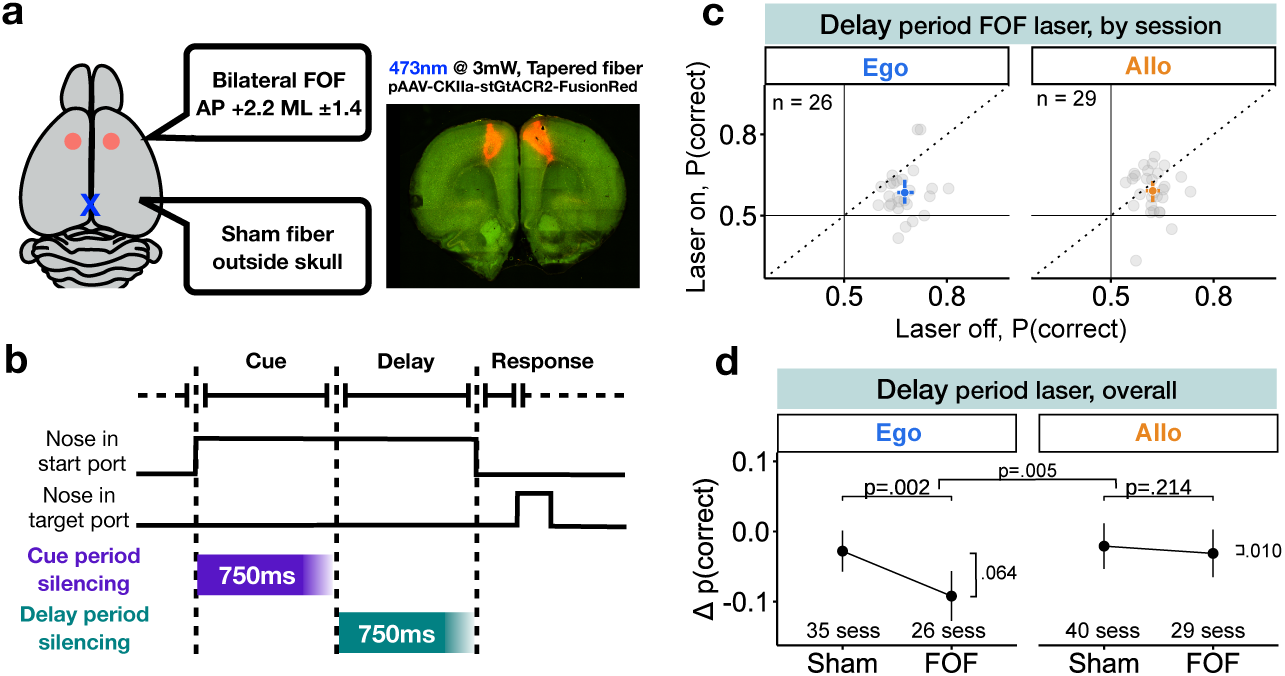
Effects of optogenetic silencing the FOF. **a.** Illustration for virus injection and fiber implantation location at bilateral FOF and a sham fiber implantation above the skull (Left). Coronal sections of a rat brain expressing stGtACR2-FusionRed (Right). **b.** Timeline of bilateral silencing of FOF during two different trial epochs (cue and delay); see Fig. S6 for cue period results. **c.** Performance modulation for delay period laser effect. Each FOF session is one point. X and Y axes show the fraction of correct choices on laser-off and laser-on trials, respectively. Crosses reflect mean and 99% bootstrapped CI of the mean across sessions. **d.** Comparison of delay period silencing between Sham and FOF sessions. Y axis: fraction of correct choices for laser-on-trials - laser-off-trials. Dot and error bar reflect mean and 95% bootstrapped CI of the mean across sessions.

For each session, we compared the performance, p(correct), between laser-off and laser-on trials during the cue and delay epochs. During the cue period, there was a small decrease in performance in both tasks (Fig. S6e). In contrast, during the delay period, nearly all Ego-task sessions exhibited a marked reduction in performance on laser-on trials relative to laser-off trials, whereas in the Allo there was no apparent difference between performance on laser-on vs. laser-off trials (Fig. 4c).

To more rigorously quantify the effect of silencing the FOF, we fit logistic generalized linear mixed-effects models (GLMMs) to quantify the impact of optogenetic inactivation of FOF on performance compared to sham sessions, during the cue and delay periods of the Ego or Allo task, separately (cue/delay × Ego/Allo = 4 models). For succinctness, *p* values of likelihood ratio tests are reported in the corresponding figures. All details of the GLMMs are reported in methods and statistical appendix.

Silencing FOF in the cue period caused small impairments in both tasks (Fig. S6f). Silencing FOF in the delay period caused a larger performance impairment in the Ego task (Fig. 4d left). Consistent with weak choice coding for the Allo task in FOF, we didn’t observe significant performance impairment for delay period FOF silencing for the Allo task (Fig. 4d right). Thus, silencing FOF during the delay period significantly impaired the Ego task but not the Allo task performance. Moreover, the FOF silencing effect was also observed at the individual subject level (Fig. S6c,d; Table S3). We also directly compared the optogenetic effect across tasks. For the cue period the effect of silencing was not significantly different between the two tasks (*p* = .569, LRT). For the delay period, there was a significant difference (*p* = .005, LRT). Thus, during the short-term memory period (i.e. delay period), the silencing FOF generated a significantly greater impairment in performance in the Ego versus the Allo task. Detailed statistical results can be found in Table S4 and the statistic appendix.

One potential confound is that the overall performance on the Allo task is worse than the Ego task, which could make it hard to detect a significant effect of silencing. To address that concern, we re-ran our analysis using a subset of sessions from each task within the same range of performance (60 to 67 % correct). Even in this subset of the data, silencing FOF impaired Ego performance significantly more than Allo performance (p=.043, Fig. S6h). This finding is especially striking given that harder tasks are typically more sensitive to perturbations^35^.

Choice-selective FOF activity emerged near or after the go cue in the Allo task, and we did not silence FOF during this epoch. Our results therefore do not exclude a post-go contribution of FOF in the Allo task. However, earlier work has shown that silencing the FOF after the go cue does not produce a behavioral effect in typical memory-guided tasks^9,36^, putatively because the movement signals are widely distributed across the brain at that point^37^.

### Auditory cortex carries a retrospective sensory trace in the Allo task

We provided strong evidence that FOF supports the Ego task through movement planning in the delay period. For the Allo task, having ruled out a role for FOF, we set out to find an area with task-related delay-period activity. We recorded from several candidate areas: auditory cortex (AC), which could support a retrospective sensory strategy^38^ (H3); hippocampus^39,40^ and retrosplenial cortex^41,42^ (RSC), which could support a locationbased strategy (H2); medial prefrontal cortex (mPFC)^43–46^ and the superior colliculus (SC)^47^ which could support abstract non-spatial short term memory. Of all the explored regions, the weight of evidence indicated that the auditory cortex was the most likely candidate for supporting Allo task performance (Fig. S9b).

Consistent with previous AC recordings^48^, AC neurons showed strong cue-evoked responses to the 30 Hz and 68 Hz click-train cues (Fig. 5a, left), with some neurons responding to individual clicks within the trains. We also observed neurons that were selective for the sound cue during the silent delay period (Fig. 5a, middle). On error trials, animals chose the target corresponding to the opposite cue. These trials allowed us to distinguish whether delay-period activity encoded the presented sound or the animal’s upcoming choice. We found that delay-period activity encoded the sound in the Allo task but the upcoming choice in the Ego task (Fig. 5a).

**Figure 5.**
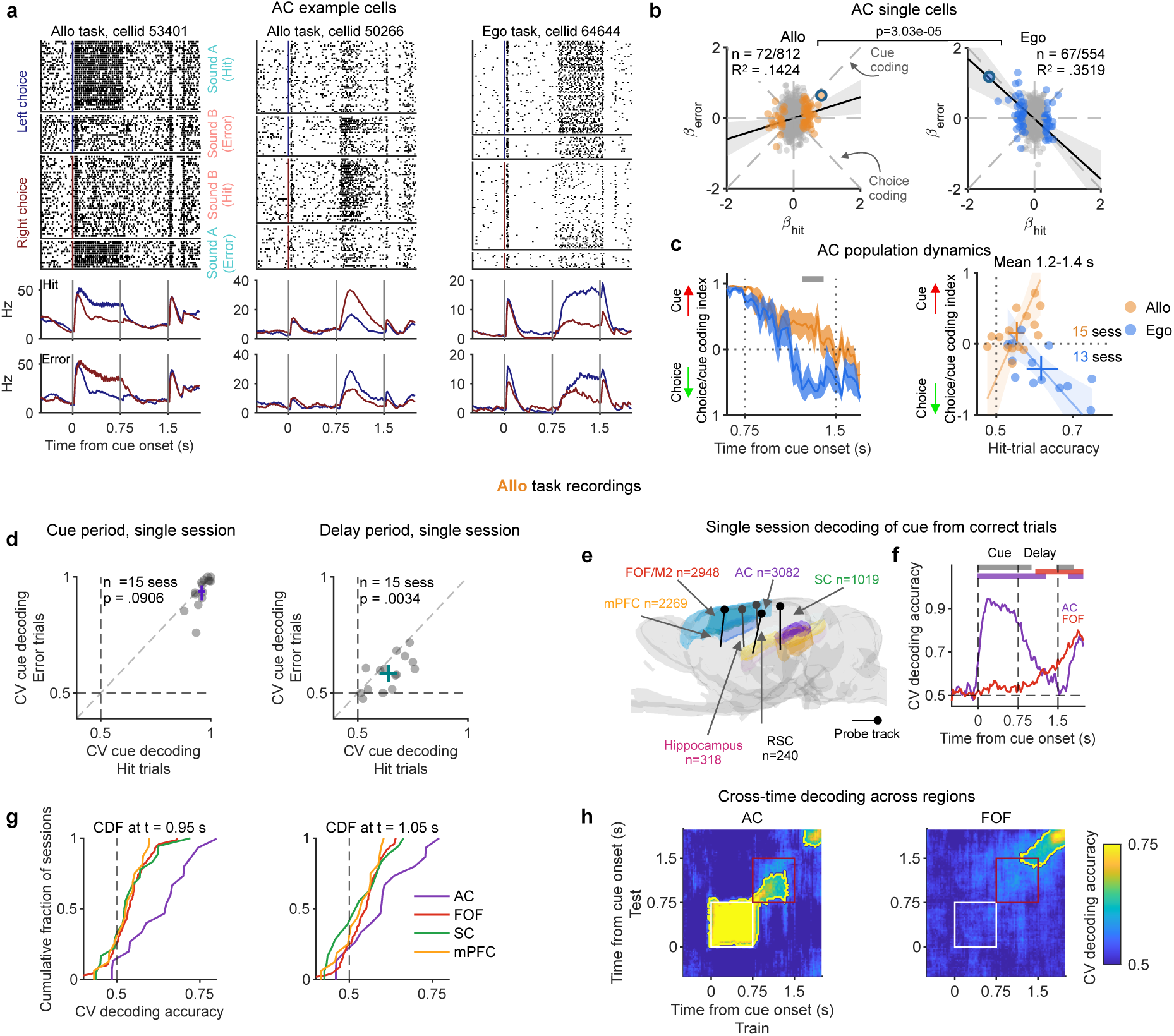
Auditory cortex (AC) activity in the Allo and Ego task. **a.** Example AC neurons recorded in the Allo (left and middle) and Ego (right) tasks, aligned to sound cue onset. Upper/lower raster groups show left/right choices, each further divided by sound cue and labeled by trial outcome. Upper/lower PSTHs show correct/error trials. Sound A/B instructed left/right choices, respectively. Vertical lines indicate cue onset, cue offset and the go cue. **b.** Comparison of coefficients of single AC neuron cue coding on correct versus error trials in the Allo (left) and Ego (right) tasks. Coefficients were estimated from z-scored spike counts in a 300-ms window centered at 1.3 s after cue onset. Each point is a single neuron. Colored points indicate significant cue coding on correct trials; the delay-period examples from panel a (middle and right) are circled. **c.** AC population cue/choice coding index: positive/negative values indicate cue/choice coding. Left: mean across the top half of sessions ranked by correct-trial decoding accuracy (1.2–1.4 s). Shade: SEM; grey bar: task difference (*p <* .05, cluster permutation test). Right: session means (1.2–1.4 s) versus correct-trial accuracy, all sessions. Crosses: mean and 95% bootstrap CI. Orange/blue lines and shading: total least-squares fits for Allo/Ego sessions with 95% CI. **d.** Comparing cue decoding on correct and error trials using 10-fold cross-validation during the cue (left-purple) and delay (right-cyan). Points and error bars indicate mean and 95% CI. **e.** Schematic of recorded regions and probe tracks: frontal orienting field (FOF), medial prefrontal cortex (mPFC), auditory cortex (AC), hippocampus, retrosplenial cortex (RSC) and superior colliculus (SC) **f.** Averaged cue (or choice) decoding accuracy for correct trials across sessions in FOF and AC. The purple and red thick bar at the top indicate when the accuracy was significantly above chance for each region (*p <* .01, permutation test). Grey bar indicates when AC accuracy was significantly higher than FOF (*p <* .05 with FDR test, permutation test). **g.** Cumulative distribution of cue/choice decoding accuracy across sessions (correct trials) in each region, shown for two representative times after cue onset (Left: 0.95 s and right: 1.05 s). **h.** Cross-time decoding of cue (or choice) on correct trials. Similar to Fig. 3d. The white and red boxes highlighted the cue and delay period. Contours, *p <* .01 (extreme pixel-based test)

To quantify this distinction, we compared cue selectivity on correct and error trials across individual neurons (Methods). Among neurons with significant cue coding on correct trials, delay-period activity on error trials continued to encode the presented sound in the Allo task (TLS slope = .299, 95% bootstrap CI [.071, .553]). In contrast, in the Ego task, activity on error trials followed the animal’s choice rather than the presented sound (TLS slope = -.852, 95% bootstrap CI [-1.375, -.412]; Fig. 5b). These relationships differed significantly between tasks (*p* = 3.03 × 10*^−^*^5^), across time and individual animals (Fig. S8a,b).

We next examined cue and choice coding in AC population activity using a cue/choice coding index (Methods). Positive values indicated encoding of the presented sound, whereas negative values indicated encoding of the animal’s choice. The index was positive in both tasks during sound presentation, but after cue offset, it remained positive in Allo (.157, 95% CI [.035, .308]) and became negative in Ego (−.349, 95% CI [-.518, -.174]; task difference: *p <* .001; Fig. 5c; Fig. S8c). In the Allo task, cue decoding was similarly strong on correct and error trials during sound presentation (correct: .96 [.94, .97]; error: .94 [.89, .96]; *p* = .091), but was significantly weaker on error trials during the delay (correct: .64 [.60, .68]; error: .58 [.55, .62]; *p* = .0034; Fig. 5d), linking the strength of the retrospective representation to behavioral performance. Thus, AC carried a cue-aligned sensory trace during the Allo task delay period but carried prospective choice information during the Ego delay. The Ego pattern closely resembles the transition from stimulus coding to later choice-related activity reported in a conventional auditory delayed-response task^49^, consistent with the possibility that late choice activity in AC reflects top-down input from decision or motor-planning circuits.

We also compared the time course of cue decoding accuracy between AC and FOF on correct trials only, where the cue and choice have a one-to-one mapping. Decoding in AC rose immediately after cue onset, reached near-perfect accuracy throughout the cue epoch, and remained above chance well into the delay. In contrast, decoding in FOF emerged only very late in the delay (Fig. 5f). We then quantified decoding strength across all four regions at two representative time-points during the delay. The decoding from AC recordings was significantly higher than FOF, SC, and mPFC (t = 0.95 s: *p* < .01 for all comparisons; t = 1.05 s: *p* < .05 for all comparisons, ranksum test, see statistical appendix; Fig. 5g). Together, these results indicate that AC provides the strongest and earliest population-level representation of the cue in the Allo task during the delay period. We next tested whether cue coding during the delay occupied the same or a distinct subspace compared with the cue period. Cross-temporal decoding, restricted to correct trials to avoid confounds between cue and choice, showed that AC representations were highly stable within the cue epoch, but after cue offset transitioned into a distinct subspace that remained stable throughout the delay (Fig. 5h). This demonstrates that delay-period memory was not simply a persistence of cue responses, but was recoded into a separate representational space, consistent with rotational dynamics previously described in auditory cortex^50^. Such a cue-to-delay subspace shift was unique to AC: in FOF, SC, and mPFC, decoding either emerged only late in the delay or remained weak. At the population level, session-wise decoding confirmed that AC provided stronger memory-related signals compared to FOF, mPFC and SC, especially during early delay period.

### A dynamical model of cost-sensitive working memory

Given the vast literature implicating the broad frontal premotor (secondary motor) area M2 in similar memory-guided tasks—particularly mouse anterior lateral motor cortex (ALM) in directional licking and the rat frontal orienting field (FOF) in head-orienting movements^12,13,51^—we were surprised to discover that performance in the Allo task was not supported by FOF delay period activity. However, it is well-known that frontal cortex dependent premotor planning is not the only form of working-memory^14^. For example, in delayed-comparison tasks, sensory features during the delay can be observed in sensory and prefrontal cortices^38,52,53^. Our recordings from AC suggest that animals in the Allo task used a sensory memory strategy. Why were animals using a motor strategy in the Ego task and a sensory strategy in the Allo task? We posited that animals make a tradeoff when considering a mnemonic strategy in order to reduce overall costs^54,55^.

The cost of a mnemonic strategy could arise from maintaining its representation, the precision needed to distinguish movements, or the different steps of transformation between sensory, location, and motor representations (Fig. 1c:H2). Our model relies on two assumptions. First, that other things being equal motor planning is the preferred strategy for delayed-response tasks. Second, the FOF supports planning *strictly* in an egocentric reference frame (Fig. 3i,k,m)^15^. In Ego, each cue specifies a movement direction regardless of start position, and early planning offers quicker movements and better preparedness for their sensory consequences^56^. Planning in a self-centered, premotor coordinate frame is also an evolutionarily ancient form of cognition^57–59^. In Allo, constructing a motor plan requires combining the cue-defined target with the start position and distinguishing movements with similar angles but different lengths. Constructing and maintaining this plan may therefore be more costly than retaining the auditory cue and computing the movement later. Together, these two assumptions mean that for Ego, motor planning is more efficient, but for Allo, sensory working memory is more efficient (Fig. 6b).

**Figure 6.**
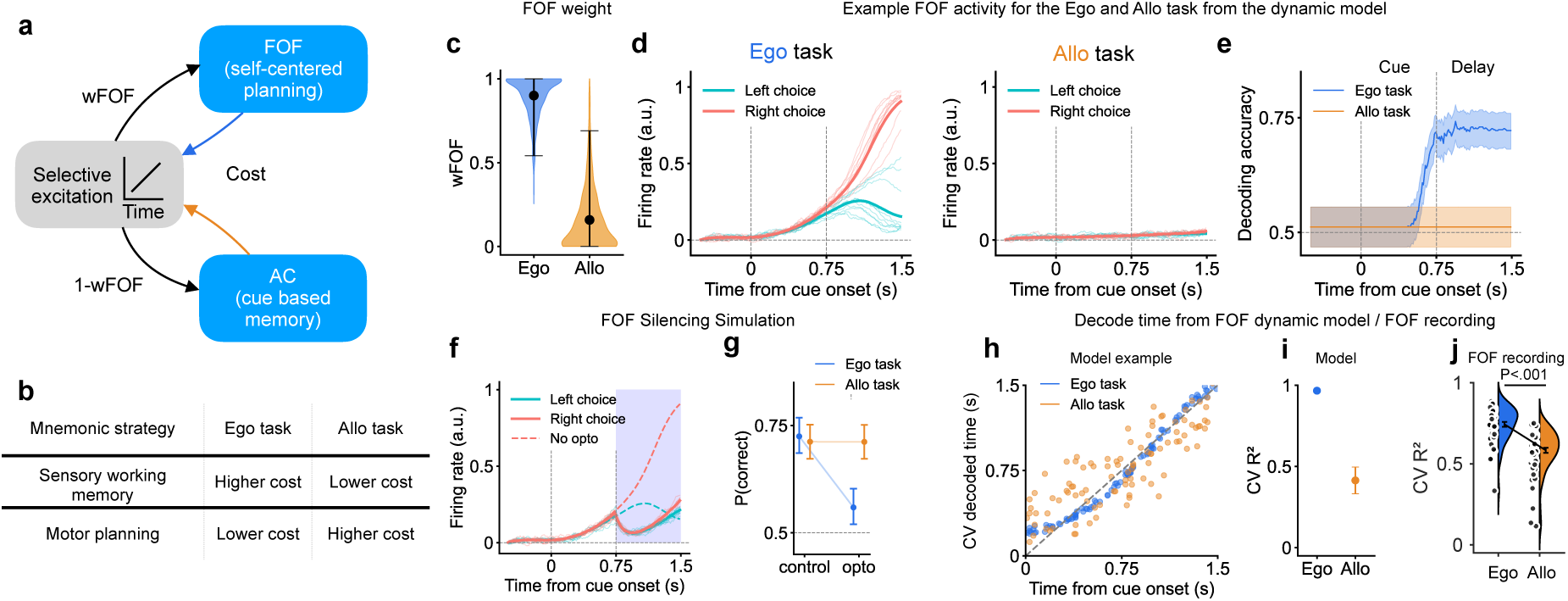
Dynamical model of cost-sensitive WM format selection. **a.** Schematic of the model. The selective excitation dynamically shifts between FOF and AC depending on the cost of the strategy. **b.** Hypothesized cost relationship between mnemonic strategies in the two tasks. **c.** Distribution of *w*_FOF_ (weight of using FOF network) in ‘expert’ models from 1000 simulations. Error bars represent 95% of distribution. **d.** Example activity from the FOF node in the Ego task (left) and the Allo task (right). Solid line represent the mean activity for each choice. Light traces represent activity from selected example trials. **e.** Time-course of decoding accuracy in the FOF network for left/right choice, colored differently for Allo and Ego tasks. Lines represent the mean, shaded areas are 95% CI. **f.** Example FOF node activity during silencing in the Ego task during the delay period. Similar to d, solid traces show the mean activity for future left/right choices, while dashed lines represent the mean activity without silencing. Shaded areas indicate the silencing period (0.75 to 1.5s). **g.** Effect of simulated optogenetic silencing the FOF node for both Ego and Allo tasks. Circles and error bars represent the mean and 95% CI, respectively. **h.** Examples of decoding results for time during the trial from Ego and Allo simulations. **i.** Time decoding mean *R*^2^ for the Allo and Ego tasks from FOF model activity. Error bar represent SEM. **j.** Time decoding *R*^2^ for the Allo and Ego tasks from FOF recording data. Circles for individual recording sessions. Violins show kernel density estimation for *R*^2^ across all sessions. Error bars: SEM

We instantiated this hypothesis in a dynamical model with three modules: one representing the FOF, one representing AC, and finally a module that provides selective excitation. We instantiated the FOF module with a bistable attractor in order to replicate the results from the optogenetics experiments. The AC module was a fixed performance module (did not have any dynamics). For simplicity, we simulated costs associated with maintaining each memory format (Methods). The selective excitation module drove either the FOF or the AC module based on these costs. Its relative weight, *w*_FOF_, shifted toward the lower-cost strategy through gradient descent subject to noise *σ_w_*, and a learning rate *α*. A similar rule could also produce similar strategy selection if costs arose from movement resolution or different transformations, provided they followed the relationship in Fig. 6b. We also implemented a richer dynamical model that could recapitulate the results more completely (including the results from AC and the late transfer of information from AC to FOF), but used the same excitatory drive mechanism for switching between mnemonic strategies (Fig. S13)

By design, in the dynamical simulations, the model heavily favors the FOF (*w*_FOF ≈_ 1) for the Ego task but favors the AC for the Allo task (Fig. 6c). In the Ego task, the FOF module received excitation and showed selective activity for upcoming left/right choice (Fig. 6d left). In the Allo task, there was no selective activity (Fig. 6d right). This simulation result recapitulated our FOF recordings (Fig. 2). We were able to decode choice from the FOF module during the delay period, for the Ego task but not the Allo task (Fig. 6e). This replicated our findings from FOF recording in real animals during the delay period (Fig. 3b). Moreover, silencing the FOF module during the delay period inhibited the selective activity (Fig. 6f) and impaired task performance for the model in the Ego task but not the Allo task (Fig. 6g) similar to our optogenetic results (Fig. 4).

In the dynamical model, the selective excitation starts at the time of the sound and ramps until go cue (which is the end of the trial) consistent with observations of ramping dynamics in frontal cortex^60,61^. This generated selective activity in FOF module that qualitatively matched the time course of real FOF activity. We realized that the presence of ramping excitation in Ego but not Allo, would predict that decoding of time in the trial should be stronger in Ego vs. Allo. We quantified the time decoding performance as R^2^ between estimated time and actual time during the trial: the FOF module had higher time-in-trial decoding for the Ego task than the Allo task (Fig. 6h,i). This finding was then confirmed in real data (Fig. 6j), providing evidence for selective excitation as a mechanism for switching mnemonic strategies.

### Rapid reinstatement of FOF dependence following the Allo-to2AFC task transformation

In the dynamical model, the selection of mnemonic strategy flexibly adapts to cost. An alternative model is that the selection of mnemonic strategy is a consequence of learning trajectory: since the Allo and Ego animals were trained on different tasks, they may have adopted different strategies through learning. Our dynamical model would rapidly switch to a FOF dependent motor planning strategy if the task was simplified so that the egocentric strategy fell below the cost of using the auditory mnemonic strategy. In contrast, the learning hypothesis would predict that Allo animals would continue to use an FOF independent strategy in the case of a simplified task. We simulated this idea in our model by removing all start positions except the center port (Fig. 7a). In this reduced task, the egocentric strategy has a cost of 1, and the selective excitation to FOF, w_FOF_, increased within hundreds of trials (Fig. 7b). An increase in w_FOF_ predicts several behavioral and neural consequences: faster movement times, performance impairments with silencing FOF, and appearance of delay period activity encoding upcoming choice. We tested these predictions using a within-subject experiment. Animals, who had trained for months on the Allo task, were transferred to a reduced version of the task, 2AFC, where all trials were center-start trials (Fig. 7a, see Fig. S10a for details). We compared center-start trials between Allo and 2AFC and verified the predictions of the model. After task transformation, movement times began to speed up, with a clear reduction emerging by day 2 (Fig. 7c, Table S7). Overall, movement times were 25.6 ms faster after the rapid task reduction for correct trials (Fig. S10g from n=12 rats, Allo task: Median MT = 247.2 ms [244.0 249.9] 95% CI, 2053 trials; reduced 2AFC task: 221.6 ms [220.3 222.6] 95% CI, 9259 trials, *p* < .001, LRT). These faster responses provided behavioral evidence that animals were rapidly shifting to a motor-planning strategy: moving the sensory-motor transformation to earlier in the trial.

**Figure 7.**
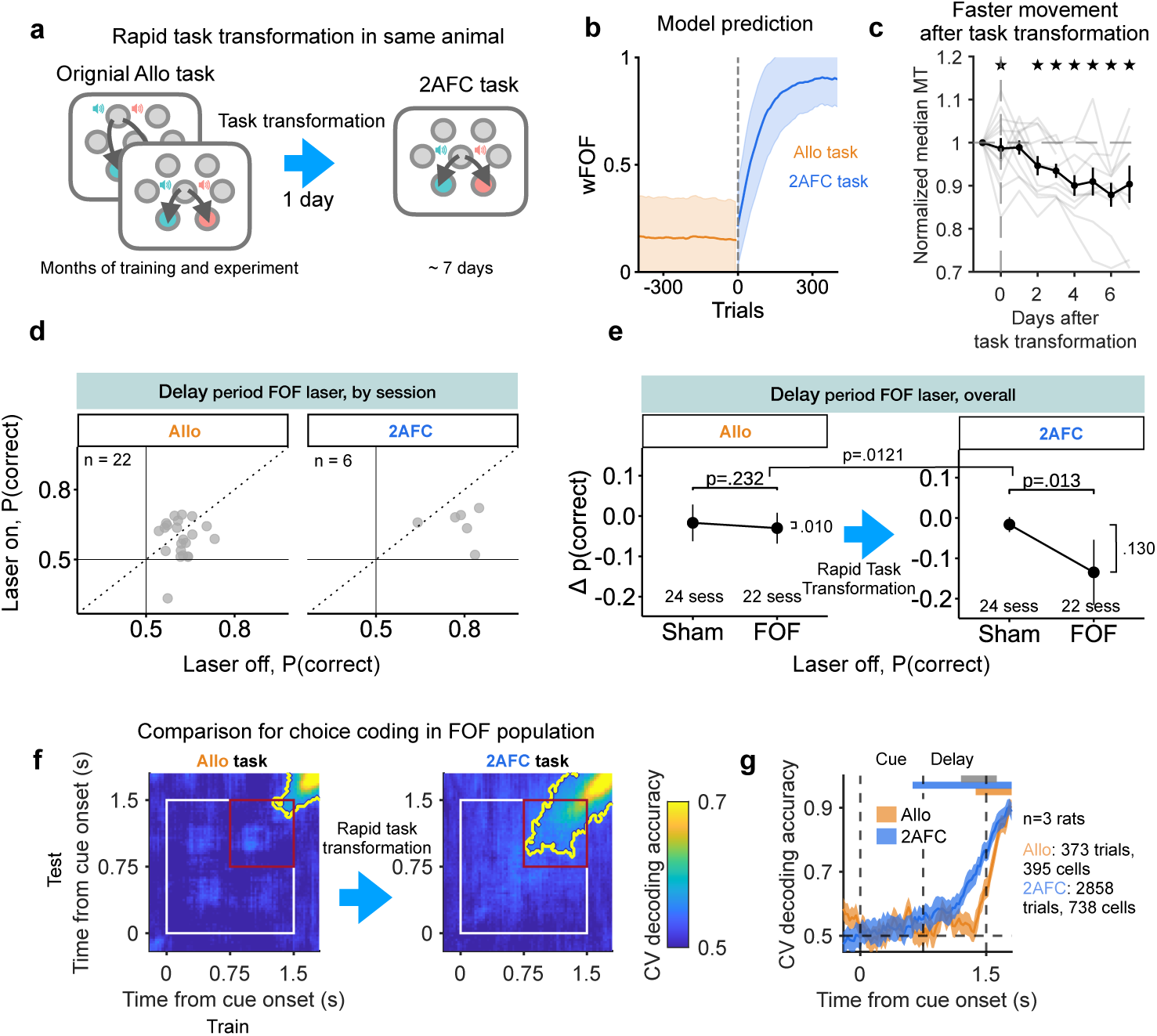
Rats shift to an FOF dependent strategy after task transformation. **a.** We took expert Allo rats and transformed the task into a 2AFC task by removing all start positions except for center-start trials. **b.** Time course of *w*_FOF_ after the Allo-to-2AFC transformation. The solid line and shaded area represent the mean *w*_FOF_ and standard deviation (SD) of *w*_FOF_ across 800 simulated trials. **c.** The thin lines show each rat’s median MT from center-start trials on each day (normalized to the baseline MT for each subject), aligned to the day of task transformation. We plotted the pooled median MT from the 7 days preceding the transformation at day=-1. The black line shows the mean ± SEM across rats. The stars mark days with a significant MT reduction relative to baseline (LRT, *p <* .01). **d.** Performance modulation for the delay period laser effect. Each point represents a single session one FOF session is one point. **e.** Comparison of delay period silencing between Sham and FOF sessions. Y axis: fraction of correct choices for laser-on-trials - laser-off-trials. Dot and error bar reflect mean and 95% bootstrapped CI of the mean across sessions. For comparison, the Allo-task results shown in panels **d,e** (left) are restricted to animals that underwent task transformation (compare with Fig. 4c,d, right). **f.** Across-time cross-validated choice decoding accuracy averaged across 100 pseudopopulations for the Allo (left) and 2AFC (right) tasks. The decoders were trained at one time window (x-axis) and tested at another (y-axis). Contours, *p <* .01 (extreme pixel-based test). The white and red box highlight the fixation and delay period, respectively. Compare with Fig. 3d. **g.** Cross-validated decoding accuracy for choice, across the trial, averaged over 100 pseudo populations. Shade is 99% CI. The orange and blue thick bars at the top indicate above-chance accuracy for the Allo and 2AFC task. Grey bar indicates significant difference between Allo vs. 2AFC tasks (*p <* .05, permutation test with Bonferroni correction for number of time points). Compare with Fig. 3b.

We then compared the causal role of FOF in the Allo task with the 2AFC task, in the same subjects. We followed the same experimental and analysis methods used to compare the causal contribution of FOF to the Ego and Allo tasks Fig. 4, but now for Allo vs. 2AFC task. In the 2AFC task, we observed a significant impairment (Fig. 7d,e-right). Directly comparing the impairment between tasks, we found that silencing the FOF in the 2AFC task resulted in a significantly larger impairment than in the Allo task (*p* = .0121, LRT, Fig. 7e). This confirmed our model prediction: within 2-6 days after simplifying the task, strategies shifted to be FOF dependent. Detailed statistical results can be found in Table S5 and the statistical appendix.

We then compared choice decoding performance before and after task simplification for center-start trials. Within days of task transformation, the decoding of upcoming choice emerged earlier in the trial compared to the Allo task (Fig. 7f). A significant time window of stable choice coding emerged after task transformation (Fig. 7f-right). During the second half of the delay period, decoding accuracy in the reduced 2AFC task was significantly higher than the original Allo task (Fig. 7g, thick grey bar for *p* < .01, from permutation test). While coding in the 2AFC task was weaker than the coding in the Ego task (Fig. 3b), we note that this activity appeared within days of reducing the task from Allo to 2AFC. Together, the optogenetic and electrophysiological experiments support our model predictions and provide evidence for cost-sensitive rapid switching of working memory strategies.

## Discussion

Here, we have shown that, by selecting *when* to perform a sensorimotor transformation, animals flexibly choose the format and locus of working-memory (WM). The conditions under which frontal motor-planning is deployed are set by relative costs of alternative strategies: in Ego where the costs of motor planning are low, animals adopt early sensorimotor transformation and prospective memory in frontal cortex; when costs of motor planning are high, they defer transformation and rely on a retrospective sensory trace instead. Extending a well-established framework, our findings reveal that the reliable appearance of frontal WM in classic delayed-response tasks likely reflects the structure of those tasks—subjects are head-fixed or fixating and must execute brief, precise movements like licks, gaze shifts or reaches—rather than a universal property of how memory is organized. Our results open a path toward a circuit-level mechanistic understanding of the diversity of WM strategies and the flexible control of WM format.

The present study demonstrates that the frontal orienting fields (FOF) play a crucial role in maintaining prospective memory for the Ego task but not the Allo task. In the Ego task, FOF activity during the delay period was essential for performance, maintaining motor plans in self-centered coordinates (Hypothesis 1 in Fig. 1). Strikingly, silencing FOF during the delay did not impair Allo task performance, and delay-period activity was observed in auditory cortex (AC), suggesting that the animals relied on a sensory form of working memory (Hypothesis 3 in Fig. 1), an outcome we had not anticipated from the prior literature. A cost-sensitive model of working memory allocation accurately recapitulated these behavioral and neural changes across tasks (Fig. 6,Fig. S13) and, crucially, made a causal prediction. When we simplified the Allo task, both the neural engagement of FOF and the behavioral response speed shifted to the pattern observed in the Ego task, validating the model’s prediction that animals flexibly allocate mnemonic resources based on computational cost. Together, these findings indicate that FOF supports planning in self-centered coordinates and that rats can rapidly switch strategies when task demands change.

Although our analyses focused on choice-related delay-period activity, we also observed representations of start position, a world-centered variable, in the FOF. Startposition coding in FOF was notably stronger in the Allo task, where world-centered reference frames are behaviorally relevant. In the Allo context, the target ports are in world coordinates, so animals must subtract their start position from the target port associated with the sound cue to determine the correct movement plan, which may explain the stronger representation of positional information. Despite the presence of positional information in FOF, animals in the Allo tasks did not use a FOF-dependent location based working memory strategy (Hypothesis 2 in Fig. 1), suggesting that the FOF cannot maintain a world-centered goal in working-memory, but instead world-centered coding reflects the animals current position^15^. The source of the positional signal remains unknown, but FOF is strongly interconnected with posterior parietal cortex (PPC) and retrosplenial cortex (RSC), regions previously implicated in spatial navigation and position-based computations^62,63^. In our task, animals quickly move to the reward port after indicating their choice. For these kinds of movement sequences, position can could provide an internal error signal through efference copy^56^. Future studies should examine how world- and self-centered representations coexist and interact across these regions to support flexible spatial behavior.

Our findings speak to broader questions about where and how working memory is maintained. In delayed-response tasks, persistent delay activity has been consistently observed in frontal regions^1–5,64^. Frontal cortex therefore appears to be part of a canonical circuit for working memory, especially when animals plan future actions. There are exceptions to this rule: in mice, the locus of short-term memory shifts between frontal and posterior cortex depending on which behavioral strategy the animal adopts^14,18,65,66^. Our results extend this picture in two important respects. First, we provide an experimentally controlled mechanism for inducing mnemonic strategies rather than unexplained individual differences. Second, by manipulating that cost directly (simplifying the Allo task), we show that the shift in WM locus is driven by task structure rather than being a fixed consequence of training curriculum^67,68^. The mechanisms of working memory differ strikingly between FOF in the Ego task and AC in the Allo task. In the Ego task frontal cortex encodes a self-centered, prospective plan, whereas in the Allo task AC encodes a retrospective sensory trace. We also saw that in AC, there are two distinct subspaces: representing sensory *input* during the cue period and sensory *memory* during the delay period (Fig. 5h). In the FOF motor plans ramped up starting in the cue period and occupied a relatively consistent subspace. This motor planning space was not perfectly stable: decoders trained at the beginning of the delay (*t* = 0.75 s) generalized up until the go cue (*t* = 1.5 s) and then decayed (Fig. 3d). This is consistent with previous literature: both sensory and motor WM have been shown to occupy distinct subspaces from there respective inputs or outputs^30,50,69^. Notably, stability of sensory vs. motor memory subspaces differ sharply. Frontal working memory is robust, showing ramping dynamics consistent with bistable attractor dynamics (Fig. 6d)^7,9,70^. In contrast, sensory traces in AC are decaying (Fig. 5)^38^, consistent with a single attractor. This dissociation mirrors findings in human studies, where sensory working memory is more vulnerable to distraction than action-linked memory^71,72^. Our results offer a potential neural account of why action-related WM is ‘prioritized’^73,74^: motor-planning codes are inherently more stable. This helps explain why animals normally rely on a frontal, action-oriented strategy in delayed-response tasks, switching to a sensory code only when the cost of motor planning becomes high.

Given that sensory memory in AC decays before the go cue, the question arises: how do animals in the Allo task maintain their choice? The most parsimonious explanation is that information is transferred to FOF or another downstream structure, such as striatum^28,75^. Consistent with this view, in the Allo task we observed choice-selective activity appearing at the end of the delay in FOF, suggesting late transfer of a stored representation from AC (Fig. S13. This strategy is facilitated by the fixed timing of our task: there was always 1.5 s from the start of fixation until the go cue. To search more broadly for the WM substrate, we recorded from several candidate areas: hippocampus^39,40^ and retrosplenial cortex^41,42^ (RSC) as potential loci of a location-based strategy, and medial prefrontal cortex (mPFC)^43–46^ and superior colliculus (SC)^47^ as potential loci of abstract non-spatial short-term memory. None of these areas showed robust choice-selective delay activity in the Allo task (Fig. S9), though we caution that the hippocampus and RSC datasets are limited in size. Other possibilities cannot be excluded: unrecorded areas may maintain the memory through persistent or activity-silent mechanisms^76,77^, and the absence of choice-selective delay activity does not rule out contributions via non-persistent coding schemes.

Our evidence for a role of AC in the Allo task is correlational. AC carried a retrospective sensory trace whose strength was associated with Allo performance, but we did not perturb AC. We therefore cannot say whether this trace is necessary for bridging the delay, redundantly represented elsewhere, or epiphenomenal. A causal double dissociation would require delay-specific silencing of AC in both tasks, with careful validation of the spatial extent of silencing, which is technically challenging in the rat because AC spans a large cortical volume. Such experiments are an important direction for future work.

Simplifying the Allo task shifted working memory engagement back to frontal cortex, resulting in faster responses, confirming a key prediction of the cost-sensitive model. In the model, cost is abstract. The two tasks differ in several ways, and future work will be required to disentangle the precise nature of this cost. One hypothesis is that it is related to the precision of working memory. There are more possible (and potentially more precise) movements required in Allo vs. Ego. A second hypothesis is that Allo is more difficult since it requires combining two kinds of information on each trial (start position and sound cue). One way to distinguish these hypotheses is to compare version of Ego and Allo with fixation but no working memory requirement. If WM cost is driving the differences in neural mechanism, then removing the WM requirement should remove the pressure to use different strategies in the two tasks, but if the second hypothesis is correct then removing WM requirements should not.

These findings resonate with theories of resource-rational cognition^55,78–80^ and metacognitive control^81^, which propose that animals and humans allocate limited cognitive resources to maximize performance efficiency. In our model, cost sensitivity controls WM format through selective excitation of neural populations. The cost of performing each task using different strategies could be learned by trial and error^82^. One likely candidate for the “controller” is the anterior cingulate cortex (ACC), which has been implicated in performance monitoring and cost–benefit computation^83,84^. In particular, ACC has been shown to learn the cognitive effort associated with certain contexts or actions^85^, but frontal-parietal networks have also been linked to WM effort^86,87^. Altering our paradigm to encourage within-session task switches could reveal neural correlates of cost evaluation and strategy selection at finer temporal resolution and allow for circuit-dissection of these processes.

In conclusion, our results reveal a new principle: working-memory format is dynamically selected according to representational cost and supported by distinct sensory-frontal circuits. By choosing *when* to perform the sensorimotor transformation, animals route memory through whichever circuit best balances precision and cost: frontal motor planning when the mapping is simple and stable, and sensory maintenance when it is not. FOF supports memory when planning in self-centered coordinates is efficient, and rats can flexibly engage alternative strategies when environmental demands shift. This flexibility reflects a form of resource-rational cognition, where animals learn multiple strategies and select the most efficient WM format for a given context. These results open new directions for investigating the neural basis of metacognition, adaptive control, and the dynamic allocation of working memory resources across distributed cortical circuits.

**Supplementary Figure 1.**
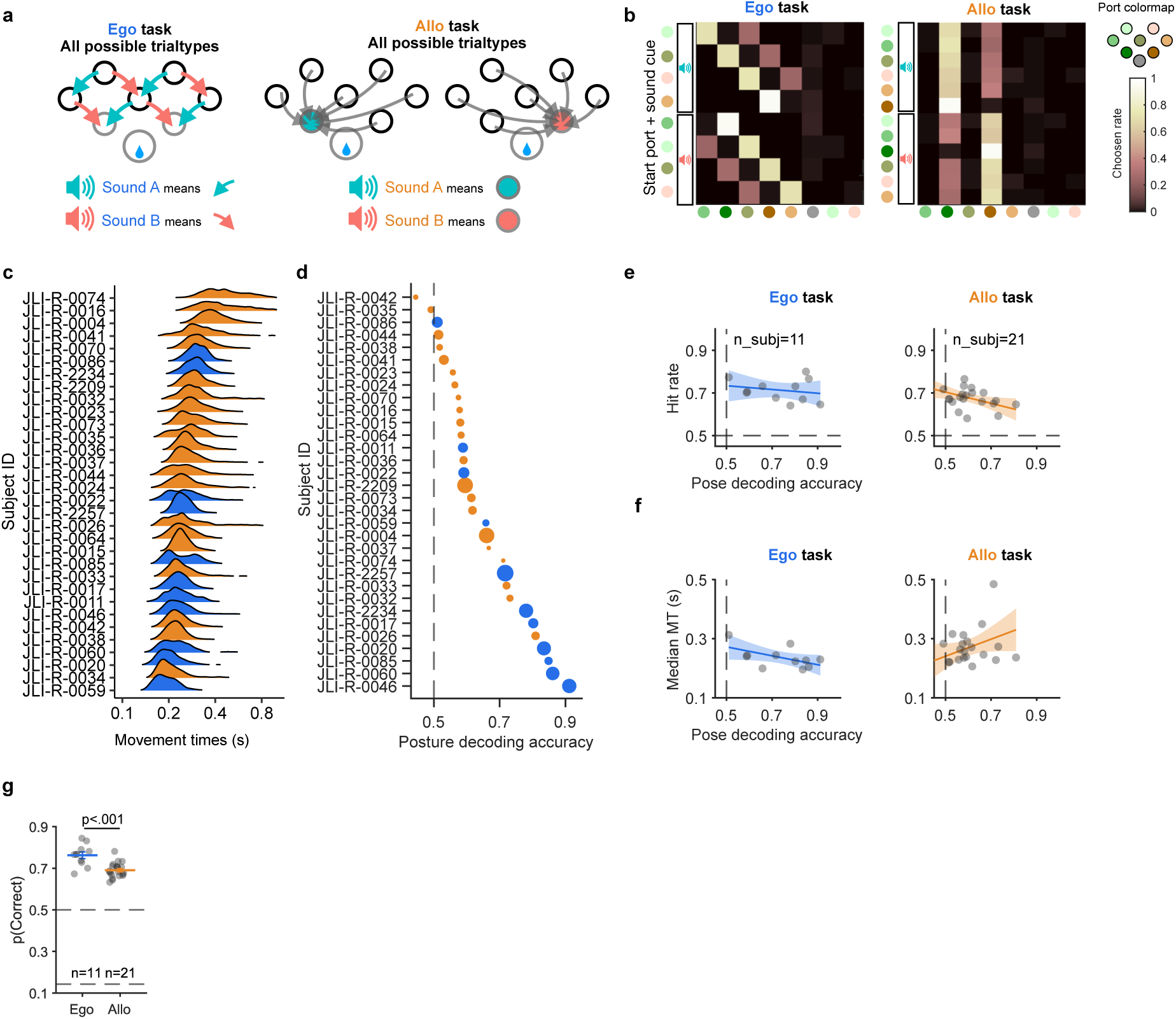
Task and behavioral details. **a.** Trial types for each task. The Ego task had 5 different start ports with 8 different start-port auditory cue combinations. There were only two possible movement vectors in the Ego task: down to the left or right. The Allo task had 7 different start ports with 12 different start-port auditory cue combinations. **b.** Choice contingency table for each trial-type for the Ego task (Left) and the Allo task (Right). Rows: trial-type, sort by start-position (colored dots) and auditory cue (colored speaker), columns: chosen port. Color indicates the proportion of choices to the given port in each trial-type. The colors in each row sum to 1. Note that there are very few out-of-task errors: choices are either correct or to the port that would have been indicated by the unplayed sound. **c.** Distribution of movement times (MT) for each individual rat for center-start trials. MT is defined as the time from center port withdrawal to choice port entry, hit trials only. Orange: Allo animals; Blue: Ego animals. Allo animals are generally slower than Ego animals. **d.** Decoding accuracy for choice from postural features (at *t* = 1.3 s after cue onset) for each individual rat (colored dot) for center-start trials. Dot size indicates the number of trials. Decoding accuracy was generally higher for the subjects in the Ego task. **e.** Scatter plot of overall hit rate vs. pose decoding accuracy on center-start trials, for Ego task (Left) or Allo task (Right). Lines are TLS fit. Shades are for 95% CI. There is no evidence that task performance improves with a more overt postural strategy. **f.** Scatter plot of MT vs. posture decoding accuracy on center-start trials. Each dot is a subject, for Ego task (Left) or Allo task (Right). Lines are TLS fit. Shades are for 95% CI. The correlation between MT and overt postural strategies across subjects is weak. **g.** Performance. Grey dots are individual rats and the line indicates the mean for each task.

**Supplementary Figure 2.**
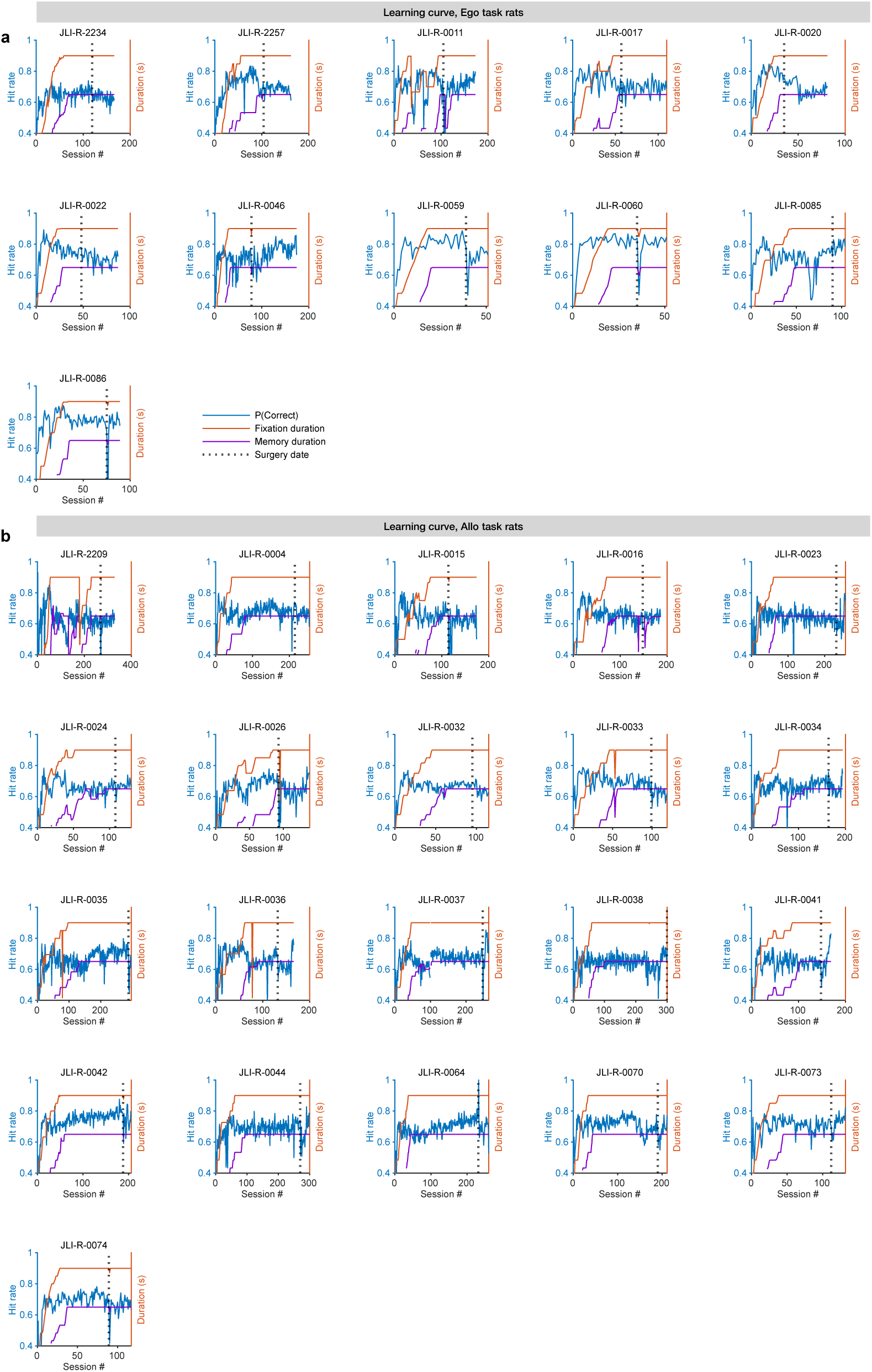
Learning curves over training. **a.** Learning curve for Ego task animals. Correct rate and timing (fixation duration and memory period duration) development over the number of sessions. **b.** Learning curve for Allo task animals. As in a.

**Supplementary Figure 3.**
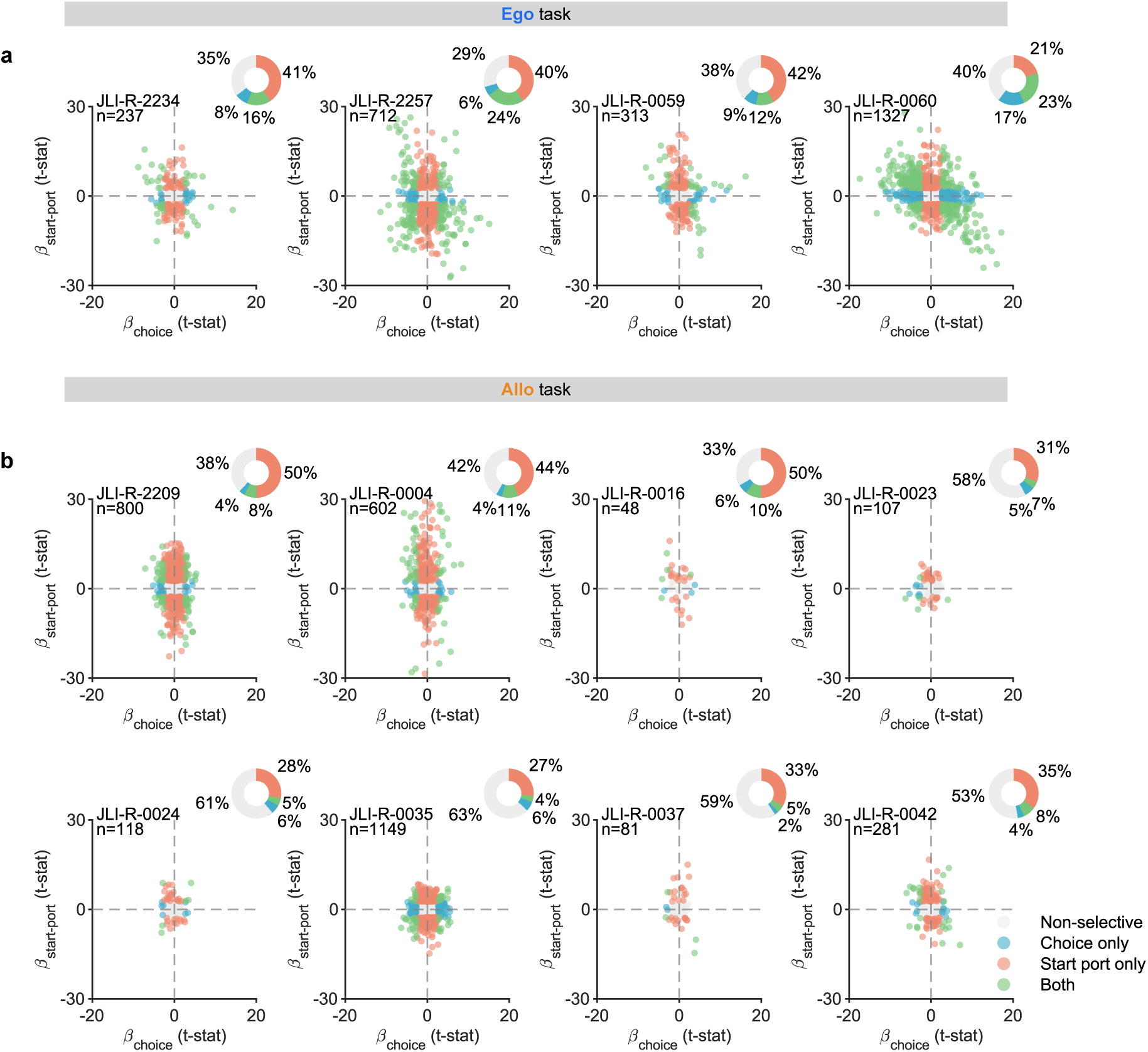
Distribution of single-neuron selectivity indices across subjects. **a.** Scatter plot showing the distribution of coefficients of the start port against choice from the mixed-effects linear models for individual neurons recorded from each rat performing the Ego task. Each dot represents a single neuron, with its position reflecting selectivity for upcoming choice (x-axis) and start port position (y-axis), corresponding to analyses in Fig. 2d. **b.** Same as (a), but for the Allo task, corresponding to Fig. 2h. Pie plots next to each panel summarize the proportions of neurons classified as non-selective (gray), choice-selective only (blue), start port-selective only (orange), or selective for both (purple) at the *p <* .01 level.

**Supplementary Figure 4.**
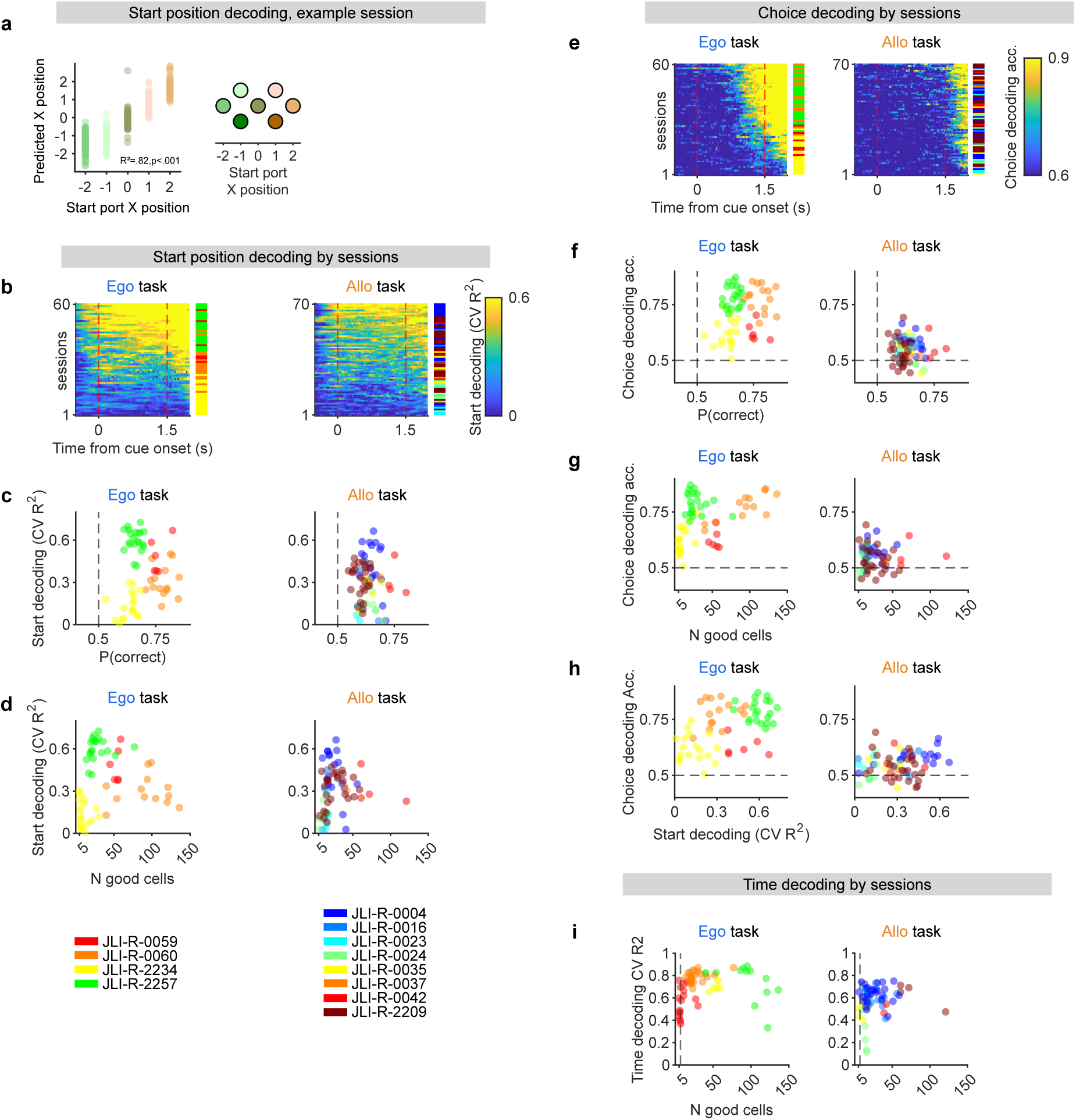
Session-wise decoding performance for start position, choice, and time across subjects. **a.** Example session of start port position decoding. Each dot indicates a trial, performance was quantified as *R*^2^ between actual and predicted X position. **b-i.** The map from subjects to colors of points is shown in the lower left corner of the figure. For all panels, Ego task results are shown on the left and the Allo task results on the right. **b.** Heatmaps of decoding performance for start position in the Ego (left panel) and Allo (right panel) task. Each row represents a session, with color reflecting decoding performance (cross-validated (CV) *R*^2^). Sessions are ordered based on the center of mass of decoding performance (calculated as the dot product between time and decoding performance). The color bar adjacent to each heatmap indicates the subject identity for each session, consistent with the color legend shown below. **c.** Scatter plots showing the relationship between start decoding *R*^2^ and behavioral performance for each session. Each point represents a session, color-coded by subject (Ego-left, Allo-right). **d.** Decoding performance as a function of the number of good cells recorded per session, as determined by quality metrics (see *Methods*). **e.** Like **b** but for choice decoding on center-start trials, with the color indicating the CV decoding accuracy. Note, that choice decoding in the Allo task emerges after the go cue. **f.** Like **c** but for choice decoding. **g.** Like **d** but for choice decoding. **h.** Relationship between choice and start position decoding performance across sessions. The dashed lines represent chance-level performance thresholds (50% for choice decoding). **i.** Time decoding performance (CV *R*^2^) against the number of good cells for Ego and Allo tasks. Each point in the scatter plots corresponds to an individual session, color-coded by subject. The dashed lines represents the inclusion threshold based on cell count for a comparison in Fig. 6j.

**Supplementary Figure 5.**
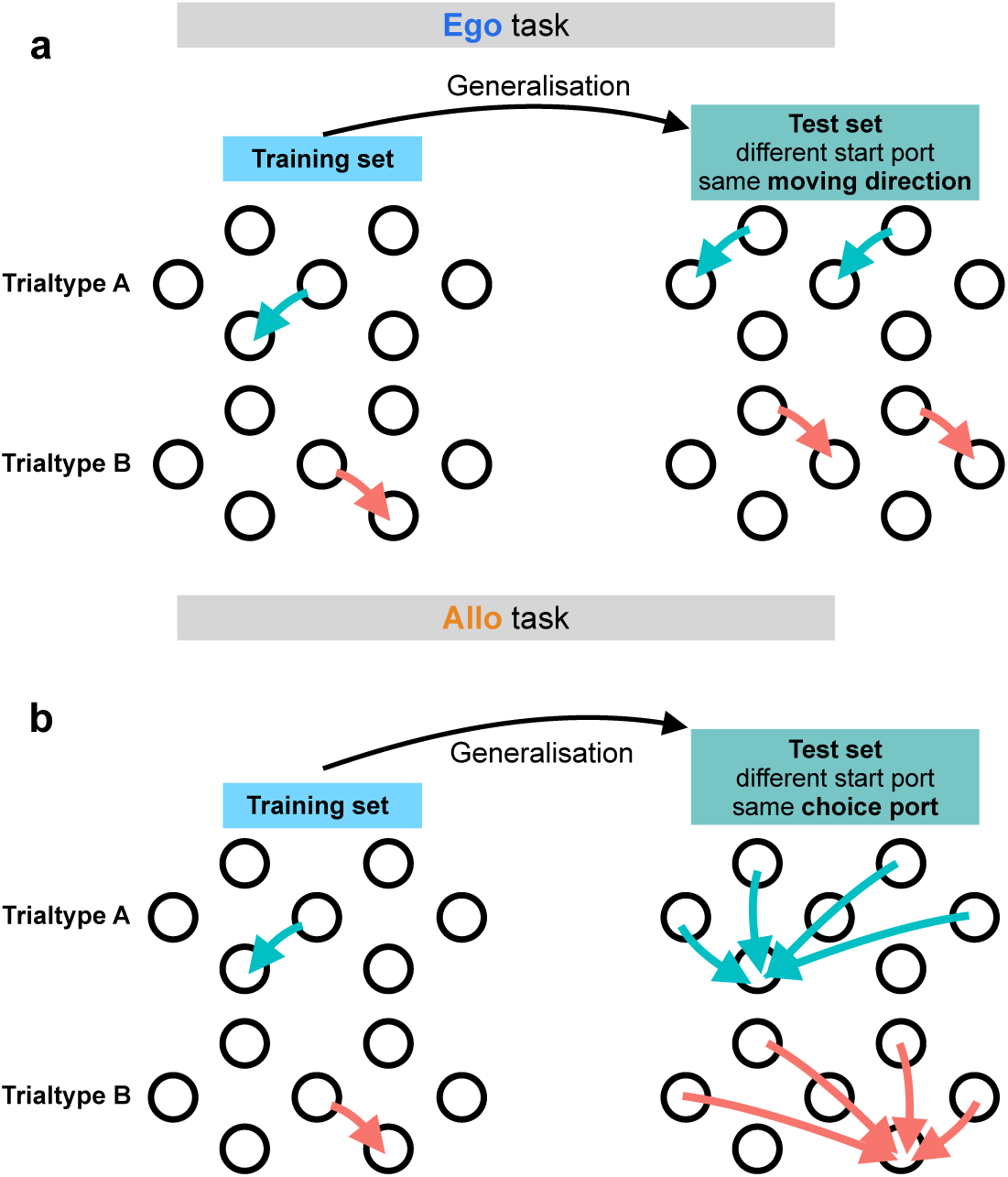
Trial type details for decoding generalization analysis **a.** for the Ego task, center-start trials were selected for the training set (Left) and other trials started from other ports but moved to the same direction were selected as test set (Right). Decoding results were shown in Fig. 3k and Fig. 3m. **b.** for the Allo task, center-start trials were selected for the training set (Left), trials started from other ports but moved to the same choice port were selected as test set (Right), decoding results were shown in Fig. 3l and Fig. 3n.

**Supplementary Figure 6.**
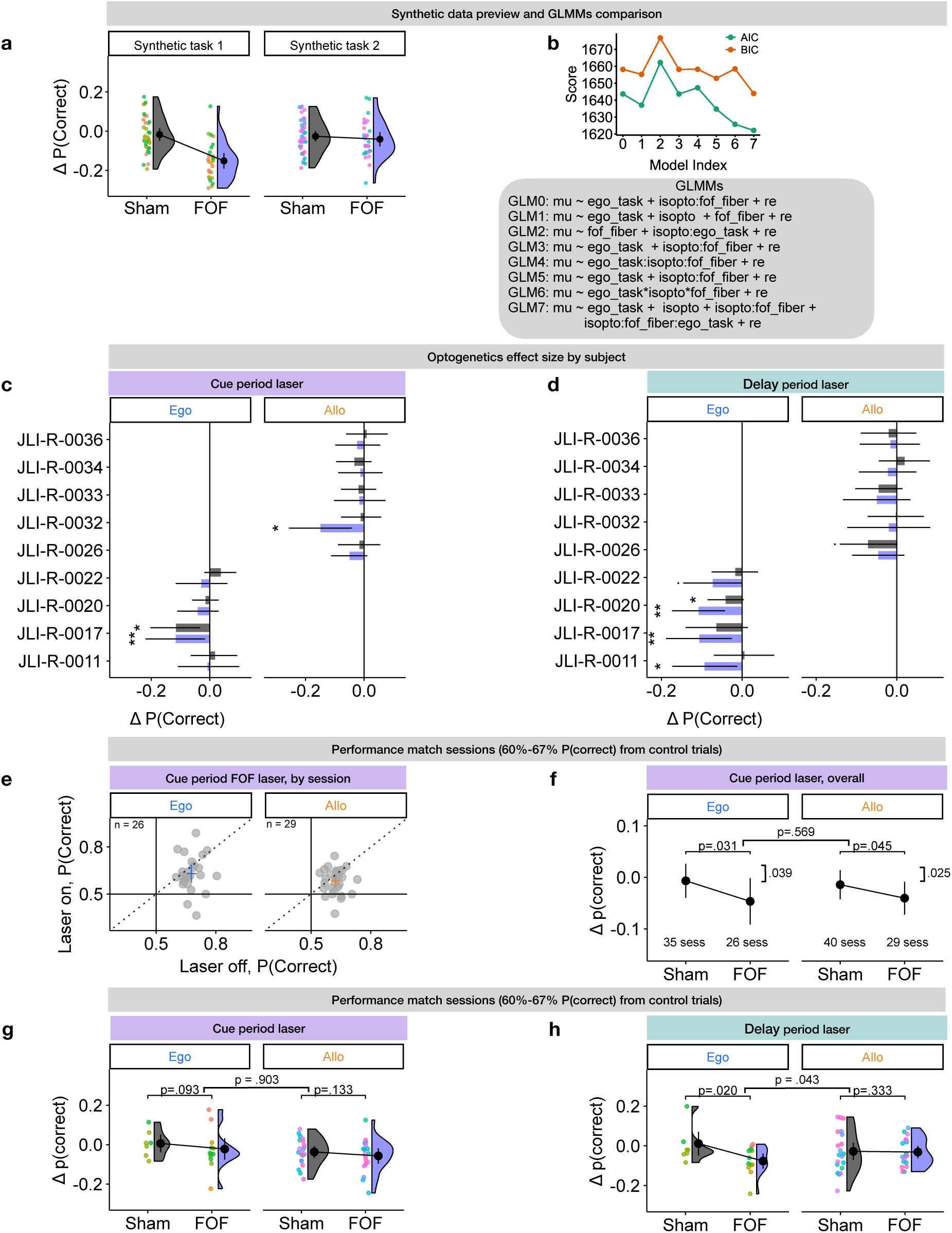
**a.** Synthetic data that simulated optogenetic modulation for some animals and sessions across two tasks. Y axis: fraction of correct choices for laser on trials - laser off trials for each session (dots, colored by different animals). Shade: density estimation with Gaussian kernel, black dot: mean over all sessions, error bar: 95% CI (bootstrap). **b.** Performance of 8 potential GLMM models on the synthetic data. The best model, GLM7, was used for the main results. **c.** Performance modulation for the cue period laser effect for each animal. Y-axis: individual animals, X-axis: change in performance, (laser-on trials) - (laser-off trials), averaged across sessions for each subject. Error bars are 95% CI (bootstrapped over sessions). Purple: FOF sessions; Grey: Sham sessions. **d.** As in **c**, but for the delay period. **e.** As in Fig. 4c, but for the cue period. Each FOF session is one point; Crosses reflect mean and 99% bootstrapped CI of the mean across sessions. **f.** As in Fig. 4d, but for the cue period. Dot and error bar reflect mean and 95% bootstrapped CI of the mean across sessions. **g,h.** As in panel **f** and Fig. 4d, respectively, but restricting analyses to sessions with 60-67% performance on laser-off trials. Dots are sessions colored by different animals. Violins show kernel density estimation.

**Supplementary Figure 7.**
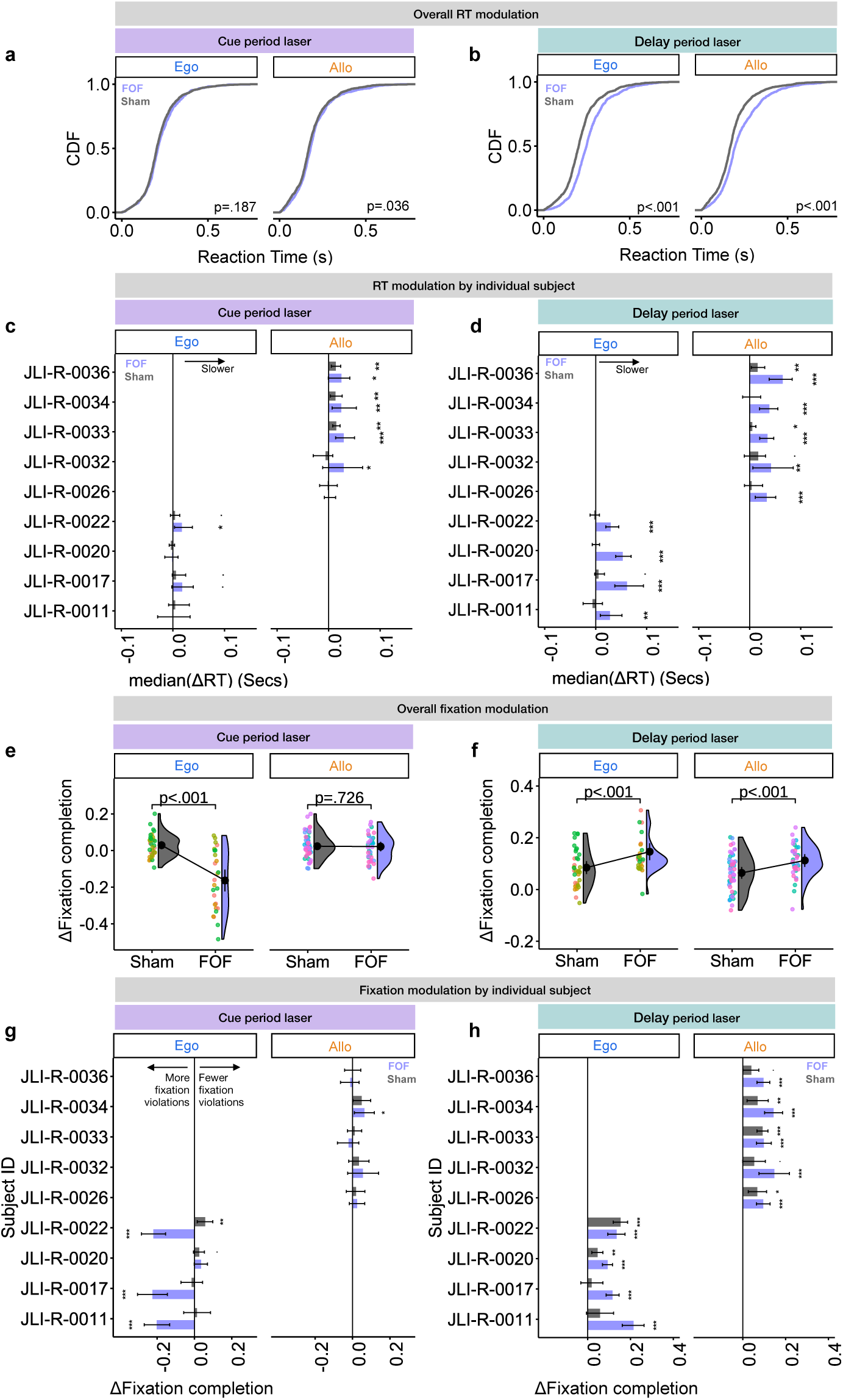
Influence of FOF silencing on reaction times (RT) and fixation violations. RT was defined as the time from go cue (at 1.5 s after the auditory cue onset) to withdrawal from the start port. A fixation violation was defined as exiting from the start port before the go cue. We quantified fixation completion rate as the fraction of trials without fixation violations. **a.** The distributions of RT for cue period laser-ON trials. The grey and purple lines indicated the cumulative distribution function (CDF) sham fiber trials and FOF fiber trials, respectively. p-values were from bootstrap. **b.** Same as **a** but for the delay period. Silencing FOF during the delay significantly slowed RT, compared to sham trials. **c.** Influence of cue period laser on RT for each animal. Y-axis: individual animals, X-axis: median of ΔRT across sessions for (laser-on trials - laser-off trials). Error bars are 95% CI (bootstrapped across sessions). Purple: FOF sessions; Grey: Sham sessions. **d.** Same as **c** but for the delay period. **e.** Effects of cue period laser on fixation. Y axis: change in fixation completion rate for (laser-on trials - laser-off trials) for each session (dots, colored by different animals). **f.** Same as **e** but for delay period laser effect. **g.** Effect of cue period laser on fixation for each animal. Y-axis: individual animals, X-axis: Change in fixation completion rate for (laser-on trials - laser-off trials) for each session. Error bars are 95% CI (bootstrapped across sessions). Purple: FOF sessions; Grey: Sham sessions. **h.** Same as **g** but for the delay period.

**Supplementary Figure 8.**
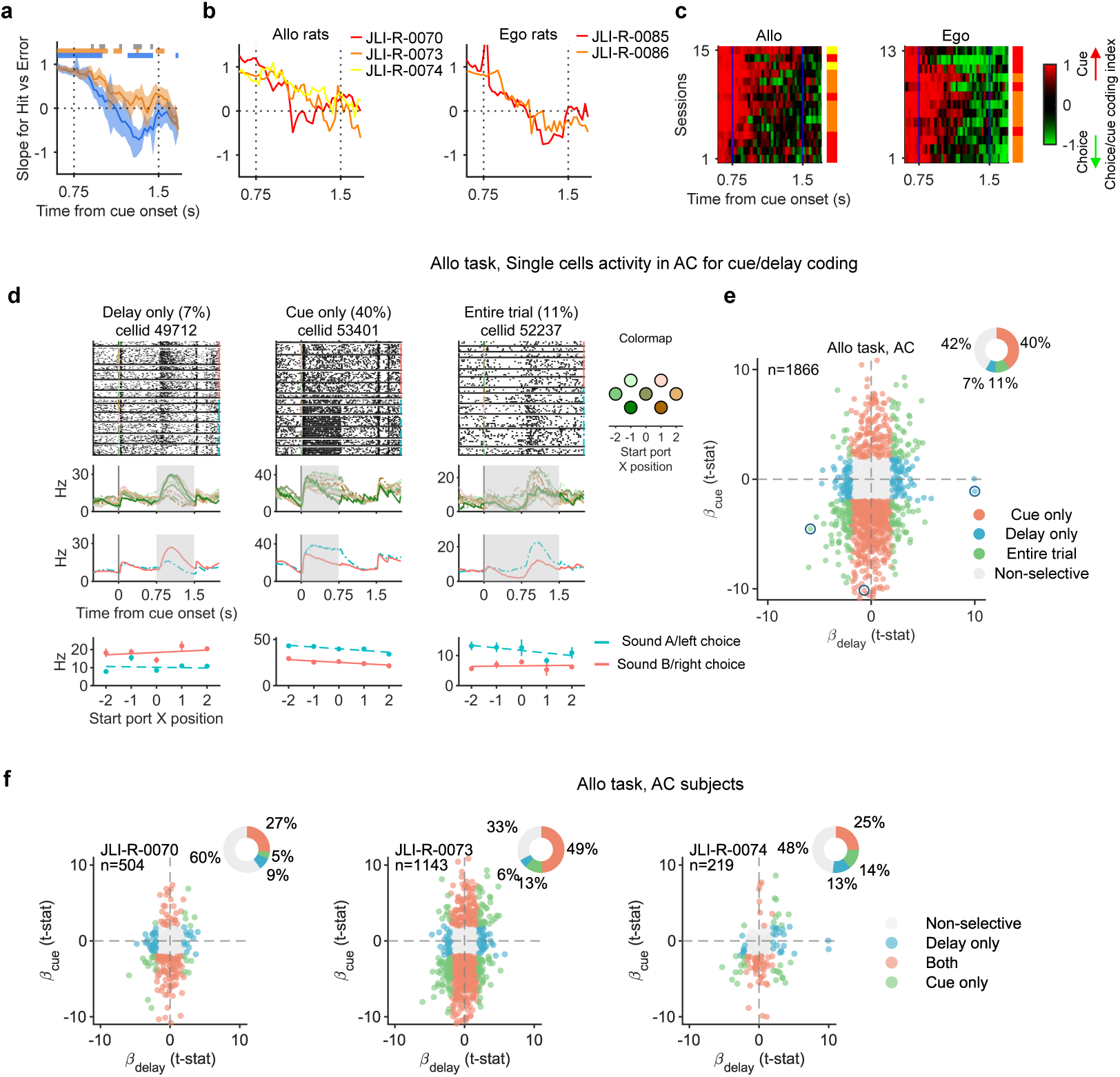
Detailed results for AC recording in the Allo and Ego task. **a.** Time course of the total least-squares regression slope between AC cue-coding coefficients on correct and error trials, shown separately for the Allo and Ego tasks. At each time point, only neurons with significant cue coding on correct trials were included. Positive slopes indicate cue coding, whereas negative slopes indicate choice coding. Lines show slopes and shaded areas show bootstrapped 95% CI. Colored bars indicate slopes significantly different from zero for each task, and gray bars indicate significant differences between tasks (p < .05, FDR corrected). **b.** Same analysis as a, separated by each rat from the Allo (left) or Ego (right) task. **c.** Cue/choice coding index for individual Allo and Ego recording sessions, corresponding to the population summary in Fig. 5c. Each row represents one session. Positive values indicate cue coding and negative values indicate choice coding. Sessions were sorted by mean correct-trial decoding accuracy during the delay period. Colored bars beside the heatmaps indicate individual rats as in panel b. **d.** Example neurons from the AC recorded in the Allo task. Spike raster plots and PSTHs of three representative neurons show different dynamics for cue coding in the delay period (Left), cue period (Middle), or both (Right). Notation is the same as Fig. 2e-g. **e.** Distribution of coefficients of single AC neuron cue coding for cue period and delay period spike count. Example neurons from (a) are circled. **f.** Scatter plot showing the distribution of coefficients of the sound coding in the cue period against the delay period from the mixed-effects linear models for individual neurons recorded from each rat performing the Allo task. Corresponding to analyses in Fig. 5b.

**Supplementary Figure 9.**
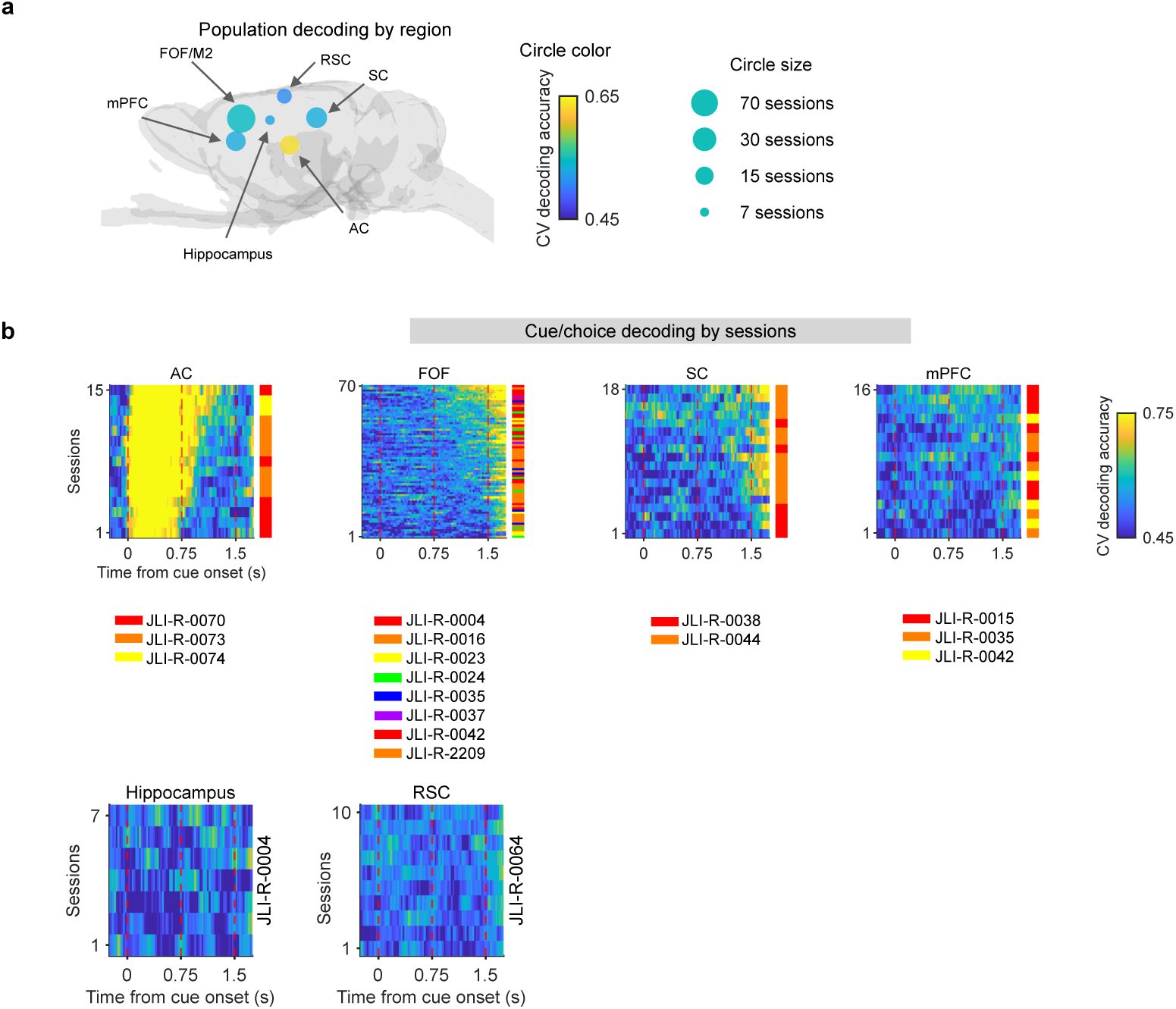
Detailed results for AC, FOF, SC, and mPFC recording experiment. **a.** Session-wise average decoding accuracy at 0.75 to 1.25s after cue onset from each recording region. Dot color reflecting averaged decoding accuracy, size reflecting number of sessions. **b.** Heatmaps showing session-wise decoding performance for choice on correct trials in the Allo task from AC, FOF, SC, mPFC, Hippocampus and RSC sessions. Each row represents a session, with blue-yellow color reflecting cross-validated decoding accuracy. Sessions are ordered based on the center of mass of decoding performance (calculated as the dot product between time and decoding performance). The color bar to the right of each heatmap indicates the subject identity for each session, consistent with the color legend shown below each panel.

**Supplementary Figure 10.**
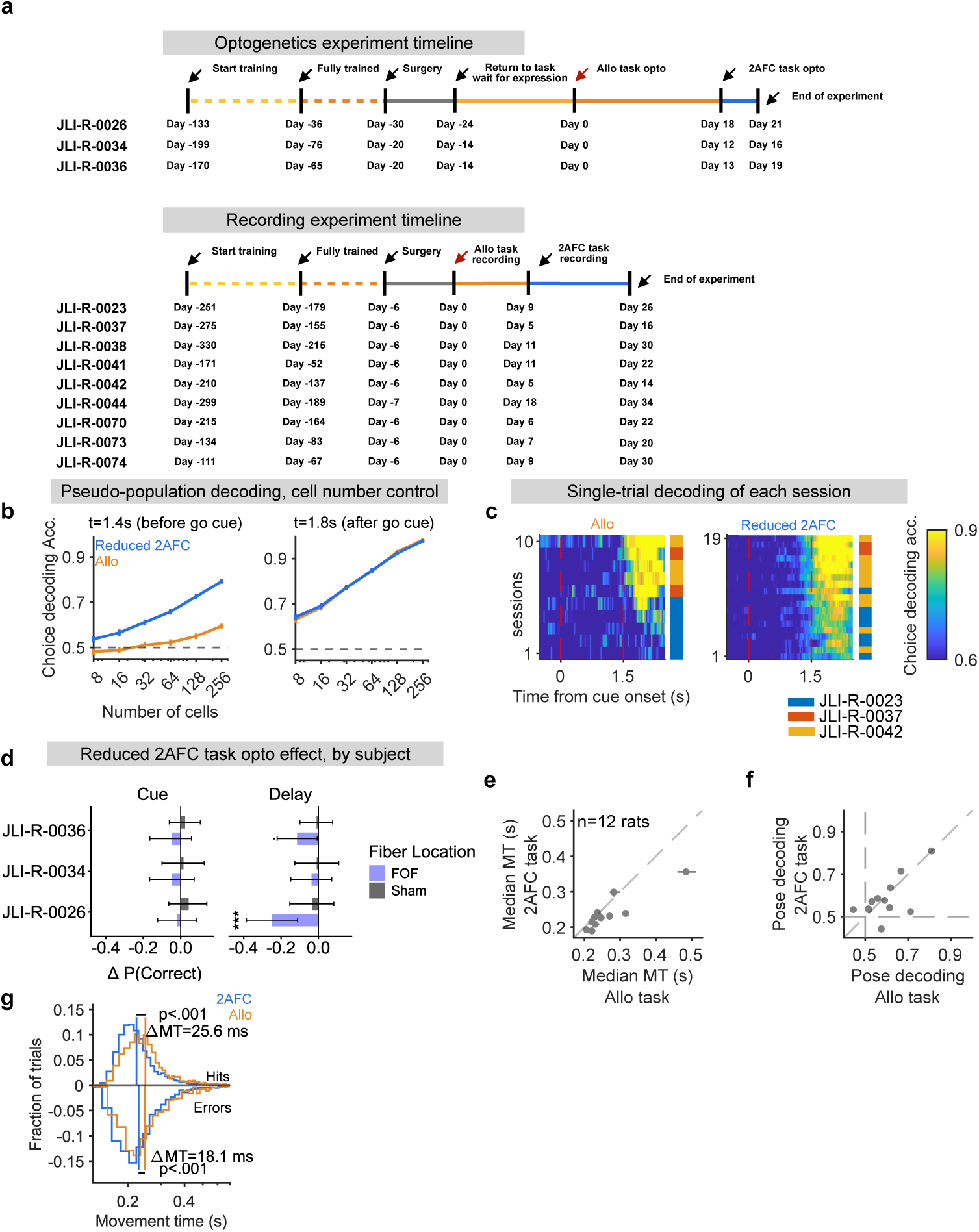
Details for task transformation experiments. **a.** Experimental timeline for each subject, showing the sequence of sessions during the Allo and reduced 2AFC tasks with optogenetic manipulations and electrophysiological recording experiments. Each row represents an individual subject. **b.** Pseudo-population decoding accuracy for choice as a function of the number of cells in the pseudo-population. Decoding was performed at two key time points, during the delay period (t = 1.4 s from cue onset) and during the movement period (t = 1.8 s from cue onset). Error bars indicate SEM over pseudopopulations. Orange line represents data from animals in the Allo task. Blue line is from the same animals after the task transformation to 2AFC. The horizontal dashed line represents chance-level decoding. **c.** Session-wise heatmaps of choice decoding accuracy from electrophysiological data for the Allo (left) and reduced 2AFC (right) task. Each row represents a session, and color intensity reflects cross-validated decoding accuracy over time, aligned to cue onset. Sessions are sorted by the center of mass of decoding accuracy. The color bar to the right indicates the subject identity for each session, consistent with the color legend shown below. **d.** Optogenetic silencing effects on task performance (Δ*P* (correct)) for individual subjects during the cue (left) and delay period (right), for the reduced 2AFC task. Each row represents a subject, and error bars indicate 95% CI estimated from bootstrap resampling. **e.** Comparison of movement times (MT) on center-start trials for each subject before and after task transformation from the Allo to the reduced 2AFC task. Each dot and error bar represents the median and 95%CI of the median (bootstrapped across trials) for each each subject. **f.** Comparison of posture decoding accuracy at t=1.3s after cue onset (during late-delay period) for each subject before and after task transformation from the Allo to the reduced 2AFC task on center-start trials. Each dot represents an individual subject. The horizontal and vertical dashed lines represent chance-level decoding thresholds. **g.** Movement time (MT) distribution for correct (Top) and error (Bottom) trials in the Allo (orange) and 2AFC (blue) tasks. Vertical lines: medians of MT.

**Supplementary Figure 11.**
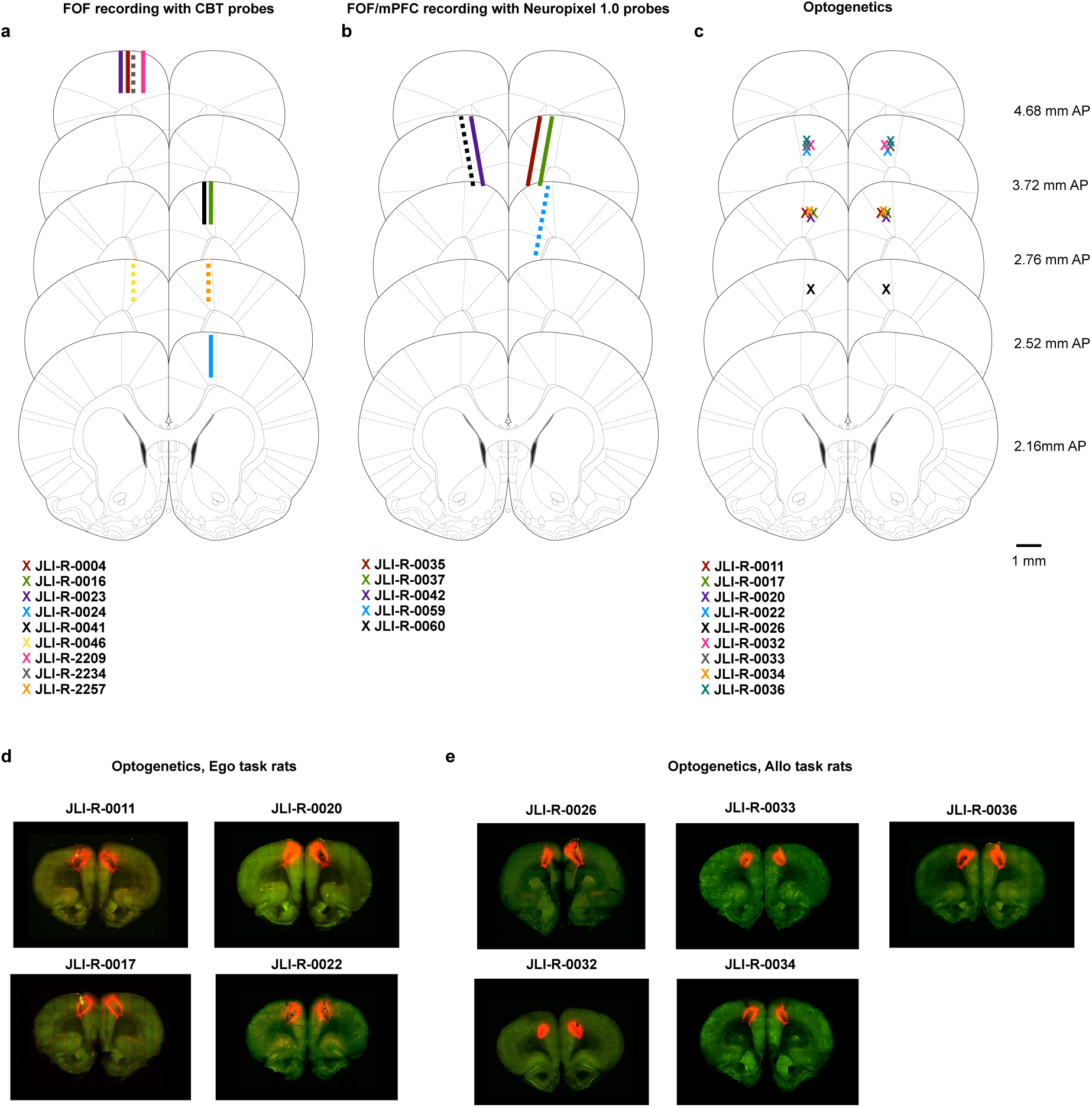
Histology results for FOF experiment. Note, that in the nomenclature of Paxinos and Watson ^88^ the area that we describe as the FOF is considered to be part of M2. **a.** Reconstructed probe track for each rat that implanted Cambridge Neurotech silicon probes in FOF. Solid line for the Allo task, dash line for the Ego task. **b.** Reconstructed probe track for each rat that implanted Neuropixel 1.0 probes in FOF. Solid line for the Allo task, dash line for the Ego task. **c.** Reconstructed optic fiber track for each rat with optogenetics surgery in bilateral FOF. **d.** Histology image for each rat from optogenetics experiment, in the Ego task. **e.** Histology image for each rat from optogenetics experiment, in the Allo task.

**Supplementary Figure 12.**
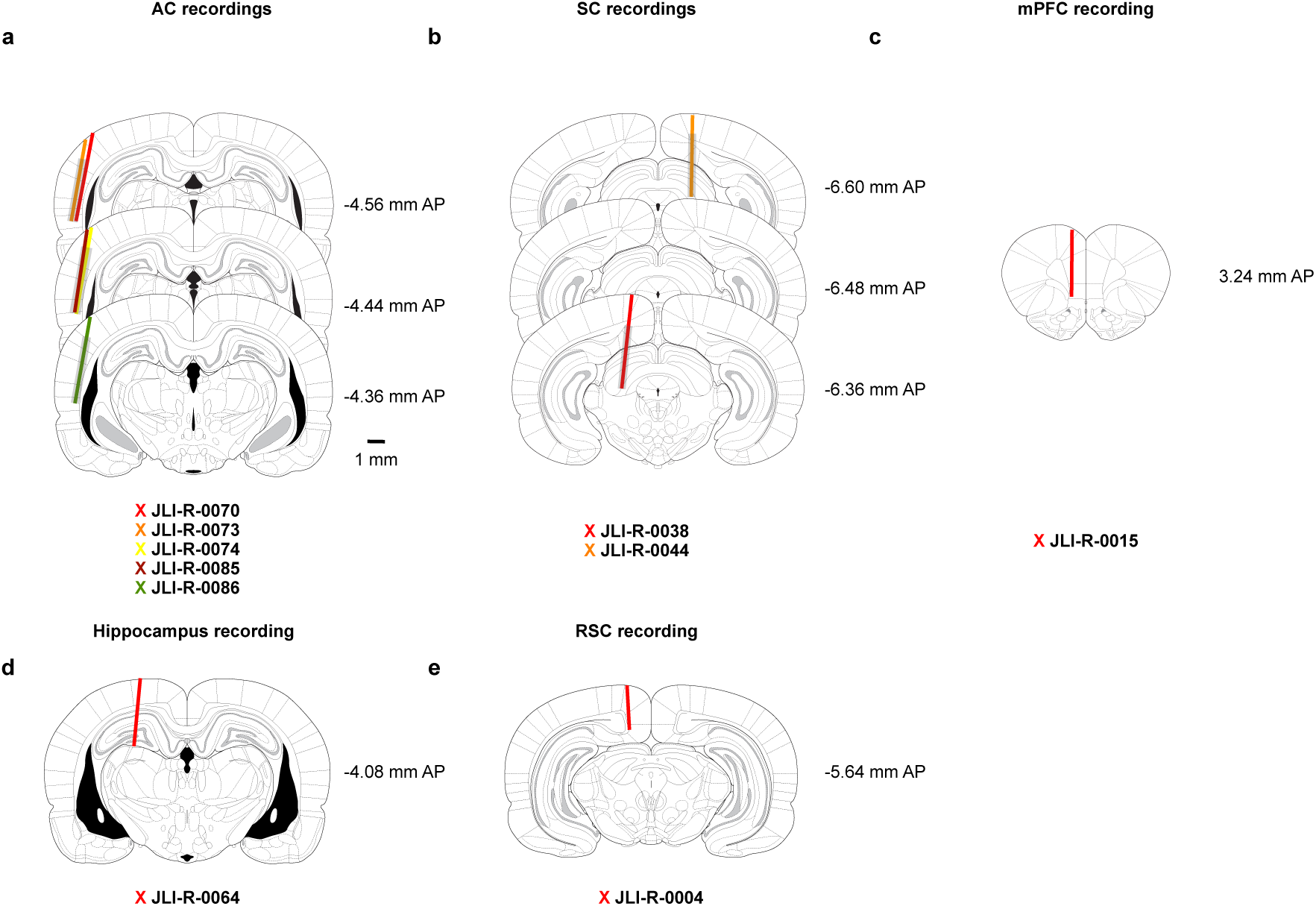
Histology results for AC/SC/mPFC recordings. **a.** Reconstructed probe track for each AC recording rat. The grey bar indicates the channel selection positions for bank0. **b.** Reconstructed probe track for each SC recording rat. The grey bar indicates the channel selection positions for bank0. **c.** Reconstructed probe track for one single mPFC implanted rat with Cambridge Neurotech probe. Other rats that have both FOF and mPFC recordings using Neuropixel probes are shown in Fig. S11b) **d.** Reconstructed probe track for the Hippocampus recording rat. **e.** Reconstructed probe track for the RSC recording rat.

### A shared AC–FOF network model of working memory

The simple model in the main text illustrates the central idea of flexible working-memory organisation. As a supplementary extension, we developed a more detailed AC–FOF network to reproduce the observed task-dependent dynamics across both regions. Changing only one parameter, the gain of external excitation to FOF, we recapitulated key experimental findings across the Ego and Allo tasks: distinct sensory encoding and memory representations in AC, choice-related AC delay activity in Ego versus cue-related activity in Allo, and earlier FOF choice signals in Ego than Allo.

The model consisted of two auditory sensory nodes, two separate auditory memory nodes^50,89^, and two competing FOF nodes (Fig. S13a). The AC sensory nodes received the auditory cue and provided input to FOF. Sensory activity was loaded into the AC memory nodes during a brief window after cue offset (Fig. S13c). Both the AC memory and FOF node pairs used recurrent excitation and mutual inhibition, with different parameters. Therefore, AC memory exhibited bistable dynamics subject to leak and noise, whereas increasing external excitation could drive FOF into a bistable regime with ramping selective activity. FOF provided feedback to the AC memory nodes in both tasks. All network parameters and inputs were shared between tasks, except for the gain of external excitation to FOF. We simulated left or right choices from the centre port, without simulating other start positions or movement vectors. Parameters were not fitted to the recording data.

In the simulations, FOF developed strong choice-planning activity early in the Ego task, whereas choice signals emerged only late and remained weaker in Allo (Fig. S13d,e). This task-dependent timing parallels the single-neuron choice selectivity and population choice decoding observed in FOF recordings (Fig. 3a,b).

AC showed strong sensory responses in both tasks, but its delay-period activity differed (Fig. S13f,g). On error trials, AC memory nodes reversed their cue preference in Ego, following the model choice, but retained their cue preference in Allo (Fig. S13h), resembling the example AC neurons in Fig. 5a. Applying the same cue/choice coding index based on correct and error trials as in the main paper confirmed choice-related delay activity in Ego and cue-related delay activity in Allo (Fig. S13i), consistent with the recordings (Fig. 5d). In the model, the choice-related AC activity in Ego reflected stronger top-down input from FOF, consistent with non-sensory signals reported in auditory cortex projection neurons during behavior^90^.

Together, the model captured these task-dependent FOF and AC dynamics by changing only the gain of the external excitation ramp to FOF.

#### Simulation and state variables

We simulated 2,000 trials per task with equal numbers of the two auditory cues, using Euler integration with the parameters in Table 1. Time *t* is measured in seconds relative to cue onset, with integration step Δ*t* = 0.01 s. We denote the cue by *c* ∊ {1, +1}, with +1 indicating the left-choice cue and 1 the right-choice cue. The AC sensory states *S*_1_*, S*_2_ and memory states *M*_1_*, M*_2_ were deviations from baseline; *r_L_, r_R_* denote FOF activities. Sensory and memory states were initialized from zero-mean Gaussian distributions with standard deviations *σ_S_*/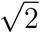 and *σ_M_* /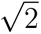, respectively; FOF states were initialised at zero. Initial conditions, cue sequences and noise samples were identical across tasks.

**Supplementary Figure 13.**
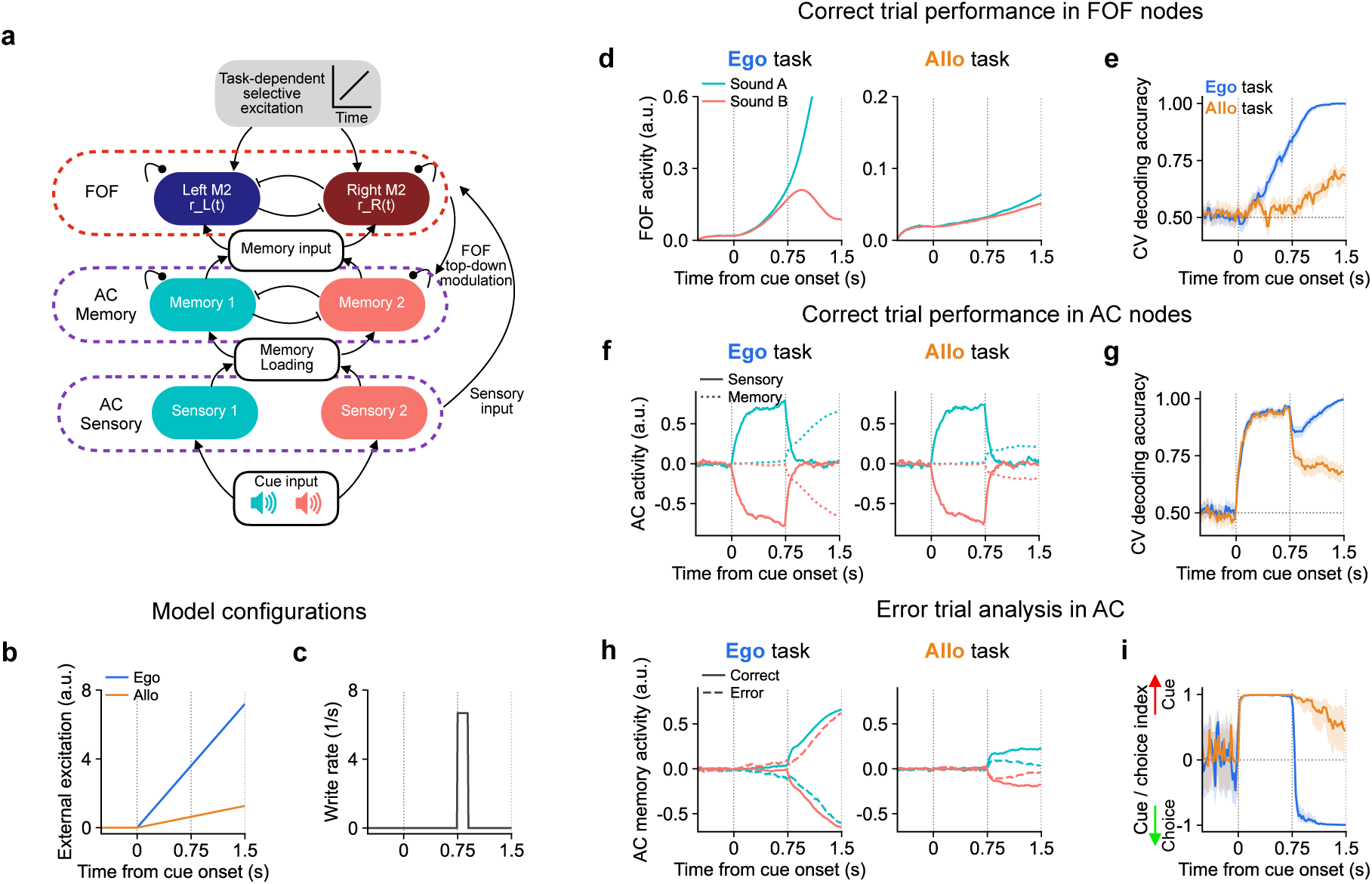
A shared AC–FOF circuit with task-dependent external excitation. **a,** Model schematic. Circle-ended connections denote excitation and bar-ended connections denote inhibition; boxes and arrows indicate information transfer between node groups. **b,** External excitation to FOF, the only task-dependent input or parameter. **c,** Shared sensory-to-memory write rate, *κ*(*t*), during the brief loading window after cue offset. **d,** Mean activity grouped by cue in an example FOF node for the Ego (left) and Allo (right) tasks. **e,** Cross-validated decoding of choice for both tasks from model FOF activity. **f,** Mean activity grouped by cue in example AC sensory (solid) and memory (dotted) nodes for the Ego (left) and Allo (right) tasks. **g,** Cross-validated decoding of cue for both tasks from model AC activity. **h,** Mean activity grouped by cue in an example AC memory node for the Ego (left) and Allo (right) tasks. Solid and dashed curves denote correct and error trials, respectively. **i,** AC cue/choice coding index for both tasks, as in Fig. 5d. Shading in **e,g,i** indicates 95% CI.

#### Auditory sensory nodes

The cue drove an opponent sensory response in AC along the unit axis *v* = (1*, −*1)/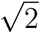. For node *i ∈ {*1, 2*}*, the sensory update was

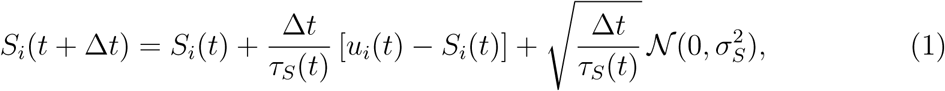

where *u_i_*(*t*) = *cv_i_* during the cue and zero otherwise, and *v_i_* is the corresponding component of *v*. The sensory time constant *τ_S_*(*t*) and noise amplitude *σ_S_* are given in Table 1. Independent samples of Gaussian noise, *N*(0, σ^2^), were added to the nodes at each time step. The Gaussian samples had zero mean and variance *σ*^2^, and were independent across nodes, time steps and trials, and identical across tasks.

#### Separate recurrent AC memory nodes

The AC memory nodes combined leak, recurrent excitation and mutual inhibition. The sensory-to-memory write rate *κ*(*t*) was non-zero only during 0.75 ≤ *t <* 0.90 s (Fig. S13c; Table 1). The binary indicator *ℓ*(*t*) controlled recurrence, which remained active after loading ended:

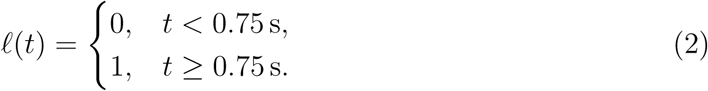

Both recurrent node pairs used the sigmoid response function *F*(*x*) = 1/(1 + exp(*−x*)). The deterministic rate of change *D_i_*(*t*) was calculated using the other memory node *j ≠ i*:

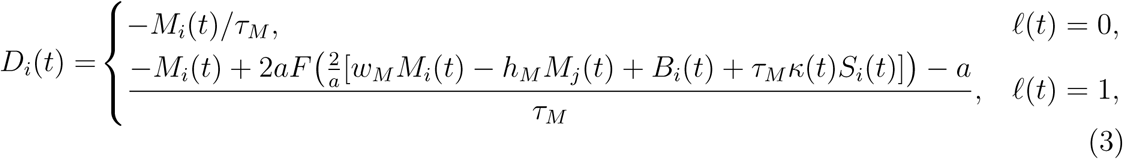

where *B_i_*(*t*) is FOF feedback. The time constant *τ_M_*, recurrent weights *w_M_, h_M_* and nonlinear saturation scale *a* are defined in Table 1. The full update was

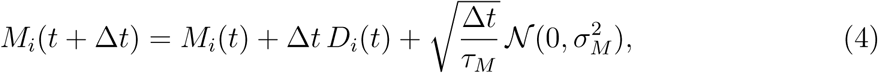

where *σ_M_* is the memory noise amplitude.

#### Sensory and memory input to FOF

We projected the two AC pairs onto their opponent axes,

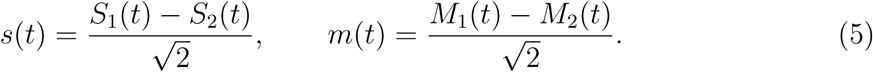

The AC inputs to the left and right FOF nodes were

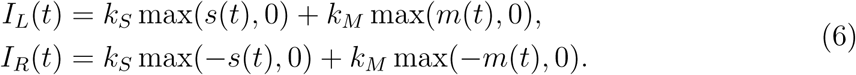

where *k_S_* = 0.05 and *k_M_* = 0.4 are constant sensory and memory input weights, respectively. Both weights were identical across tasks and remained fixed throughout the trial.

#### FOF network and external excitation

We modeled FOF as two mutually inhibiting nodes with recurrent excitation^7,9^. The two nodes represented left and right model choices. For *i* ∈ {*L, R*}, with *j* denoting the other node, their update was

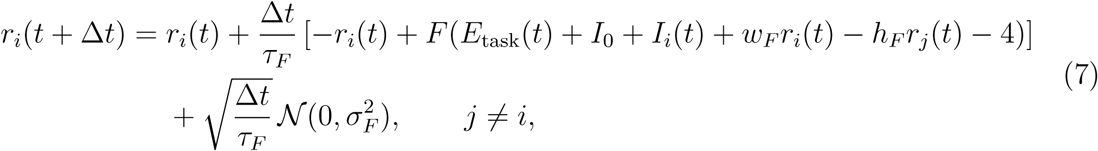

where the time constant *τ_F_*, tonic input *I*_0_, recurrent weights *w_F_, h_F_* and noise amplitude *σ_F_* are defined in Table 1. The external excitation *E*_task_(*t*) was zero before cue onset and increased linearly thereafter, scaled by the task-dependent gain *w*_task_:

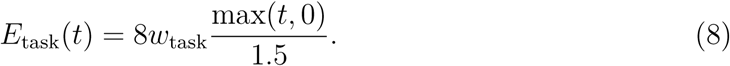

The task-dependent gains were fixed at the values in Table 1; all other network parameters and inputs were shared.

#### Feedback from FOF to AC

FOF feedback, a top-down choice signal to AC^90^, acted on the two memory nodes with opposite signs^91^:

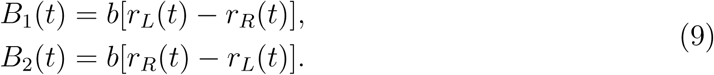

where *b* = 0.8 was constant throughout the trial and identical across tasks. It entered the active memory dynamics after memory loading began, as specified in Eq. 3.

#### Model choices and downstream readout noise

We assigned one model choice per trial from the final FOF state, after the update beginning at 1.49 s. In the state-update equations this is *T* = 1.50 s, corresponding to the saved sample labelled 1.49 s. The decision variable was

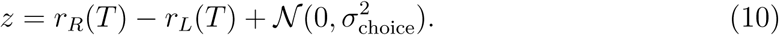

Independent Gaussian readout noise was drawn for each trial, with standard deviation *σ*_choice_ (Table 1). The model chose right (*ĉ* = 1) if *z >* 0 and left (*ĉ* = +1) otherwise, and was correct when *ĉ* = *c*. Readout noise was applied after neural integration^92–94^. The same readout rule and noise amplitude were used in both tasks.

#### Model parameters

Table 1 summarises the parameters used in the simulations. Only the external excitation gain differed between tasks.

#### Decoding from network activity

Decoding was performed at each timepoint with 20 ms intervals, using logistic regression with L2 regularization and a 50/50 split of correct trials for training and testing (Fig. S13e,g). On correct trials, choice corresponded to the cue. Mean activity in panels d,f was also calculated from correct trials.

For the cue/choice coding index, cue classifiers were trained on correct trials and evaluated on cross-validated correct trials (50/50 split) and error trials at 20 ms intervals. The index was calculated using the formula in the main Methods (Fig. 5c,d). 95% confidence intervals were estimated from 1,000 bootstrap resamples.

**Supplementary Table 1.**
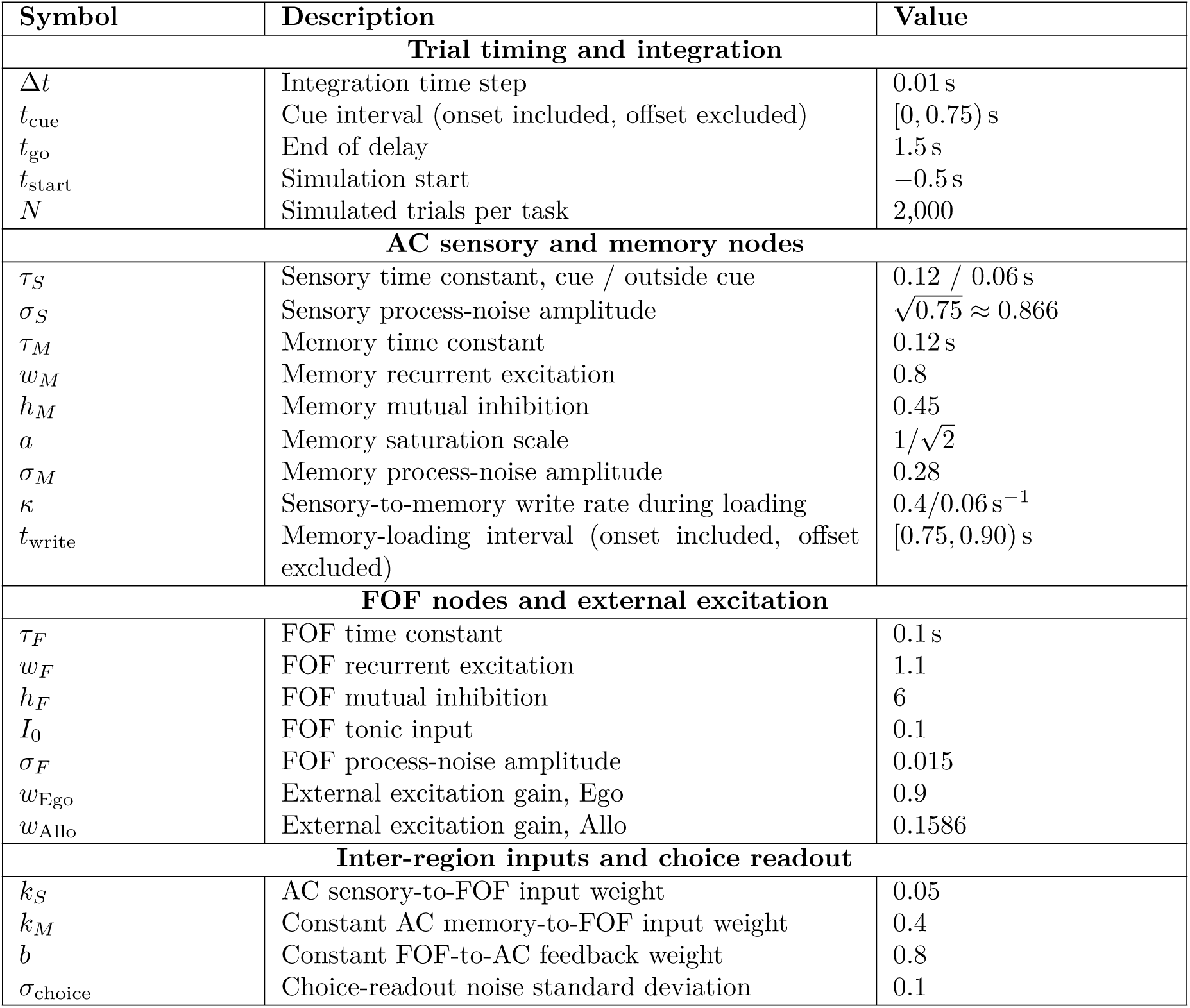
Parameters of the shared AC–FOF network model. Process-noise amplitudes are dimensionless and follow the scaling specified in the state-update equations.

| Symbol | Description | Value |
|---|---|---|
| **Trial timing and integration** |  |  |
| Δt | Integration time step | 0.01 s |
| t<sub>cue</sub> | Cue interval (onset included, offset excluded) | [0, 0.75) s |
| t<sub>go</sub> | End of delay | 1.5 s |
| t<sub>start</sub> | Simulation start | −0.5 s |
| N | Simulated trials per task | 2,000 |
| **AC sensory and memory nodes** |  |  |
| τ<sub>S</sub> | Sensory time constant, cue / outside cue | 0.12 / 0.06 s |
| σ<sub>S</sub> | Sensory process-noise amplitude | √0.75 ≈ 0.866 |
| τ<sub>M</sub> | Memory time constant | 0.12 s |
| w<sub>M</sub> | Memory recurrent excitation | 0.8 |
| h<sub>M</sub> | Memory mutual inhibition | 0.45 |
| a | Memory saturation scale | 1/√2 |
| σ<sub>M</sub> | Memory process-noise amplitude | 0.28 |
| κ | Sensory-to-memory write rate during loading | 0.4/0.06 s<sup>−1</sup> |
| t<sub>write</sub> | Memory-loading interval (onset included, offset excluded) | [0.75, 0.90) s |
| **FOF nodes and external excitation** |  |  |
| τ<sub>F</sub> | FOF time constant | 0.1 s |
| w<sub>F</sub> | FOF recurrent excitation | 1.1 |
| h<sub>F</sub> | FOF mutual inhibition | 6 |
| I<sub>0</sub> | FOF tonic input | 0.1 |
| σ<sub>F</sub> | FOF process-noise amplitude | 0.015 |
| w<sub>Ego</sub> | External excitation gain, Ego | 0.9 |
| w<sub>Allo</sub> | External excitation gain, Allo | 0.1586 |
| **Inter-region inputs and choice readout** |  |  |
| k<sub>S</sub> | AC sensory-to-FOF input weight | 0.05 |
| k<sub>M</sub> | Constant AC memory-to-FOF input weight | 0.4 |
| b | Constant FOF-to-AC feedback weight | 0.8 |
| σ<sub>choice</sub> | Choice-readout noise standard deviation | 0.1 |

## Statistical appendix table for optogenetics experiment

**Supplementary Table 2.**
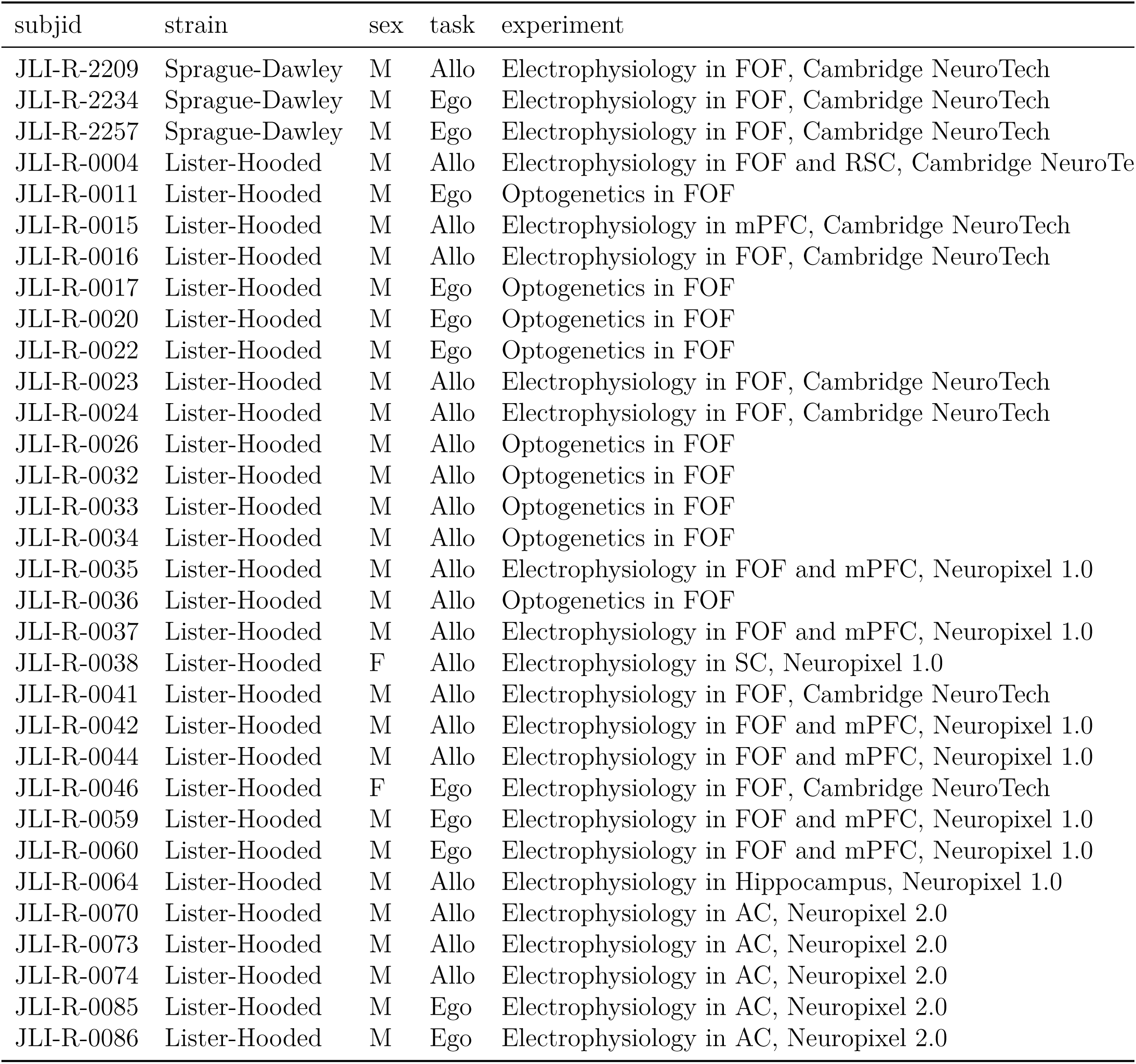
Experiment type for all subjects.

| subjid | strain | sex | task | experiment |
|---|---|---|---|---|
| JLI-R-2209 | Sprague-Dawley | M | Allo | Electrophysiology in FOF, Cambridge NeuroTech |
| JLI-R-2234 | Sprague-Dawley | M | Ego | Electrophysiology in FOF, Cambridge NeuroTech |
| JLI-R-2257 | Sprague-Dawley | M | Ego | Electrophysiology in FOF, Cambridge NeuroTech |
| JLI-R-0004 | Lister-Hooded | M | Allo | Electrophysiology in FOF and RSC, Cambridge NeuroTe[illegible] |
| JLI-R-0011 | Lister-Hooded | M | Ego | Optogenetics in FOF |
| JLI-R-0015 | Lister-Hooded | M | Allo | Electrophysiology in mPFC, Cambridge NeuroTech |
| JLI-R-0016 | Lister-Hooded | M | Allo | Electrophysiology in FOF, Cambridge NeuroTech |
| JLI-R-0017 | Lister-Hooded | M | Ego | Optogenetics in FOF |
| JLI-R-0020 | Lister-Hooded | M | Ego | Optogenetics in FOF |
| JLI-R-0022 | Lister-Hooded | M | Ego | Optogenetics in FOF |
| JLI-R-0023 | Lister-Hooded | M | Allo | Electrophysiology in FOF, Cambridge NeuroTech |
| JLI-R-0024 | Lister-Hooded | M | Allo | Electrophysiology in FOF, Cambridge NeuroTech |
| JLI-R-0026 | Lister-Hooded | M | Allo | Optogenetics in FOF |
| JLI-R-0032 | Lister-Hooded | M | Allo | Optogenetics in FOF |
| JLI-R-0033 | Lister-Hooded | M | Allo | Optogenetics in FOF |
| JLI-R-0034 | Lister-Hooded | M | Allo | Optogenetics in FOF |
| JLI-R-0035 | Lister-Hooded | M | Allo | Electrophysiology in FOF and mPFC, Neuropixel 1.0 |
| JLI-R-0036 | Lister-Hooded | M | Allo | Optogenetics in FOF |
| JLI-R-0037 | Lister-Hooded | M | Allo | Electrophysiology in FOF and mPFC, Neuropixel 1.0 |
| JLI-R-0038 | Lister-Hooded | F | Allo | Electrophysiology in SC, Neuropixel 1.0 |
| JLI-R-0041 | Lister-Hooded | M | Allo | Electrophysiology in FOF, Cambridge NeuroTech |
| JLI-R-0042 | Lister-Hooded | M | Allo | Electrophysiology in FOF and mPFC, Neuropixel 1.0 |
| JLI-R-0044 | Lister-Hooded | M | Allo | Electrophysiology in FOF and mPFC, Neuropixel 1.0 |
| JLI-R-0046 | Lister-Hooded | F | Ego | Electrophysiology in FOF, Cambridge NeuroTech |
| JLI-R-0059 | Lister-Hooded | M | Ego | Electrophysiology in FOF and mPFC, Neuropixel 1.0 |
| JLI-R-0060 | Lister-Hooded | M | Ego | Electrophysiology in FOF and mPFC, Neuropixel 1.0 |
| JLI-R-0064 | Lister-Hooded | M | Allo | Electrophysiology in Hippocampus, Neuropixel 1.0 |
| JLI-R-0070 | Lister-Hooded | M | Allo | Electrophysiology in AC, Neuropixel 2.0 |
| JLI-R-0073 | Lister-Hooded | M | Allo | Electrophysiology in AC, Neuropixel 2.0 |
| JLI-R-0074 | Lister-Hooded | M | Allo | Electrophysiology in AC, Neuropixel 2.0 |
| JLI-R-0085 | Lister-Hooded | M | Ego | Electrophysiology in AC, Neuropixel 2.0 |
| JLI-R-0086 | Lister-Hooded | M | Ego | Electrophysiology in AC, Neuropixel 2.0 |

**Supplementary Table 3.**
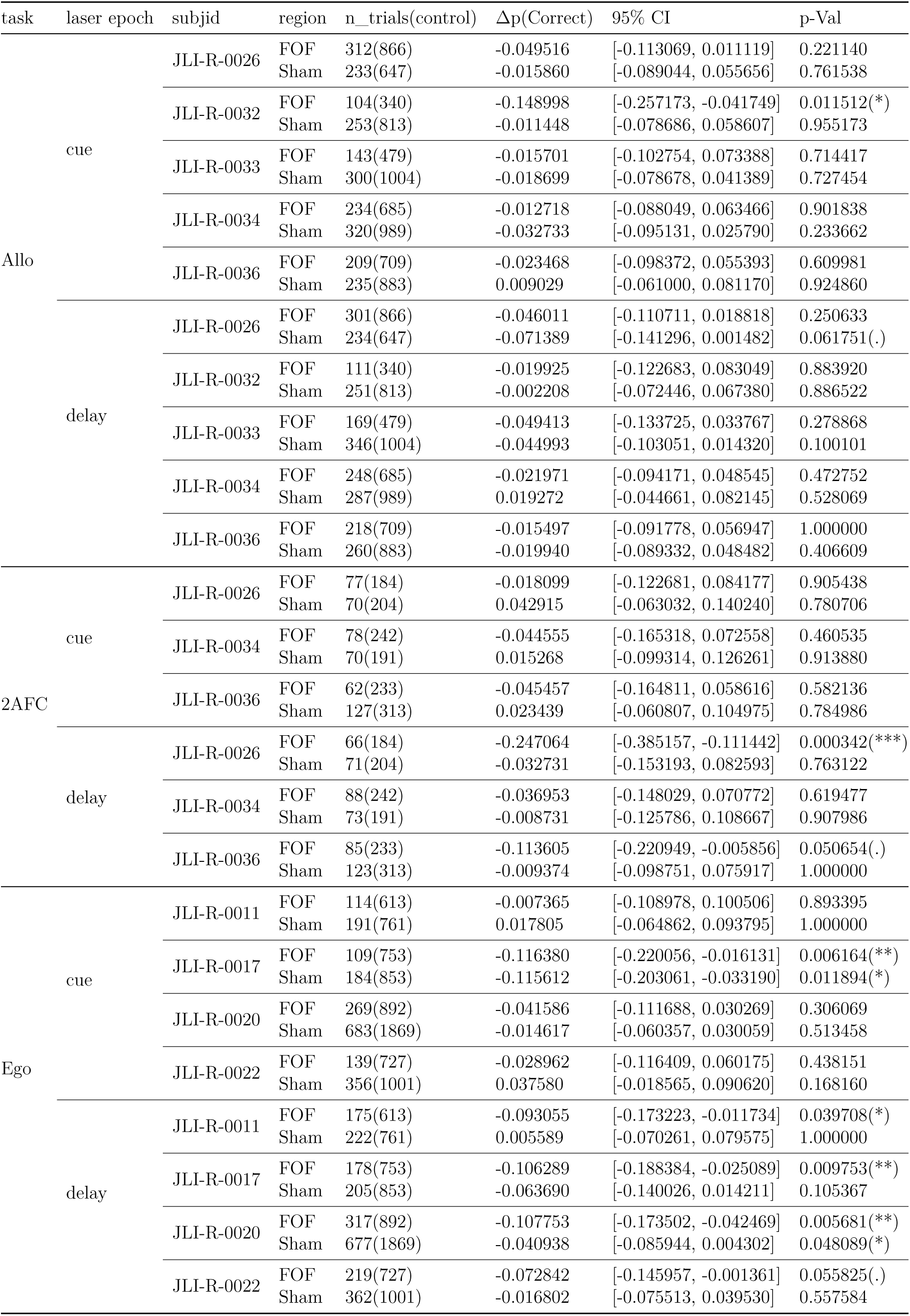
Δp(Correct) and 95% CI between laser on/off trials for each rat on each experiment.

**Supplementary Table 4.**
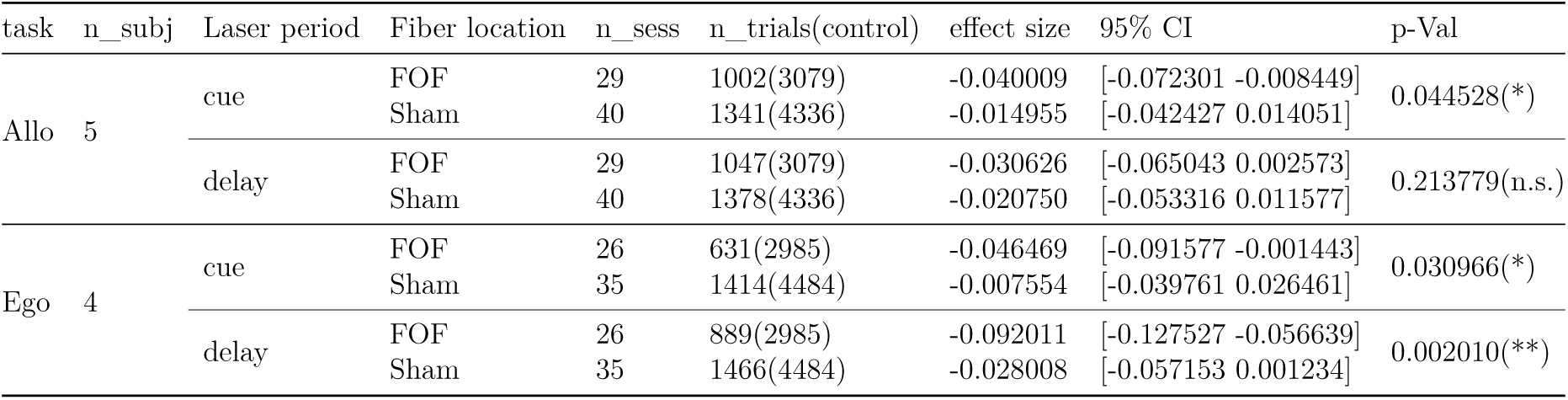
Effect size for Δ*p*(correct) between laser on/off trials in Allo/Ego task.

| task | n_subj | Laser period | Fiber location | n_sess | n_trials(control) | effect size | 95% CI | p-Val |
|---|---|---|---|---|---|---|---|---|
| Allo | 5 | cue | FOF | 29 | 1002(3079) | -0.040009 | [-0.072301 -0.008449] | 0.044528(*) |
|  |  |  | Sham | 40 | 1341(4336) | -0.014955 | [-0.042427 0.014051] |  |
|  |  | delay | FOF | 29 | 1047(3079) | -0.030626 | [-0.065043 0.002573] | 0.213779(n.s.) |
|  |  |  | Sham | 40 | 1378(4336) | -0.020750 | [-0.053316 0.011577] |  |
| Ego | 4 | cue | FOF | 26 | 631(2985) | -0.046469 | [-0.091577 -0.001443] | 0.030966(*) |
|  |  |  | Sham | 35 | 1414(4484) | -0.007554 | [-0.039761 0.026461] |  |
|  |  | delay | FOF | 26 | 889(2985) | -0.092011 | [-0.127527 -0.056639] | 0.002010(**) |
|  |  |  | Sham | 35 | 1466(4484) | -0.028008 | [-0.057153 0.001234] |  |

**Supplementary Table 5.** Effect size for Δ*p*(correct) between laser on/off trials before and after task transformation (n=3 rats)

| task | n_subj | Laser period | Fiber location | n_sess | n_trials(control) | effect size | 95% CI | p-Val |
|---|---|---|---|---|---|---|---|---|
| Allo | 3 | cue | FOF | 22 | 755(2260) | -0.029123 | [-0.061580 0.003018] | 0.202307(n.s.) |
|  |  |  | Sham | 24 | 788(2519) | -0.014743 | [-0.049044 0.020055] |  |
|  |  | delay | FOF | 22 | 767(2260) | -0.029118 | [-0.068147 0.008136] | 0.231717(n.s.) |
|  |  |  | Sham | 24 | 781(2519) | -0.017182 | [-0.062346 0.028509] |  |
| 2AFC | 3 | cue | FOF | 6 | 217(659) | -0.034595 | [-0.064793 0.000592] | 0.150659(n.s.) |
|  |  |  | Sham | 7 | 267(708) | 0.026149 | [-0.050796 0.110651] |  |
|  |  | delay | FOF | 6 | 239(659) | -0.129932 | [-0.215686 -0.053892] | 0.013355(*) |
|  |  |  | Sham | 7 | 267(708) | -0.015467 | [-0.034758 0.002421] |  |

## Code and data sharing

Code and data are available at https://gitlab.com/sainsbury-wellcome-centre/delab/publications/soundmap-2025.

## Acknowledgments

We thank Lillianne Teachen, Yingkun Li, Aline Rampini, Nengneng Gao, Santthia Rajagobalan, Rebecca Bell, Mingming Chen and Nicolas Chapelot for training animals and technical assistance. We thank Rob Campbell and SWC Advanced Microscopy Facility for the infrastructure for brain imaging. Chunyu A. Duan, Tiago Branco, Thomas Mrsic-Flogel, Agostina Palmigiano, Tim Behrens, Stephen Fleming, Toby Wise, Matteo Carrandini, Nuo Li, Jenny Bizley, Edmund Chong, & Ivana Orsolic gave helpful comments and feedback at different stages of the project. J.L. was supported by Sainsbury Wellcome Centre PhD program from the Gatsby Charitable Foundation (GAT3755) and Wellcome Trust (219627/Z/19/Z). J.L. and L.L. were supported by the NYU Shanghai Doctoral Fellowship, Winston Foundation Fund, and Bright Oceans Corporations Fund for Science and Research. The research was also supported by grants to J.C.E. from: the Program of Shanghai Academic/Technology Research Leader (15XD1503000); the Science and Technology Commission of Shanghai Municipality (15JC1400104); the 111 Project, Base B16018; the National Natural Science Foundation of China (NSFC; 31970962); NYU-ECNU Institute of Brain and Cognitive Science at NYU Shanghai; the Sainsbury Wellcome Centre Core Grant from the Gatsby Charitable Foundation (GAT3755,GAT4057) and Wellcome Trust (219627/Z/19/Z,318818/Z/24/Z).

## Contributions

J.L. and J.C.E. designed the behavioral task and conceptualized the study. J.L. setup the systems for optogenetics and electrophysiology recording, with C.B., and developed enhancements to the automated training facility. J.L. implemented and programmed the task and training pipeline. A.X. and J.L. trained the SLEAP model for video analysis and J.L. analyzed and quantified the video data. J.L. analyzed the behavioral data. C.B. supervised the development of surgical techniques. J.L. and C.B. performed the surgeries with A.X. and L.L. assisting. J.L., A.X., C.B., L.L. collected the electrophysiology and optogenetics data. J.L. developed the analysis pipeline and analyzed the data from electrophysiology with A.X. J.L. analyzed the data from the optogenetics experiments.

A.A. and J.L. implemented the dynamical model under the supervision of C.C. The paper was written by J.L. and J.C.E. with comments from the other authors. J.C.E. supervised all aspects of the work.

## Declaration of generative AI and AI-assisted technologies in the writing process

During the preparation of this work the author(s) used large language models, including ChatGPT, Claude and DeepSeek R1, in order to provide programming and writing assistance. After using these tools/services, the authors reviewed and edited the content and take full responsibility for the content of the published article.

## Methods

### Subjects

A total of 3 male Sprague Dawley rats (Vital River, Beijing, China) and 29 male and female Lister Hooded rats (#603, Charles River, London, UK) were used for this study. Details of each animals role in the study can be found in Table S2. Animals were paired housed during the training period and then single housed after electrode or fiber implantation. Of these 32 animals, 9 Lister Hooded rats were used for FOF optogenetics experiment, another 23 rats were used for *in vivo* electrophysiology recording experiment in FOF, mPFC, SC, RSC, Hippocampus and AC. Animals were kept on controlled-wateraccess in order to motivate them to work for water rewards. Body weight was closely monitored daily to ensure their health according to the relevant animal welfare guidelines. All rats were kept on a reversed 12 hour light-dark cycle and were trained during their dark cycle. Animal use procedures for 3 Sprague Dawley rats were approved by New York University Shanghai International Animal Care and Use Committee follows both US and Chinese regulations. For 29 Lister Hooded rats, animal use procedures were approved by the Home Office of the United Kingdom (Project License: PE4FA53CB).

### Behavioral apparatus

Animal training took place in custom behavioral chambers inside boxed to attenuate sound and light. Each chamber (23 23 23 cm for training rigs and 23 23 45 cm for recording rigs) was fit with 8 nose ports arranged in four rows (Fig. 1a), with stereo speakers located on the left and right side. Each nose port contained a pair of blue and a pair of yellow light emitting diodes (LED) for delivering visual cue, as well as two infrared LED/phototransistor pairs for detecting rats’ interactions with the port. The port in the bottom row contained a stainless steel tube and a valve for dispensing precisely measured liquid rewards (the reward port). The LEDs, speakers, water valves and the IR pairs were controlled by Bpod), with some design improvements: more I/O to control more LEDs; used a hysteresis comparator array to avoid jittering of IR beam breaks during detection of pokes; and added better ground/isolation layout to prevent ground loops. Additionally, we integrated all of the LEDs and IR pairs into a single PCB (the port wall, as in Fig. 1a) which was powered by 5 V input and connected to the bpod main hardware with a single ribbon cable. Speaker volume was calibrated to 78 dB peak amplitude using uniform white noise. Those designs greatly improved the stability and reliability of the behavioral system. Hardware designs and related software are freely available (bpod-auto).

### Behavioral task and training pipeline

We developed two task variations of auditory memory-guided orienting task^4^ for two groups of water-deprived rats. Both task variations followed the same timeline. At the beginning of each trial, one of the operant ports was randomly illuminated as the start port. Rats were required to fixate in the start port for 1.5 s until a go cue, constituting the fixation period. During the first 0.75 s of fixation, one of two auditory cues was presented: 30 Hz or 68 Hz click trains, generated using `sound.make_clicks`. Each click was an 8 kHz pure tone lasting 0.003 s. This was followed by a 0.75 s silent delay period. A go cue, 0.2 s 15 KHz pure tone, was played at the end of the delay. After the go cue, rats withdrew from the start port and moved to a target port. A correct response resulted in the illumination and water reward availability at the reward port. Incorrect choices triggered an error sound (1 s white noise), and a new trial became available after a short inter-trial interval (2 s to 10 s; Fig. 1a). Early withdrawal from the start port before the go cue, required animals to restart the fixation from the beginning. These trials (about 20%) were excluded from analyses except when calculating early withdrawal rates (Fig. S7).

The mapping from sound cue to target port depended on the task context and start port:

- **Egocentric (Ego) Task:** The correct target port was either the left or right port adjacent to the start port, determined by the sound cue (blue/red for 30 Hz/68 Hz clicks, respectively; Fig. 1b, left). This resulted in eight trial types spanning two movement directions (Fig. S1a, left).
- **Allocentric (Allo) Task:** The correct target port was fixed in physical space, with the bottom-left/bottom-right port corresponding to the blue/red sound cues, respectively (Fig. 1b, right). This allocentric spatial coding yielded 12 trial types from the combination of different start ports and target ports(Fig. S1a, right).

We classified errors in the task in two ways. The first, in-task errors, were errors where the subject’s choice port was associated with the unplayed sound. The second, out-of-task errors, were errors where the subject’s choice port was not associated with either sound. These later choices were rare and were excluded from all analyses except Fig. S1b.

#### Training pipeline

Rats underwent a structured training pipeline towards achieving mastery of the final task. Training comprised three phases:

1. **Operant Conditioning Phase:** In this phase, rats became familiar with the training apparatus and learned to associate illuminated ports with water rewards. Training progressed through distinct stages:

- **Stage 1:** The reward port was illuminated, and rats received water upon entry. Advancement to the next stage required six consecutive correct trials.
- **Stage 2:** All operant ports were illuminated, and poking in any lit port activated the reward port. The number of illuminated ports was gradually reduced from eight to one. Progression required six consecutive successful trials without errors (e.g., pokes in unlit ports) or timeouts (no poke within 60 s of trial initiation).
2. **Auditory Cue and Task Rule Phase:** In this phase, rats learned the association between auditory cues and target ports. Fixation was not required; rats could initiate a trial by poking the start port, after which the auditory cue was played until choice. Initial training used blocked trial types, with block switching probabilities increasing after each correct trial (starting at 0.02).
3. **Fixation and Timing Phase:** This phase introduced fixation and timing requirements. In this phase, the sounds were played for a maximum of 1.1 s. Initial fixation times were set to 200 ms and adaptively increased by 20 ms increments based on task performance until reaching 1500 ms. As fixation grew beyond 1.1 s, the task required some short-term memory. Once fixation stabilized at 1.5 s, auditory cue durations were reduced from 1.1 s to 0.75 s using a similar adaptive approach. The progress of animals through this stage is shown in Fig. S2.

Rats were trained daily for 80 minutes. The complete training process typically required 2 to 4 months (Fig. S2).

For 12 animals, after reaching asymptotic performance on the Allo task and collecting either electrophysiological or optogenetics data, we transformed the task to a ‘reduced 2AFC’ task, where all trials started from the center-port in order to perform withinsubject tests of how reducing the working memory load for a motor planning would change animals’ strategy (Fig. 7 & Fig. S10).

### Surgery

The rats were anesthetized with isoflurane and placed in a stereotaxic apparatus (David Kopf Instruments). The scalp was shaved, washed with ethanol and iodopovidone and incised. Then, the skull was cleaned of tissue and blood. The stereotax was used to mark the locations of craniotomies for the FOF+mPFC (+2.5 AP, ±1.4 ML mm from Bregma); AC (−4.3 mm AP, ±5.5 ML); mPFC only (3.2 mm AP, ±1.0 mm ML); Hippocampus (4.0 mm AP, ±3.0 mm ML); RSC (−5.6 mm AP, ±1.3 mm ML) or SC (−6.4 mm AP, ±1.3 mm ML), relative to Bregma on the skull. Then a 1.5 mm craniotomy was drilled, followed by an entire dura resection. The craniotomy was then filled with saline saturated Gelfoam to protect the brain tissue while the skull was coated with a thin layer of C&B Metabond (Parkell, Inc; New York) and a 1-3 mm high moat built around the craniotomy using the Absolute Dentin (Parkell, Inc; New York).

For electrophysiology surgery, silicon probes (Cambridge NeuroTech/Neuropixels) were implanted into the target region. The silicon probes were adhered to nano-drives (Cambridge NeuroTech, or R2Drive^95^) with super-glue before implant. Ground wires were soldered to titanium ground screws located above primary visual cortex. The silicon probe was slowly lowered into the brain until the desired depth (FOF recording: 1.3 mm DV for the H3 probes, and 0.5 mm DV for the E probes and F probes; FOF+mPFC recording: 3.6 mm DV, Neuropixel 1.0 probes with 10 degree for ML; AC recording: 4.8 mm DV, Neuropixel 2.0 probes with 10-12 Degree in ML; mPFC recording: 3 mm DV for H3 probe; RSC recording: 1.5 mm DV for H3 probe; SC recording: 5.5 mm DV, Neuropixel 1.0 probes; Hippocampus recording: 3.6 mm DV, Neuropixel 1.0 probes).

For optogenetics surgery, we slowly injected 400 nL AAV virus solution (pAAV-CKIIastGtACR2-FusionRed, AddGene #105669, about 5 × 10^12^) into FOF bilateraly. The injection system comprised of a custom-made glass pipette (20–30 µm O.D. at the tip; Drummond Scientific, Wiretrol II Capillary Microdispenser) back-filled with mineral oil, and a fitted plunger inserted into the pipette to load or dispense viral solution. The plunger was controlled by a hydraulic manipulator (Narashige, MO-10); and the injection pipette was advanced into the brain using stereotax. After each injection, the glass pipette was left in the brain for 5 minutes before it was slowly withdrawn, to prevent backflow. Tapered fibers (`MFC_200/245-0.37_4.5mm_ZF2.5(G)_TP1.5`, Doric Lenses Inc.) were inserted 1.2 mm into the cortex measured from the brain surface for each craniotomy for bilateral FOF. A sham fiber was also implanted at the posterior part of the skull without entering the brain.

The craniotomy was filled with Dura-Gel (Cambridge NeuroTech), and then silicon probes or the optic fiber was cemented to the skull with Absolute Dentin. Vetbond (3M, U.S.) was applied to glue the surrounding tissue to Absolute Dentin. The animals were given 5 days to recover on free water before resuming training.

### Electrophysiology recording and spike sorting

For Cambridge Neurotech probes, neural activity signal was digitized at 30 kHz, amplified and bandpass filtered at 0.6-7500Hz using a 64 channel Intan headstage (RHD2164, Intan Technologies), the SPI cable of the Intan headstage was tethered to a commutator to allow free spinning (Shenzhen Moflon Technology, MMC250), and all the raw data was processed using the Open Ephys acquisition board connected to a computer to visualize and store the neural signals. The probes were turned down ≈ 100 µm for every 3-5 days until the white matter was reached. For Neuropixel probes, we used acquisition hardware from NI (a PXIe-1071 Chassis) with a Neuropixel recording module (PXIE_1000, IMEC). Data was monitored and recorded with SpikeGLX software.

During the recording, at the end of each trial a serial TTL message encoding the current trial number was sent from our behavioral control hardware to the acquisition system to permit synchronization of the neural signal with the behavior data.

Offline spike sorting was performed by Kilosort v2^96^ for Cambridge Neurotech probe recording or Kilosort v4^97^ for Neuropixel probe recording with default settings. Spike clusters were manually curated using Phy. The quality metrics and waveform metrics for sorted units were computed using ecephys spike sorting^98^. Specifically, we selected units with averaged firing rate >1 Hz, signal-to-noise ratio >1.5, and a presence ratio >0.95 over the course of recording sessions.

### Optogenetics experiment

The implanted fibers were connected to one (for the sham fiber) or two (for bilateral silencing of FOF) 0.6 m optic fiber cables (`OPT/PC-FC-FCF-200/230-HD-0.6L-MC`, Plexon Inc.). The other end of cable was connected to a dual fiber rotary joint (`FRJ_1x2i_FC-2FC_0.50`, Doric Lenses Inc.) mounted in the ceiling of the sound attenuation chamber. This was connected to a 473 nm laser (473nm LX 50 mW LASER SYSTEM, OBIS Inc.) with a variable optic attenuator (VOAMMF, Thorlabs Inc.) and an optic patch cable (M123L01, Thorlabs Inc,). The laser power was controlled by Pulse Pal (1102, Sanworks LLC) which was triggered by Bpod with 5 V TTL. The laser power was calibrated by a laser power meter (PM20A, Thorlabs Inc.) before and after every session to be 3mW at fiber end.

Before FOF silencing experiments, animals were habituated to tethering and laser delivery through the sham fiber. After habituation, FOF silencing sessions were interleaved with sham sessions. In each session, laser light was delivered on 25% of trials during either the cue or delay period, through the bilateral FOF fibers in FOF sessions or the sham fiber in sham sessions.

### Videography

We used USB web camera (FIT0730, DFROBOT) to record video from top-down view during behavior session. Video were acquired at 640 × 480 pixels and 20 frames per second. Rat performed the task in complete darkness, and videos were recorded under infrared 940 nm LED illumination. We developed a custom software controlled the video acquisition for each session (zmq-video-server).

### Histology

We imaged the brains using serial section^99^ two-photon^100^ microscopy. Our microscope was controlled by ScanImage Basic (MBF Bioscience) using BakingTray, a custom software wrapper for setting up the imaging parameters. Images were assembled using StitchIt.

### Analyses and statistics

In the text, when we use (e.g. 4.2 0.1), we are reporting the mean and standard error of the mean, unless otherwise specified. In any case where a parametric statistic resulted in a *p* value < 10*^−^*^9^, we report it as *<* 10*^−^*^9^. Details of all statistical tests are included in the Statistical Appendix.

For mixed-effects models we used either `fitlme` (for linear models) or `fitglme` (for Poisson or logistic models) in MATLAB (2023a; The Mathworks). In some cases, we used the `lme4` library (version 1.1-29)^101^in R.

#### Behavioral data analysis

Out-of-task errors were rare and were excluded from all analyses except Fig. S1b. We excluded all timeout trials (no choice made within 2 s from go cue) and fixation violation trials (failure to maintain fixation for 1.5 s in the start port, about 20% of trials). Additionally, we excluded trials where animals accidentally used their paw to break an IR beam in a choice port (port entry was detected within 0.2 s after the go cue, 3.71% of trials removed). The hit rate, p(correct), was calculated as the fraction of trials in which the animal reached the correct choice port, relative to all remaining trials after these exclusions (Fig. S1g). Movement time (MT) was defined as the interval between withdrawal from the start port and entry into the choice port. For this analysis, we included only center-start trials. To compare MT differences between the Allo and Ego tasks (Fig. 1d and Fig. S1c), we first computed the median MT for each animal. We then used the Wilcoxon rank-sum test (`wilcox.test` in R) to assess the statistical significance of differences between the Allo and Ego animal groups. The *p* values or that test are reported in the respective figure panels.

To statistically assess the MT change after task transformation (Fig. 7c), we performed day-by-day likelihood-ratio tests using log-transformed MT values. For each day after task transformation, 0 ≤ *d* ≤ 7, we compared baseline trials, `pre` (all valid trials from the 7 days before the task transformation) to trials from day *d*, `post` using mixed-effects linear models of the form:

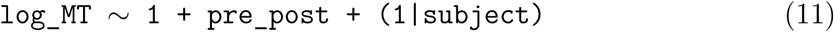

Where `pre_post` indicated whether a trial belonged to baseline or to after task transformation. This full model was compared to a reduced model without the `pre_post` term using a likelihood-ratio test. The resulting p-values determined which post-transformation days differed significantly from baseline. The same model was used to compare pooled MT (from all tethered experimental sessions) before and after task transformation (Fig. S10g). In optogenetics data analysis, we also defined reaction time (RT) as the interval between the end of the delay period (indicated by the go cue) and the withdrawal from the start port (Fig. S7a,b).

#### Video data analysis

We used SLEAP^102^ to track body key points (nose, left ear, right ear, neck, Fig. 1e) from video frames recorded for each session. Separate models were used to track animals that have different implant type (optic fiber, or recording probe). We manually labeled ≈3000 randomly picked frames from different sessions to train each model. Training was performed using the default setting parameters of SLEAP. Then the trained model was used to predict the position of each key point for all frames of all sessions. We then calculated vector angles between the pair of each point using the x,y coordinate of each points.

For each trial, the vector data (3-dims: left-ear, right-ear, neck to nose angle, Fig. 1e) was aligned to the trial onset (using `cdraster` function in Erlich lab utilities^103^, Fig. 1f), or choice port (target) arrival. At each timepoint for each individual rat, we used logistic regression with LASSO regularization and 10-folds cross-validation that took the vector angles as input to predict the animals’ choice (using `fitclinear` function in MATLAB). We concatenate all data from all sessions for each individual rats. Only center-start trials where the errors were ‘in task’ were included for the video posture analysis to control for the same up-coming movement after the go-cue. We averaged the choice decoding accuracy of all rats at each time-point for each task. This result was shown in Fig. 1g and Fig. S1d.

### Electrophysiology data analysis

#### Characterizing single-neuron selectivity

For the rasters and peri-stimulus time histogram (PSTH) plots (Fig. 2), spike times were aligned to the sound cue onset within a 2.5 s time window (−0.5 s before cue onset to 0.5 s after fixation end). The bin size was set to a resolution of 0.01 s and smoothed with a causal half-Gaussian kernel with a standard deviation of 0.2 s.

To evaluate whether the activity of each cell during the delay period (0.75 s to 1.5 s after cue onset) could predict upcoming choice, `spike_count` was calculated for each trial during this period. To quantify choice tuning of each cell, we ran a mixed-effects Poisson model with the following formula.

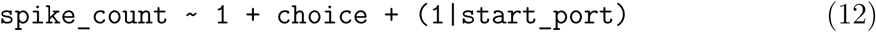

where `choice` was −1 or 1 for ipsilateral or contralateral choice (relative to the recorded hemisphere) and `start_port` (−2 to 2) denoted the X-axis location of the start ports relative to the center port. The choice was defined by contra/ipsi target direction for the Ego task or contra/ipsi target port for the Allo task. A *p <* .01 threshold for the `choice` coefficient identified choice selective cells.

A similar mixed-effects linear regression model was implemented to evaluate the contribution of start port position to firing rates in the FOF cells, independent of choice. The formula was:

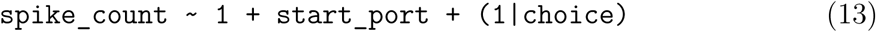

so that we could quantify the contribution of `start_port` independent of `choice`. A *p <* .01 threshold for the `start_port` coefficient identified start-port-selective cells.

To assess choice selectivity over the trial’s time course for center-start trials (Figure 3a), we applied a linear regression model:

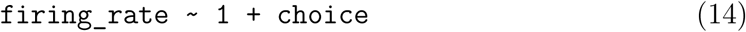

where `firing_rate` was estimated with a 0.25 s sliding window (0.025 s step size) from spike times aligned to cue onset. A *p <* 0.01 threshold for the `choice_index` coefficient was used to identify choice-selective cells at each time point. The percentage of such choice-selective cells was then calculated across time points.

To assess start-port selectivity over the trial’s time course, we used the same model as Eq. 13 using a 0.25 s window and 0.025 s step size aligned to cue onset. Cells with *p <* 0.01 for `start_port` were identified as start-port selective, and the percentage of such cells was calculated over time points.

For units recorded from auditory cortex (Fig. 5b), we quantified cue selectivity at 1.3 s after cue onset using spike counts from a 300-ms window (1.15 s to 1.45 s). For each neuron that have firing rate for at least 0.5 Hz, spike counts were z-scored across the included correct and error trials, and cue coefficients were estimated separately for the two outcomes using a Gaussian mixed-effects model:

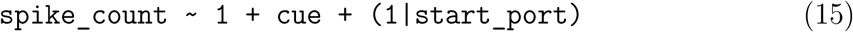

The cue coefficient, *β*_cue_, quantified the strength and direction of cue selectivity. Cells with *p <* .05 for *β*_cue_ on correct trials were classified as cue selective. To determine whether cue coding was preserved on error trials, we fitted a total least-squares regression between the correctand error-trial coefficients separately for each task. Confidence intervals for the slope were estimated by resampling neurons 10,000 times and reporting the 95% bootstrap confidence interval. Pearson correlations between correctand errortrial coefficients were compared between tasks using Fisher’s *r*-to-*z* test.

We extended this analysis across the trial time course (Fig. S8a,b), firing rates were estimated using a non-causal 0.2 s flat sliding window with a 0.025 s step, aligned to cue onset. Neurons with a mean firing rate greater than 0.5 Hz across the analyzed trial window were included. At each time point, we fitted the Gaussian mixed-effects model described above separately to correct and error trials. We then fitted a total least-squares regression between the correctand error-trial cue coefficients, using only neurons with significant cue coding on correct trials at that time point (*p <* .01). This analysis was performed after pooling neurons within each task and separately for each animal. For the task-level analysis, 95% confidence intervals were estimated by resampling neurons 5,000 times. We tested whether each task’s slope differed from zero using the bootstrap distributions and compared slopes between tasks using a permutation test with 5,000 shuffles of task identity. Statistical tests were corrected across time points from 0.6 s to 1.7 s using FDR correction (*p* < .05).

#### Pseudopopulation decoding

Following previously described approaches^15,104^, all pseudopopulation decoding was performed on pseudo-trials pooling neurons recorded across sessions and animals within each task. To compare choice decoding performance, we generated 100 pseudosessions for each task: both for Ego vs. Allo comparison and also for Allo vs. 2AFC. For each pseudosession, we randomly sampled 100 cells without replacement (see Fig. S10b for choice decoding as a function of population size in the Allo and 2AFC tasks). For each selected cell, we independently sampled without replacement 10 left-choice and 10 right-choice center-start trials from the pool of eligible trials in its recording session. Responses with the same choice label were combined across cells to form 20 balanced pseudo-trials. We included both correct trials and in-task error trials for this analysis. Sessions with fewer than 10 left and 10 right choice trials were excluded. For each selected cell, we estimated the firing rate across each trial with a 0.25 s causal sliding window with a 0.025 s step size, aligned to cue onset. This process yielded a 100 × 20 × 100 matrix (cells × trials × timepoints), *D*_original_, for each pseudosession.

Principal component analysis (PCA; using pca in MATLAB, which automatically centers data) was then applied to *D*_original_, with distinct approaches depending on the analysis type: comparing decoding across tasks (Fig. 3b, Fig. 7g) or cross-time decoding (Fig. 3d, Fig. 7f). For comparing decoding across tasks, PCA was performed independently for each timepoint (each timepoint giving a 100×20 (cells×trials) matrix), and the top five principal components (PCs) were used. Projecting the data onto the top 5 PCs resulted in a matrix *D*_independent_ of dimensions 5×20×100 (PC dims×trials×timepoints) for decoding analysis. For cross-time decoding, *D*_original_ was reshaped, creating a 2D matrix of dimension 100×2000 (cells (timepoints trials)), denoted as *D*_concatenated_. PCA was applied to *D*_concatenated_, we used top five principal components and reversing the trial concatenation, to create a 5 × 20 × 100 (PC dims trials timepoints) matrix, *D*_shared_, for decoding analysis. Decoding was performed using multivariate linear classification (`fitclinear` in MATLAB with L1 regularization) with leave-one-trial-out cross-validation over the 20 trials selected in each pseudosession.

To generate a *p*-value at each time for the comparison of decoding (Fig. 3b, Fig. 7g), we generated a 100 timepoints vector, *A⃗*_data_, by averaging the decoding accuracy over pseudosessions. We then generated a shuffled decoding distributions for each of the 100 pseudosessions by permuting the label of left/right for the 20 trials 5000 times and doing the decoding analysis for each shuffle, producing a 100 pseudosessions 5000 shuffles for each time point and averaging over pseudosessions to get a 5000×100 (pseudosessions timepoint), *A*_shuffle_. We considered a timepoint in *A*_data_ to be significantly above chance if it was higher than 0.9995 (p=0.05 / 100 using Bonferroni correction) of the corresponding shuffle distribution for that timepoint. To test if two tasks were different from each other we did a permutation test between the *A⃗*_data_ with 10,000 shuffles of the task label. For visualizing the timepoints where the tasks has significantly different, we required at least 5 consecutive timepoints to be significant in order to be included. For comparison between the Allo vs. 2AFC task, we required at least 8 consecutive time points to be significant (to account for a greater degree of noisiness in the smaller data set).

For cross-time decoding analyses (Fig. 3d, Fig. 7f), we trained a classifier for each pseudosession at every timepoint in *D*_shared_ using the label vector *L*. The classifier predicted the labels across all timepoints of *D*_shared_ with leave-one-out cross-validation, yielding a 100×100 (timepoint×timepoint) matrix of decoding accuracy that captured the stability of choice coding^105^. To assess statistical significance, we generated 1,000 null matrices by shffling the label vector *L* for each pseudosession. An extreme pixel-based permutation test^106^ was applied to compute *p*-values for each matrix element and a contour was plotted to indicate significant regions. Significant regions smaller than 7 × 7 timepoints were excluded.

To compare start position decoding we used the same methods and code as for choice decoding with the following key differences. Instead of using only center-start trials we sampled 10 left and 10 right choice trials (from correct trials and in-task error trials) for each of the top-left, top-right, and center-start trials without replacement, yielding 60 trials for each pseudosession. We used only these three start positions because they are matched across the two tasks (Fig. S1a). For each trial, we coded the X-axis position of each port (−1 for top-left port, 0 for center port and 1 for top-right port). Decoding was performed using multivariate linear regression models (`fitrlinear` in MATLAB) with 10-fold cross-validation. The decoding performance was quantified as the R^2^ between the true start position and the cross-validated model prediction for each trial.

#### Single-trial decoding of choice

To perform single-trial decoding of choice for each session (Fig. 3c; Fig. S4e-h), we selected center-start trials that were correct or were in-task errors to generate an *N*×*I*×*T* data matrix (neurons × trials × timepoints), *D*_sess_. We also had a corresponding length *I* vector for each session: *L⃗*, which was set to 1 if the animal chose right and set to 0 for chose-left on each trial. PCA was performed independently at each timepoint to perform PCA whitening. This resulted in a matrix *Z*_session_*_,t_* of dimensions *P × I* (post PCA dims × trials) for each timepoint, *t*. Then, we performed binary classification of *L⃗* using *Z*_session_*_,t_* (`fitclinear` in MATLAB with L1 regularization) with leave-one-out cross-validation. This resulted in a length T decoding accuracy vector. We performed this for each session. For Ego task FOF recording sessions (n=59), we had on average 184 ± 6 completed trials per session, with ≈ 29.47% centerstart trials per session. In Allo task FOF recording sessions (n=65), we had on average 251 ± 12 completed trials per session, with ≈ 20% center-start trials per session.

To compute the latency of the rise in choice decoding performance, we evaluated the time, for each session, at which the decoding accuracy reached 0.7 for at least 7 consecutive timepoints, based on a previously reported method^47,107^. Latency comparison plot was generated with DataViz (Fig. 3c).

#### Comparison of single-trial decoding of choice across regions

For the comparison of decoding accuracy of choice across regions (Fig. 5f,g,h), we only included correct trials to simplify comparison of the auditory cortex with other regions where error trials may be in different reference frames. That is, errors in AC are generally in a sensory reference frame, and errors the in frontal cortex are in a choice reference frame. On correct trials, decoding the choice is equivalent to decoding the sensory cue. We also excluded trials that started from the bottom-left or bottom-right ports.

In contrast to the causal sliding window used elsewhere (e.g. Fig. 3), here we used a 250 ms non-causal sliding window (25 ms step) to make it easier to estimate the decay of cue-coding during the delay. We generated an *N*×*I*×*T* (neurons × trials × timepoints) matrix, *D*_sess_, together with the length I trial label vector *L⃗* (0 for sound A, 1 for sound B). PCA whitening was applied to at each timepoint, giving *Z*_sess_ of size *N*×*I*×*T* . We then added the start-port as a dummy variable to *Z*_sess_ to allow the decoder to find start-port dependent coding of choice. Decoding was performed at each time point using logistic classification (`fitclinear` in MATLAB with L1 regularization) with 10-fold cross-validation, resulting in decoding accuracy curves for each session.

To compare regions, we averaged decoding accuracy during the delay period (0.75– 1.5 s), ranked sessions by accuracy within each region, and selected the top half for group analysis. Trial labels were shuffled 10,000 times for each session, and the same ranking and selection were repeated for each shuffled set. To balance different numbers of sessions across regions, we randomly resampled seven of the top-half sessions 200 times and compared their average accuracy against shuffled controls. We picked 7 because we only had 15 recording sessions from auditory cortex, thus the top half gave us 7 sessions. A timepoint was considered significant (Fig. 5f, purple and red bars) if more than 50% of resampled sets gave *p <* .01; significance periods shorter than 0.075 s were excluded (3 timepoints).

To test whether AC decoding exceeded that of other regions (Fig. 5f,g) we compared session-wise decoding accuracy between AC and each non-AC region (FOF, SC, or mPFC) using a Wilcoxon rank-sum test. For Fig. 5f (grey bar), we show the comparison between AC and FOF, applied FDR correction at *p* < .05, and significance periods shorter than 75 ms were removed.

For single-session cross-time decoding (Fig. 5h), we only included the top half of sessions for each region (described above). We generated a trial-concatenated matrix (neurons × [timepoints · trials]), fit a single PCA, for whitening, for each session on this concatenated matrix, and kept all components. The data were then reshaped back into a representation of *P* × trials × timepoints, so that the same PCA basis was used for all timepoints within a session. Decoding was performed with logistic classifiers (`fitclinear` in MATLAB, 10-fold cross-validation with L1 regularization), training at each timepoint and testing at every timepoint, which produced a cross-time decoding heatmap for each session. For each region, we randomly resampled 7 sessions and averaged their heatmaps, repeating this 200 times to obtain 200 resampled heatmaps. Each resample was compared against 1,000 trial-label–shuffled dummy heatmaps processed with the same selection method using the extreme pixel-based permutation test^106^, yielding a pixelwise *p*-value map. A pixel was considered significant if more than 50% of resamples reached *p <* .01 at that pixel. Small significant clusters (*<* 5×5 timepoints) were removed.

To compare decoding between hit and error trials (Fig. 5d), we constructed two data matrices of dimensions *N*×*I* (neurons×trials) for the spike count from the cue period (0–0.75 s) and the delay period (0.85–1.5 s; the 0.1 s gap was to reduce contamination from the sound encoding during the cue period). For each session, decoding was performed separately for cue and delay periods and for hit and error trials, using logistic classification classifier (`fitclinear` in MATLAB, 10-fold cross-validation with L1 regularization). For cue/delay period, decoding accuracies from hit and error trials were paired across sessions, and differences were assessed using the Wilcoxon signed-rank test.

#### Single-trial decoding of time

To decode the time in trial during the fixation period (Fig. 6h,i). For each session, we estimated the firing rate with a 400 ms sliding window (100 ms step size) from spike times aligned to cue onset, this gave an *N*×*J* matrix, *D*_time_, for each session, where *N* is the number of neurons and *J* is the number of samples for time decoding. With our sliding window over 1.5 s fixation period this gave 16 windows per trial, so *J* was 16 · *I*, where *I* is number of trials in a session. PCA was then performed on *D*_time_ for whitening, all PCs were kept. We also had a corresponding length 16 time label vector, *t⃗*, which was the time relative to the trial onset. Then, we used multivariate linear regression (`fitrlinear` in MATLAB) to predict *t⃗* with 10-fold cross-validation, resulting in *t̂*. We then calculated *R*^2^ between *t⃗* and *t̂* for each session to quantify time decoding performance.

#### Coding direction analysis

To explore how choice coding could generalize across startports we found the ‘choice-coding direction’ in neural population space from center-start trials (Fig. 3i-l)^5^.

To find the coding direction for each session, we generated an *N* × *I* (neurons × trials) matrix, *F*, of the average firing rate of each cell, on center-start trials, from 0.75 to 1.5 s relative to cue onset. We then train a logistic classifier to predict choice from *F*, using the default *L*1 regularization (`fitclinear` in MATLAB). This resulted in a weight vector, *CD*, the coding direction for center-start trials. We repeated this procedure using 10-fold cross-validation to quantify the projection of each center-start trial onto the *CD*. We used all center-start trials to find a *CD* for out-of-sample prediction of choice from other start positions.

To project each trial onto the CD, we estimated the firing rate using a 400 ms sliding window (10 ms step size) to get a length *N* vector at each timepoint, *G_i,t_* representing the population activity on trial *i* at time *t*. We then project the activity onto the *CD*. Projecting the activity for all trials and all times, *G*, gives a *I* × T (trials × timepoints) matrix of the projected activity *Q* = CD^T^ G. We then remove the grand mean of Q to center the projection to allow combining data across sessions.

#### Decoding generalization analysis

To statistically quantify the generalizability of choice coding across different start positions, we trained a multivariate linear classification model (`fitclinear` in MATLAB with L1 regularization) for center-start trials for each session to predict left/right choice label vector *L* from spike counts matrix *D_raw_* during the delay period. Here *D_raw_* is a *N*×*T* matrix for N cells and T trials in that session, *L* is the choice label vector which was 0 for left choice and 1 for right choice.

We firstly predicted the choice label vector *L̂* for all center-start trials with 10-fold cross-validation and quantified the decoding accuracy for each session. For each session, we also predicted the choice label vector *L̂* for other trial types started from other ports, using the classification model trained with all center-started trials of that session as an out-of-sample prediction accuracy.

To quantify the confidence interval of the slope between the decoding accuracy of center-started trials and other trials (Fig. 3m,n), we resampled the data for TLS regression 10,000 times to approximate a distribution of the slope and reported the mean and 95% CI.

#### Cue/choice coding index analysis

For the cue/choice coding index analysis (Fig. 5c; Fig. S8c), we included correct trials and in-task error trials. On correct trials, the cue and choice labels were the same. On in-task error trials, animals chose the target associated with the opposite cue, so that the cue and choice labels were different.

For each session, we estimated the firing rate with a 250 ms non-causal sliding window (25 ms step size) from spike times aligned to cue onset. This gave an *I*_hit_ *× T × N* matrix, *D*_hit_, for correct trials and an *I*_error_ × *T* × *N* matrix, *D*_error_, for error trials, where I is the number of trials, *T* is the number of timepoints and *N* is the number of neurons. At each timepoint, PCA was performed on the correct-trial firing-rate matrix for whitening, and all PCs were kept. We also added the start-port as a dummy variable to the PC scores to allow the decoder to find start-port-dependent coding of the cue. Decoding was performed using logistic classification (`fitclinear` in MATLAB with L2 regularization). We firstly predicted the cue label, *L̂*_hit_, for correct trials with leave-one-trial-out crossvalidation and quantified the correct-trial decoding accuracy, *A*_hit_(*t*), at each timepoint. For each session, we also predicted the cue label, *L̂*_error_, for error trials, using the classification model trained with all correct trials as an out-of-sample prediction. The error-trial firing rates were transformed using the PCA estimated from correct trials before the decoding. This gave the error-trial generalization accuracy, *A*_error_(*t*).

To quantify whether the population activity followed the cue or the animal’s choice, we first centered both decoding accuracies: *H*(*t*) = *A*_hit_(*t*) 0.5 and *E*(*t*) = *A*_error_(*t*) 0.5. We then calculated the cue/choice coding index as

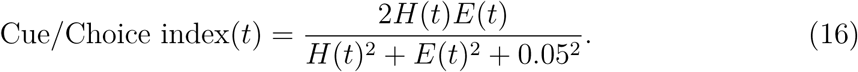

Here, 0.05 was added as a ridge term in the denominator to stabilize the index when both decoding accuracies were close to chance. If cue coding generalized from correct to error trials, *A*_error_ remained above chance and the index was positive. If the decoder followed the animal’s choice, the predicted cue label was reversed on error trials, *A*_error_ was below chance and the index was negative. The index was close to zero when correct-trial decoding was weak or error-trial decoding was close to chance.

For the time-course comparison between tasks (Fig. 5c, left), we averaged correcttrial decoding accuracy from 1.2 to 1.4 s, ranked sessions by hit-trials accuracy separately within the Allo and Ego tasks, and selected the top half of sessions from each task. We used the top half because the cue/choice coding index can only be meaningfully interpreted in sessions where the neural population could decode the cue on correct trials. To test the difference in the cue/choice coding index between the Allo and Ego tasks over time, we shuffled the task label of each session 5,000 times while preserving the number of sessions in each task. For each shuffle, sessions were ranked again by correct-trial decoding accuracy within the shuffled task groups and the top half were selected. Multiple comparisons were corrected using a cluster-mass permutation test with a cluster-forming threshold of *p* < .05 and a cluster-level threshold of *p* < .05.

For the individual-session heatmaps (Fig. S8c), we included all sessions and sorted sessions within each task by their mean correct-trial decoding accuracy during the delay period (0.75–1.5 s). For the session scatter plot (Fig. 5c, right), we showed the data from all sessions. For each session, we averaged both the correct-trial decoding accuracy and the cue/choice coding index from 1.2 to 1.4 s. We compared the summary cue/choice coding index between the Allo and Ego tasks using a permutation test with 5,000 shuffles of the session task labels.

#### Optogenetics data analysis

For optogenetic analyses, we defined control trials as the no-laser trials from the same sessions as a corresponding laser trial. Sham sessions where the sessions when optic fiber was plugged into a fiber which did not enter the brain.

For the session scatter plots in Fig. 4c and Fig. S6e, the colored crosses show the session mean and percentile bootstrap 99% CI for laser-off and laser-on performance separately. Within each task, we resampled session rows with replacement 10,000 times, preserving the pairing of laser-off and laser-on performance from each session. The intervals are the 0.5th and 99.5th percentiles of the bootstrap means on each axis. These are descriptive session-level summaries.

For Fig. 4d and Fig. S6f, the error bars are just for visualization, since all the *p*values are based on mixed-effects models (described below). To generate the error bars, we sampled (with replacement) the laser-on and laser-off trials for each session, took the difference in performance (*laser_ON -_ laser_OF F_* = Δ*p*(*correct*)), and took the mean across sessions. We repeated this process 10,000 times, and show the 95% CI of this distribution as the error bars.

To disentangle the effects of the laser light from its optogenetic effect, we used logistic mixed-effect models (`lme4`, version 1.1-29)^101^ in R (4.4.3) and plotted the results using `ggplot2` (version 3.5.1)^108^. In general, we estimated significance by performing likelihood ratio tests (`lrtest` in the `lmtest` package, version 0.9.40)^109^ between each model with a reduced model that removed the variable of interest.

To test whether bilateral FOF silencing for during different task epochs had an effect on performance, we specified a mixed-effects model where `hit` was a logistic function of laser on/off (`isopto`), fiber location (bilateral FOF or sham, `isFOF`) and their interaction as fixed effects. Session level variability were modeled as random effect in this GLMM (using Wilkonsin notation)^110^:

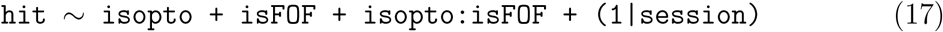

We compared that model with a reduced model

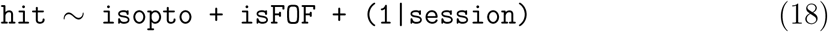

using alikelihood ratio test and reported the *p*-value of that test. By dropping the `isopto:isFOF` we are effectively testing whether knowing if the fiber was connected to the sham or FOF fibers significantly improves the model fit. the This was done 4 times: for (Ego / Allo)×(cue / delay). The p-values are reported in Fig. 4d (delay period) and Fig. S6f (cue period).

However, comparing two groups to baseline is not the same as directly comparing them to each other^111^. To directly compare the FOF silencing effect between the two tasks, we designed a mixed-effect logistic model that considered performance fluctuation, task baseline difference, visual effect of the laser, and most importantly, the interaction between the two tasks and FOF silencing:

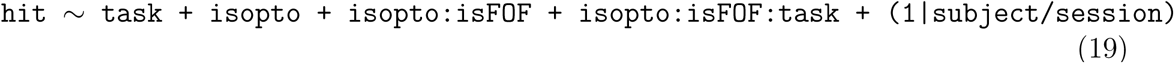

where `task` specified whether a trial was from the Ego or Allo task. and performed a likelihood ratio test against a reduced model without the three way interaction:

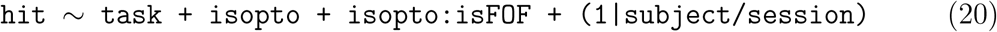

By dropping the `isopto:isFOF:task` interaction term we are effectively testing whether allowing the model to have a differential effect of FOF silencing relative to sham in each task significantly improves the model fit. We used these models to compare the tasks in the cue and delay period separately.

Analyses of the optogenetic effects after task transformation were the same, but rather than comparing Ego and Allo tasks, we were comparing within-subject effects between Allo and the reduced 2AFC task. Note, some small differences between the Allo results in Fig. 4c,d and Fig. 7d,e are due to two animals being included in the comparison with Ego, but not experiencing the task transformation (because of lost implants).

#### Synthetic data for optogenetics experiments

The GLMM model specification for task comparison (Eqs. 19,20) were picked from a set of potential models, based on their ability to discover task differences in synthetic data. We generated synthetic data where there was an effect of silencing FOF in one task but not the other, and included variability across subjects and across sessions. We also matched the trials number and type with our real data (Fig. S6a).

We included, for a sanity check, models (e.g. GLMM0,1) that we know should not fit the data well. The other models included different degrees of complexity (e.g. in the number of parameters) All models shared a similar design that estimated whether a trial would be correct or an error (`hit`) as logistic function of `task`, laser on/off (*isopto*), fiber location (*isFOF*, bilateral FOF or sham). Different interactions among those three terms were implemented to determine the best way to model those interactions. The variability for each sessions, nested into subjects, were modeled as a random effect for all models.

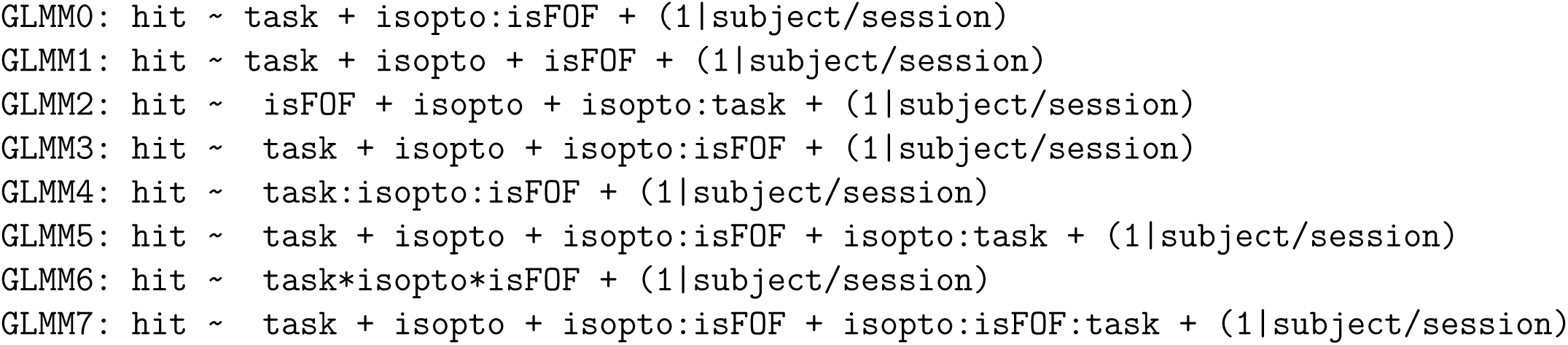

The best model, `GLMM7`, was picked based on the lowest BIC and AIC quantification (Fig. S6b).

### Dynamical model

We developed a computational model to formalize our hypothesis about how different forms of working memory may be recruited under varying task demands. The model has three parts: a selective excitation module and two decision modules. The first decision module is a bistable attractor modeling the FOF as egocentric working memory, FOF. The second module, is a fixed performance modeling the auditory cortex, AC, which supports performance in a sound reference frame. FOF and AC exhibit persistent selective activity in the presence of external excitation, enabling them hold the relevant information over a delay period. In the absence of external excitation the activity decays to zero. We instantiated the decision modules as dynamicals system using Euler’s method (described below).

The excitation to each decision module was based on net cost.

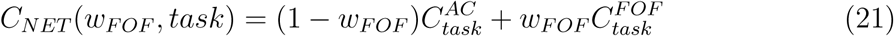

where 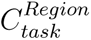 represents the cost of using a mnemonic strategy in a task. 0 ≤ *w*_FOF_ ≤ 1 *s*ets the relative excitation to FOF, and *w*_AC_ = 1 - *w*_FOF_ sets the excitation to AC. Costs are in arbitrary units, and only their ordering matters for the model: motor planning is less costly than sensory working memory in Ego and in the reduced 2AFC task, and more costly in Allo (Fig. 6b). We assigned 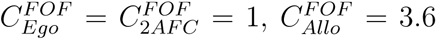= 3.6, and *C^AC^* = 3 in all tasks. These values are illustrative: they were chosen to satisfy this ordering, were not fit to data, and are not estimates of biological costs. The present study does not distinguish the possible sources of cost.

For optimization, we defined the loss as:

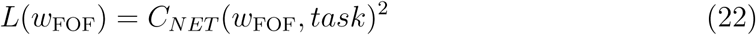

and the trajectory of *w*_FOF_ takes the form

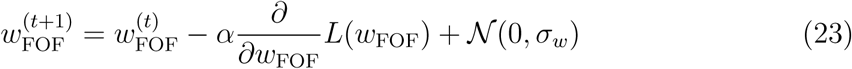

where *α* is the learning rate and *σ_w_* is a parameter controlling noise amplitude. *w*_FOF_ values are clipped between 0 and 1 after each training iteration. Here, *t* represents each training iteration step, and we considered there to be 10 trials per step in the simulation(Fig. 7b), reflecting a slow metacognitive process of assessing the cost of a specific WM format. For completeness, we include the expressions for 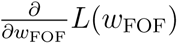 *L*(*w*_FOF_):

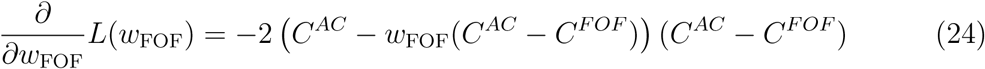

#### Network model of FOF

We modeled FOF/M2 as a binary attractor network following previous work^7,112^. We use a Wilson-Cowan model of cortical interactions^113,114^ with two competing, mutually inhibiting sub-populations, each coding for a different behavioural decision:

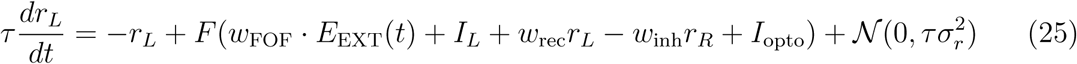

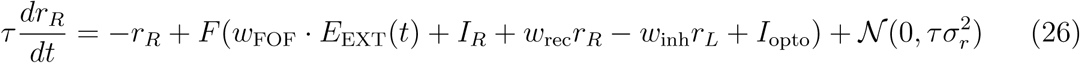

where *τ* is the population rate time constant, *r_i_* is the firing rate of population *i*, driving choice to the right, *r_R_*, or left, *r_L_*. *I_i_* represents the sound input: *I_i_* = 1, if (trialtype=*i*) ^ (0 *< t <* 0.75 s) else 0. *w*_rec_ is the strength of recurrent connectivity within each region, *w*_inh_ is the strength with which populations *L* and *R* inhibit each other, and *σ_r_* is a parameter controlling noise amplitude. *I*_opto_ is the strength of optogenetic inhibition. *F*(*x*) is the activation function, defined as a generalized sigmoid: 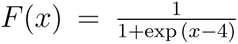. 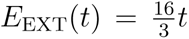 if *t >* 0 else 0. *w*_FOF_ is the gain from the *selective excitation* module. Independent samples of Gaussian noise, 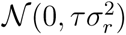, were added to the nodes at each time. We used Euler integration to solve equation dynamics. All parameters used in simulations during learning for the controller are included in Table 6.

#### Cue decoding from network activity

To quantify whether the simulated FOF network encoded future choice during the trial, the simulated activity was organized into a three-dimensional data matrix of size *N* × *I* × *T* (neurons × trials × timepoints). At each time point, population activity across all neurons and trials was extracted as a twodimensional matrix (*I* × *N*) and used as input to a logistic regression classifier trained to predict the cue label (sound A vs. B) with 10-fold cross-validation.

**Supplementary Table 6.** Parameters of network model of FOF.

| **Symbol** | **Description** | **Value** |
|---|---|---|
| *τ* | Population rate time constant | 0.1 |
| *I<sub>i</sub>* | cue input | 1 (cue present) otherwise 0 |
| *w*<sub>rec</sub> | Strength of recurrent connectivity | 1.1 |
| *w*<sub>inh</sub> | Inhibitory strength | 6 |
| *I*<sub>opto</sub> | Optogenetic inhibition strength | -3.0 (opto on)/0 (opto off) |
| *σ<sub>r</sub><sup>2</sup>* | Noise amplitude parameter | 2.25 · 10<sup>−6</sup> |
| *dt* | Discretized timestep | 0.01 |

#### Time decoding from network activity

To assess whether the attractor dynamics in the FOF model represented elapsed time within a trial, we trained linear regression models to predict the true trial time from the instantaneous network state. The same *N* × *I* × *T* activity tensor was reshaped so that each time point from every trial contributed one sample to the decoding dataset, yielding a matrix of size (*I* × *T*) × *N* as the input of regression. The corresponding timestamp for each sample (0–1.5 s, sampled every 0.01 s) were used as the target of regression. A linear regression model was trained to predict trial time from instantaneous network states with 10-fold cross-validation. Decoding performance was quantified as the cross-validated *R*^2^ between the predicted and actual timestamps across all samples.

## 1 Statistical appendix

### 1.1 Figure 2

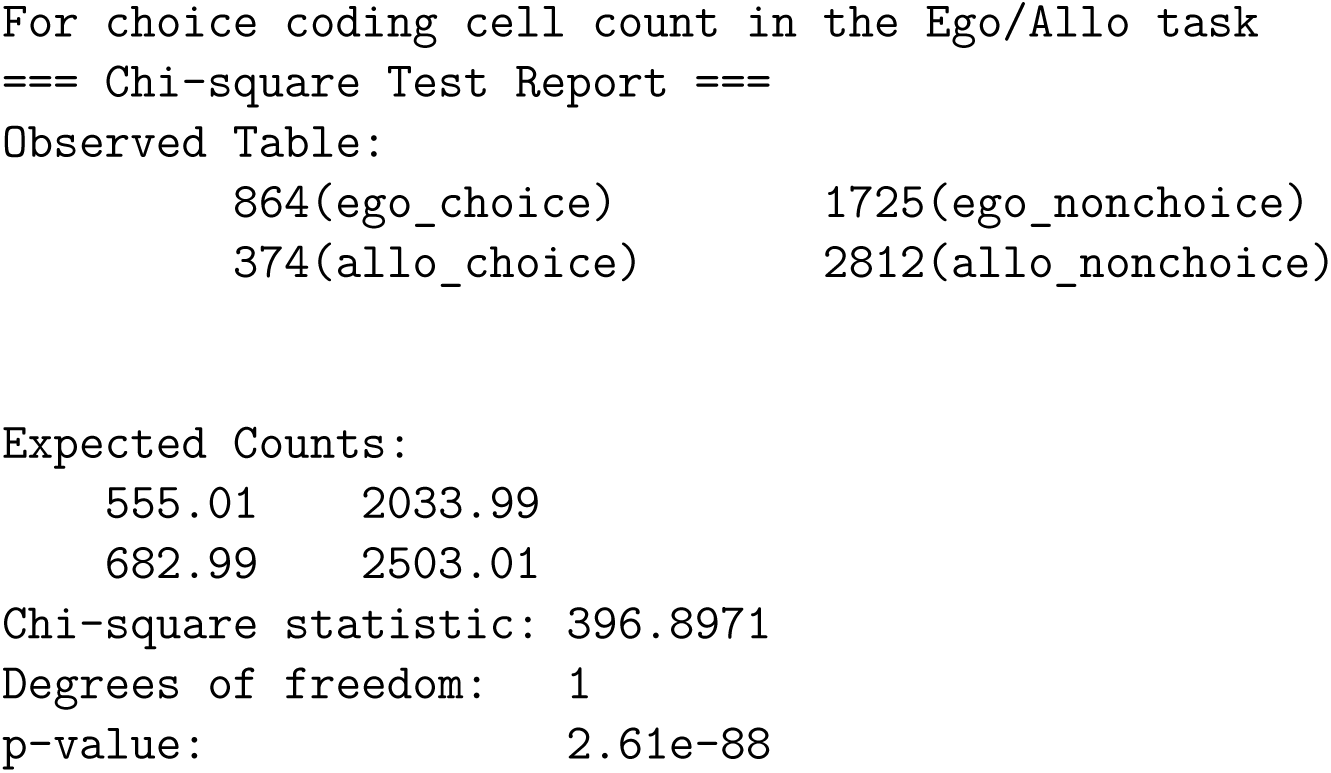

### 1.2 Figure 4 and Fig. S6

#### 1.2.1 Fig. S6f, Left Panel, Ego task, Sham/FOF comparison, Cue period

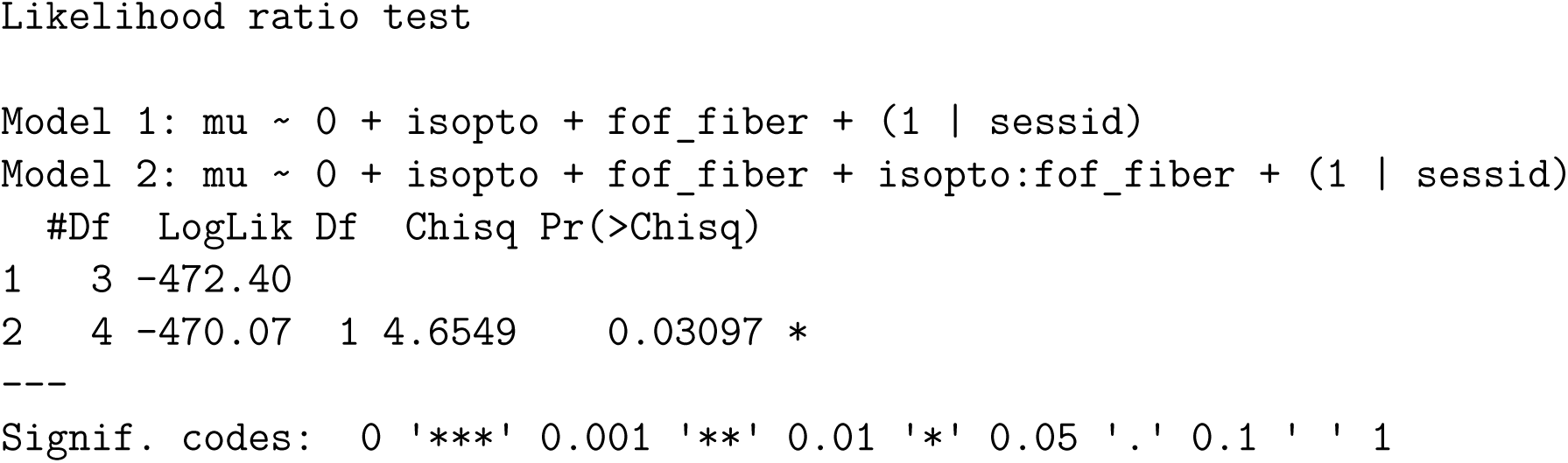

#### 1.2.2 Fig. S6f, Right Panel, Allo task, Sham/FOF comparison, Cue period

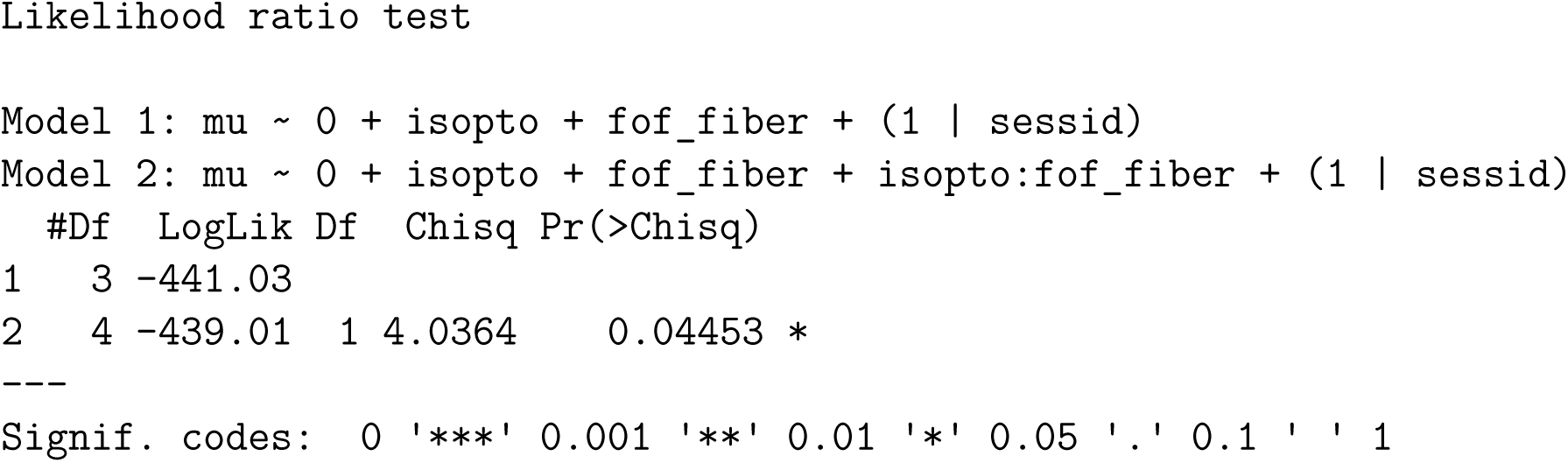

#### 1.2.3 Fig. S6f, Main comparison between Allo/Ego task, Cue period

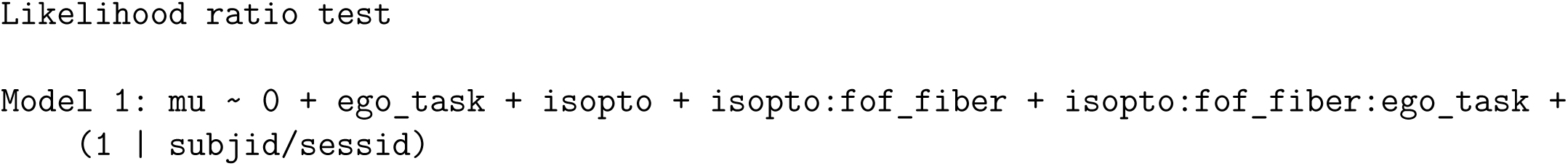
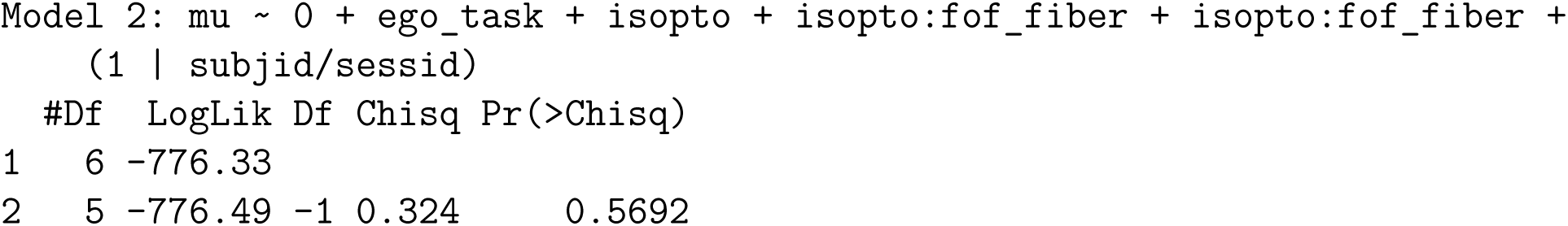

#### 1.2.4 Figure 4d, Left panel, Ego task, Sham/FOF comparison, Delay period

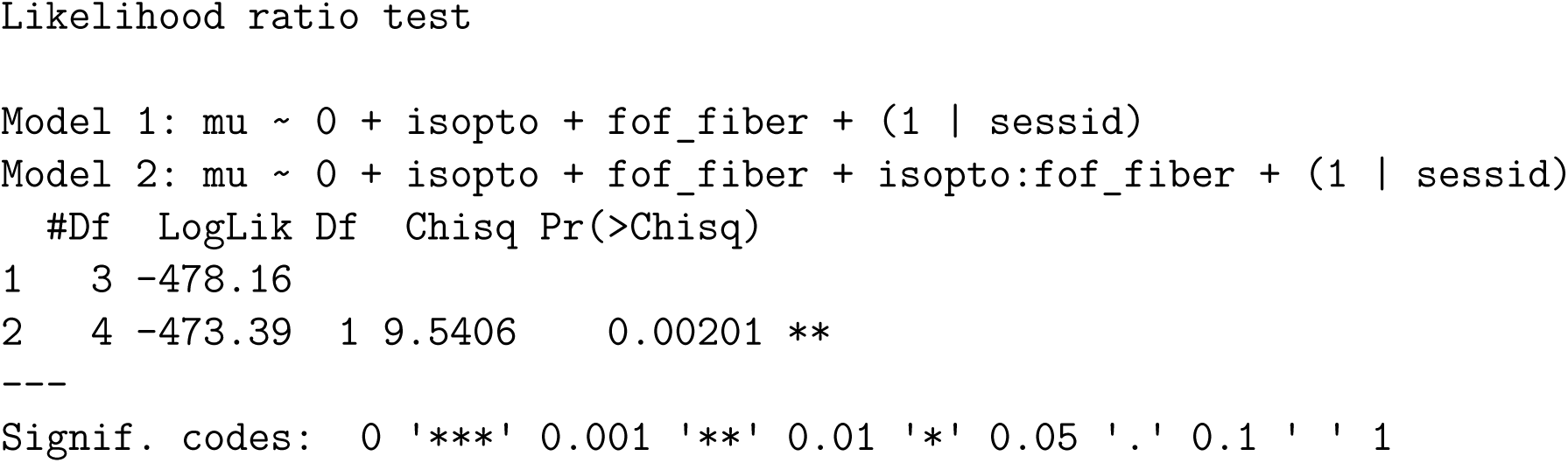

#### 1.2.5 Figure 4d, Right panel, Allo task, Sham/FOF comparison, Delay period

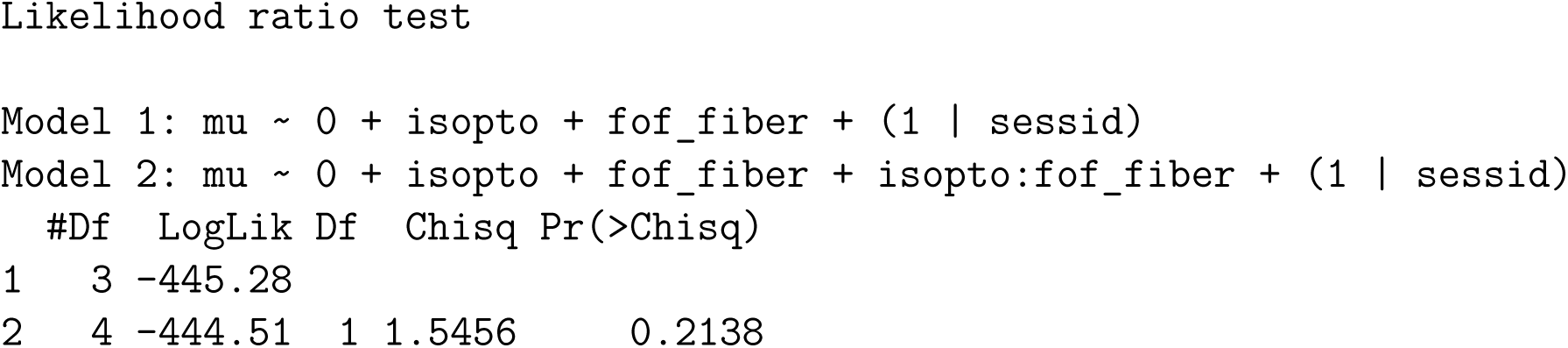

#### 1.2.6 Figure 4d, Main comparison Between Allo/Ego task, Delay period

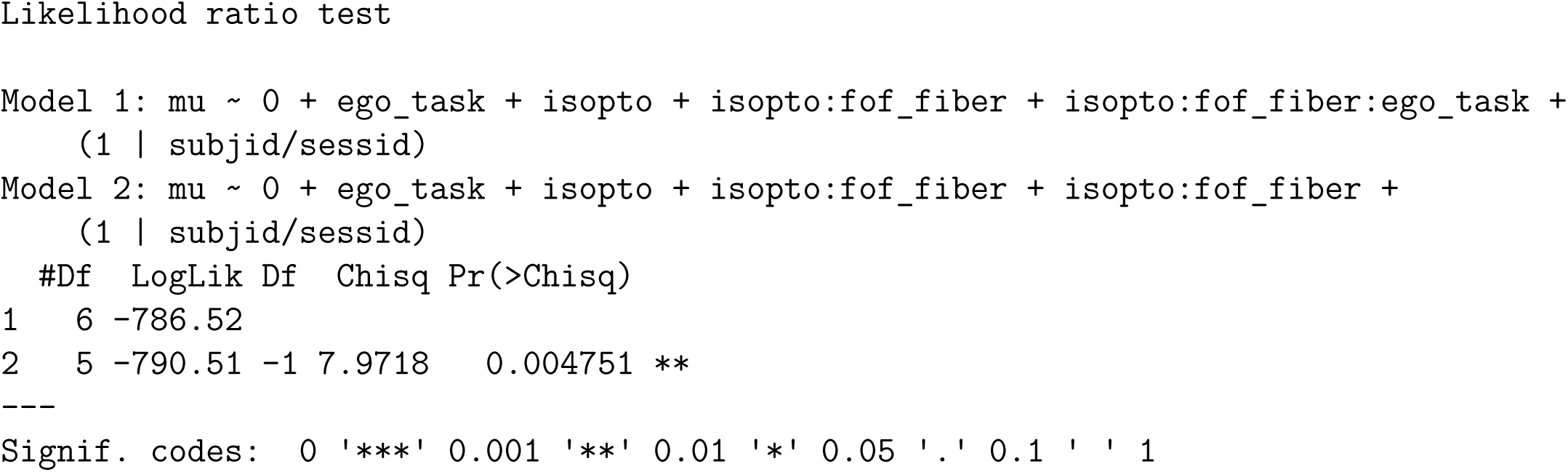

### 1.3 Figure 5

#### 1.3.1 Figure 5g

~~~
for t=0.95 (left panel)
Wilcoxon rank-sum (Mann–Whitney U) test results:
~~~

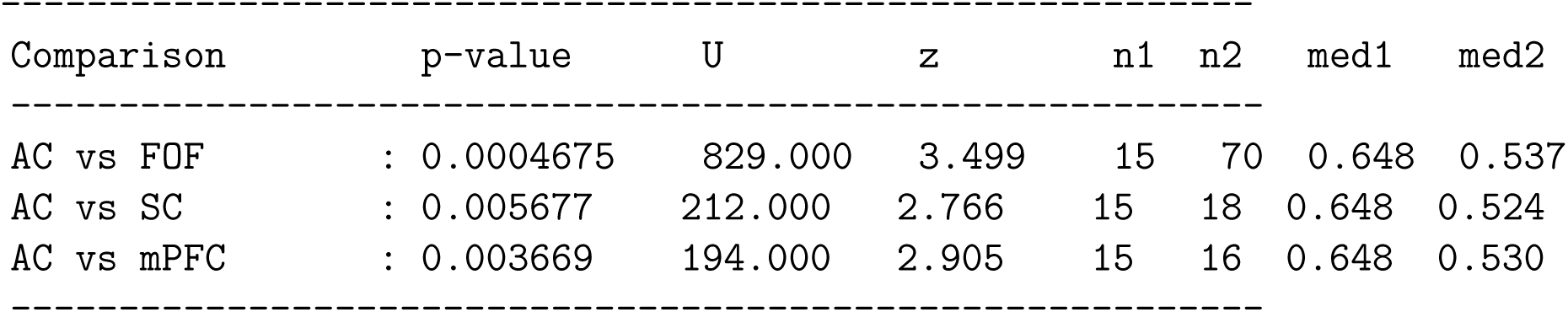

| Comparison |  | p-value | U | z | n1 | n2 | med1 | med2 |
|---|---|---|---|---|---|---|---|---|
| AC vs FOF | : | 0.0004675 | 829.000 | 3.499 | 15 | 70 | 0.648 | 0.537 |
| AC vs SC | : | 0.005677 | 212.000 | 2.766 | 15 | 18 | 0.648 | 0.524 |
| AC vs mPFC | : | 0.003669 | 194.000 | 2.905 | 15 | 16 | 0.648 | 0.530 |

~~~
for t=1.05 (right panel)
Wilcoxon rank-sum (Mann–Whitney U) test results:
~~~

| Comparison |  | p-value | U | z | n1 | n2 | med1 | med2 |
|---|---|---|---|---|---|---|---|---|
| AC vs FOF | : | 0.03391 | 709.500 | 2.121 | 15 | 70 | 0.597 | 0.551 |
| AC vs SC | : | 0.03762 | 193.000 | 2.079 | 15 | 18 | 0.597 | 0.524 |
| AC vs mPFC | : | 0.03796 | 173.000 | 2.075 | 15 | 16 | 0.597 | 0.536 |

## 1.4 Figure 7

### 1.4.1 Figure 7c

**Supplementary Table 7.** Mixed-effects likelihood-ratio test comparing movement time (MT) on each post-transformation day to baseline.

| Day | p-Val | LRStat | ΔDF | AIC<sub>*m*1</sub> | BIC<sub>*m*1</sub> | LogLik<sub>*m*1</sub> | AIC<sub>*m*0</sub> | BIC<sub>*m*0</sub> | LogLik<sub>*m*0</sub> |
|---|---|---|---|---|---|---|---|---|---|
| 0 | 0.0023833 | 9.2281 | 1 | -4013.31 | -3995.82 | 2009.66 | -4020.54 | -3997.22 | 2014.27 |
| 1 | 0.1072827 | 2.5938 | 1 | -5079.90 | -5061.46 | 2542.95 | -5080.50 | -5055.90 | 2544.25 |
| 2 | 2.2204e-16 | 67.5924 | 1 | -5068.71 | -5050.46 | 2537.36 | -5134.30 | -5109.97 | 2571.15 |
| 3 | < 10<sup>−9</sup> | 82.7915 | 1 | -4947.03 | -4928.93 | 2476.51 | -5027.82 | -5003.69 | 2517.91 |
| 4 | < 10<sup>−9</sup> | 143.4446 | 1 | -4276.74 | -4258.81 | 2141.37 | -4418.18 | -4394.28 | 2213.09 |
| 5 | < 10<sup>−9</sup> | 126.1329 | 1 | -4217.40 | -4199.36 | 2111.70 | -4341.54 | -4317.48 | 2174.77 |
| 6 | < 10<sup>−9</sup> | 165.5597 | 1 | -4344.47 | -4326.62 | 2175.23 | -4508.03 | -4484.24 | 2258.01 |
| 7 | < 10<sup>−9</sup> | 100.9583 | 1 | -4246.73 | -4228.90 | 2126.37 | -4345.69 | -4321.91 | 2176.84 |

#### 1.4.2 Figure 7e, Left panel, Allo task, Sham/FOF comparison

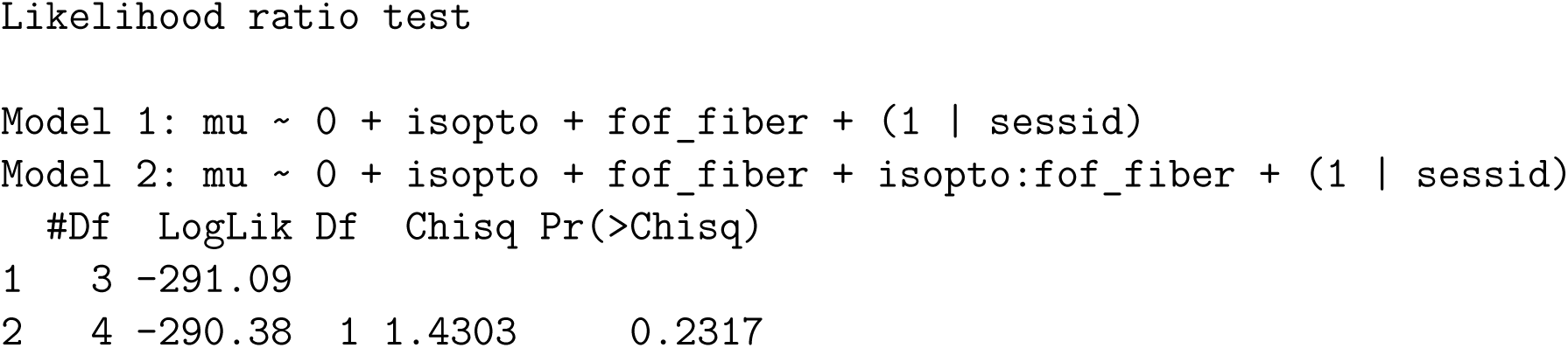

#### 1.4.3 Figure 7e, Right panel, 2AFC task, Sham/FOF comparison

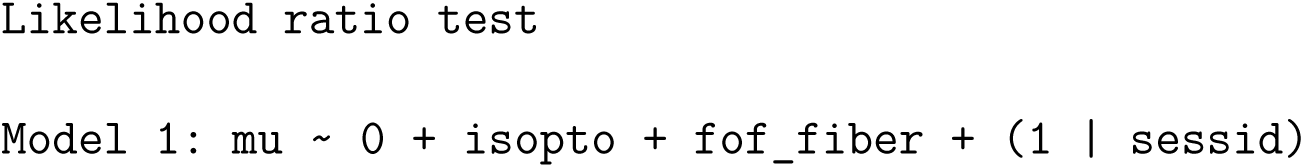
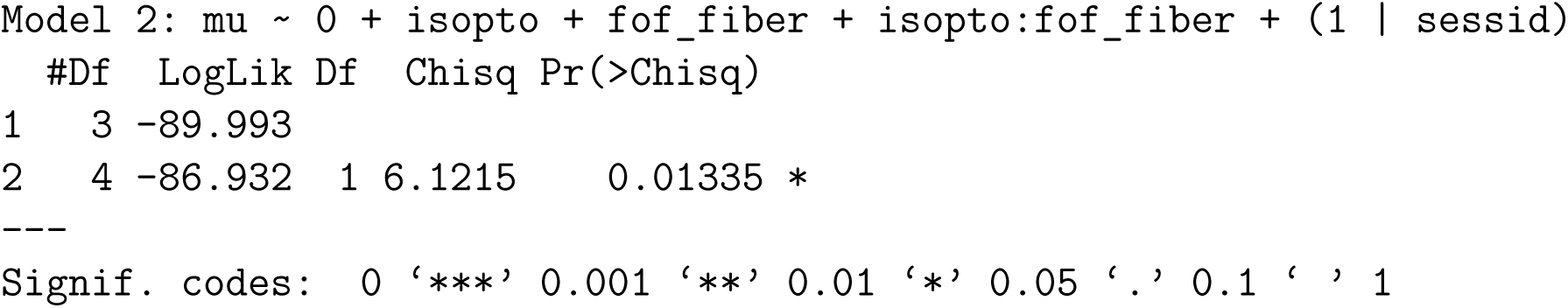

#### 1.4.4 Figure 7e, Main comparison between Allo/2AFC task

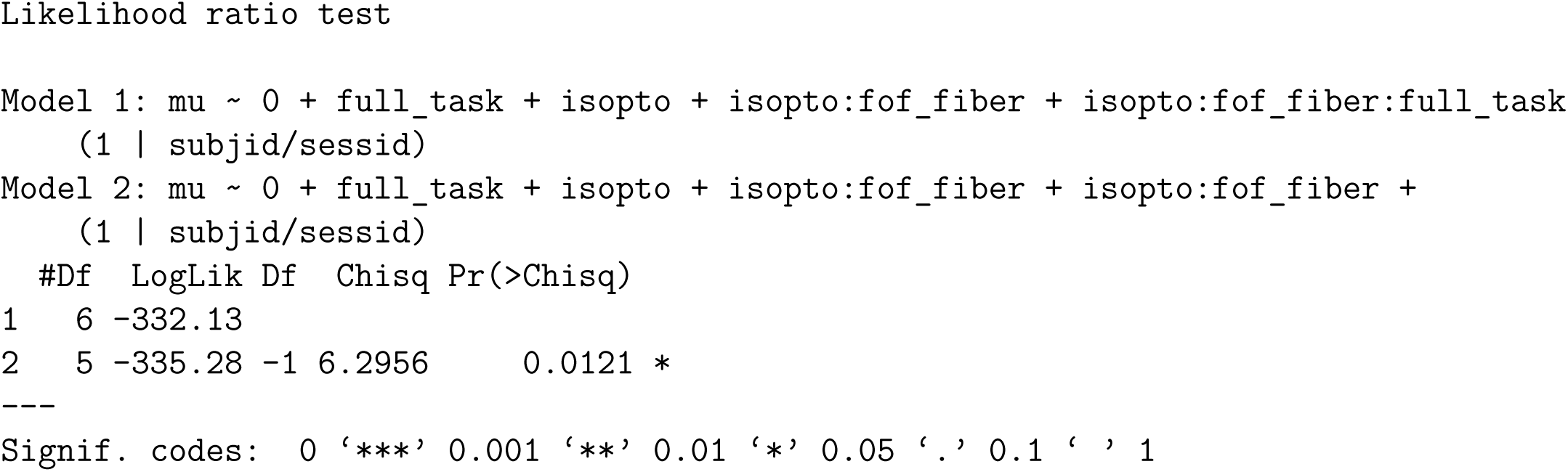

### 1.5 Supplementary figure 6

### 1.6 Supplementary **figure 6e**, Left panel, Ego task, Sham/FOF comparison

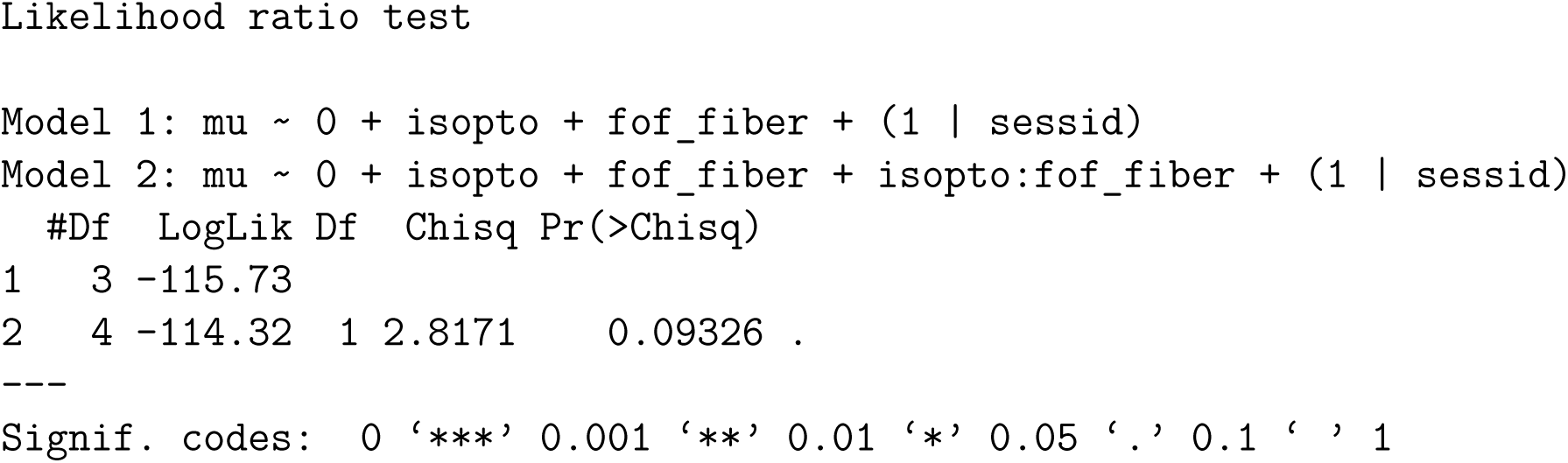

### 1.7 Supplementary figure 6e, Right panel, Allo task, Sham/FOF comparison

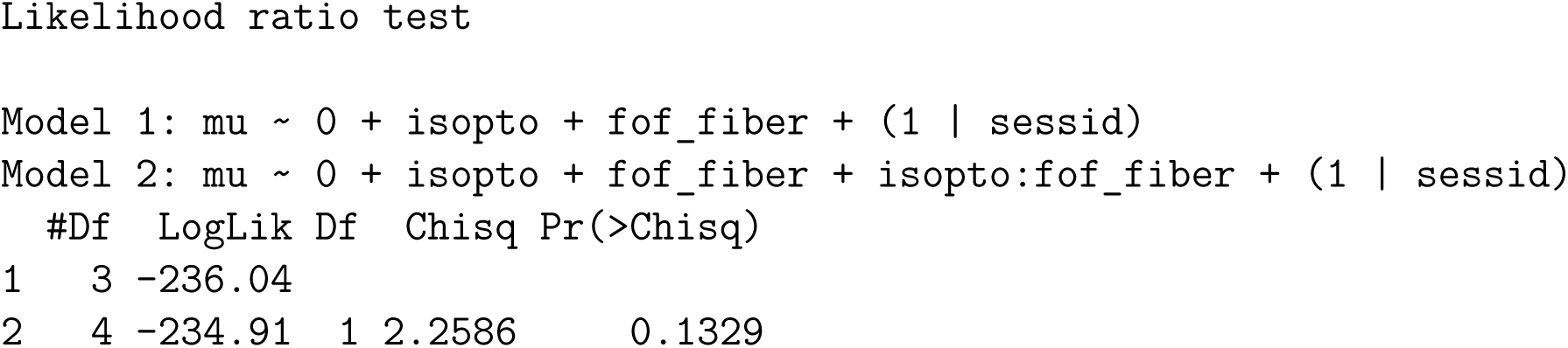

#### 1.7.1 Supplementary figure 6e, Main comparison between Allo/Ego task

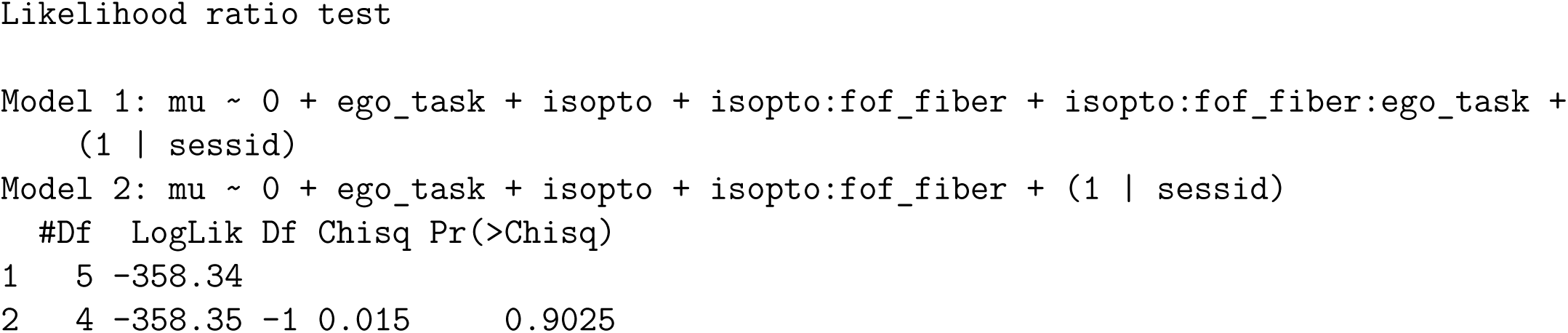

### 1.8 Supplementary Figure 6f, Left Panel, Ego task, Sham/FOF comparison

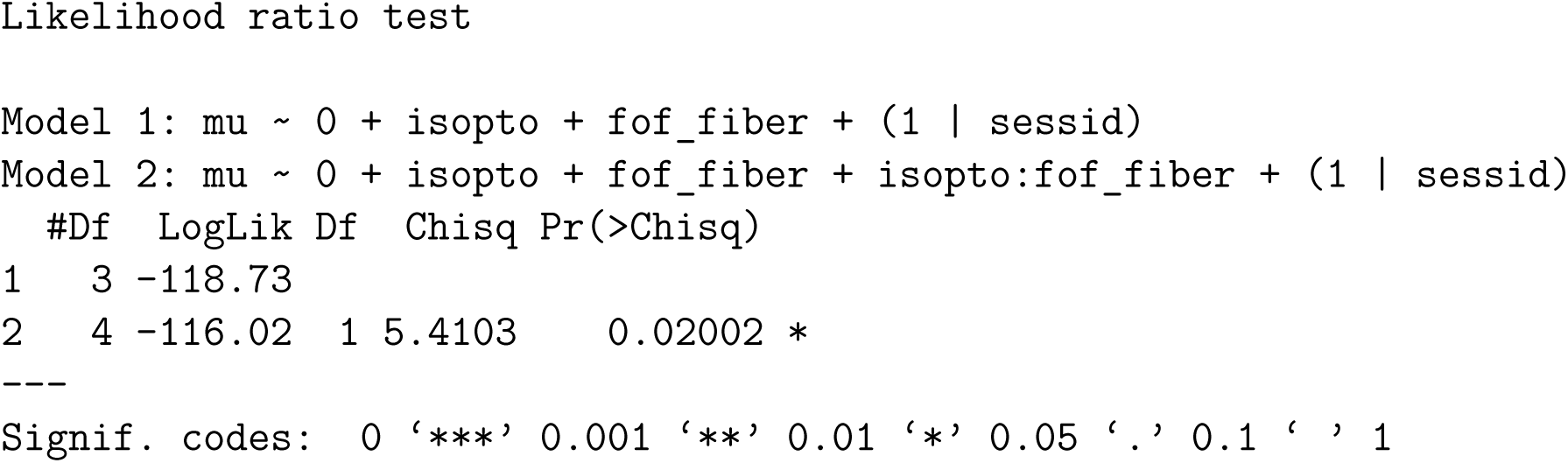

### 1.9 Supplementary Figure 6f, Right panel, Allo task, Sham/FOF comparison

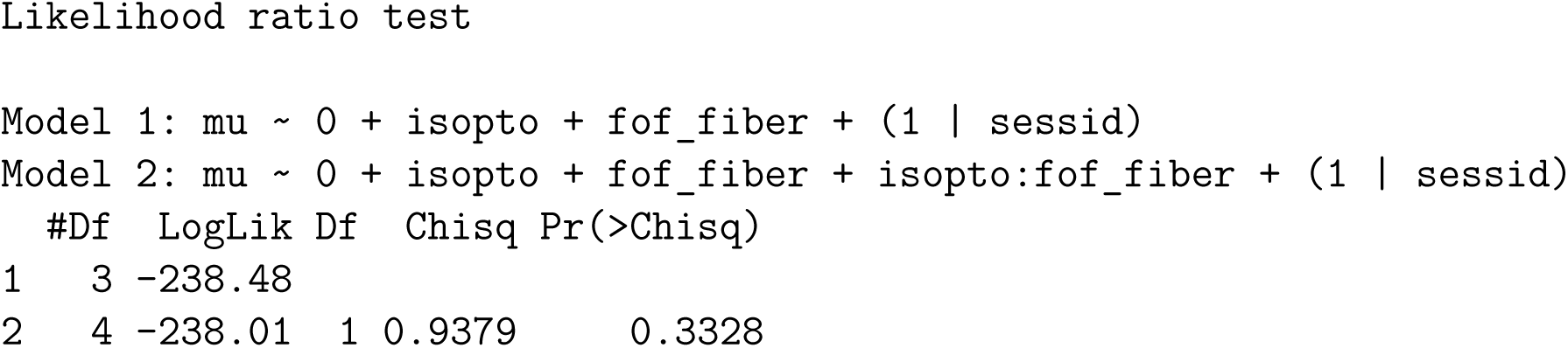

#### 1.9.1 Supplementary figure 6f, Main comparison between Allo/Ego task

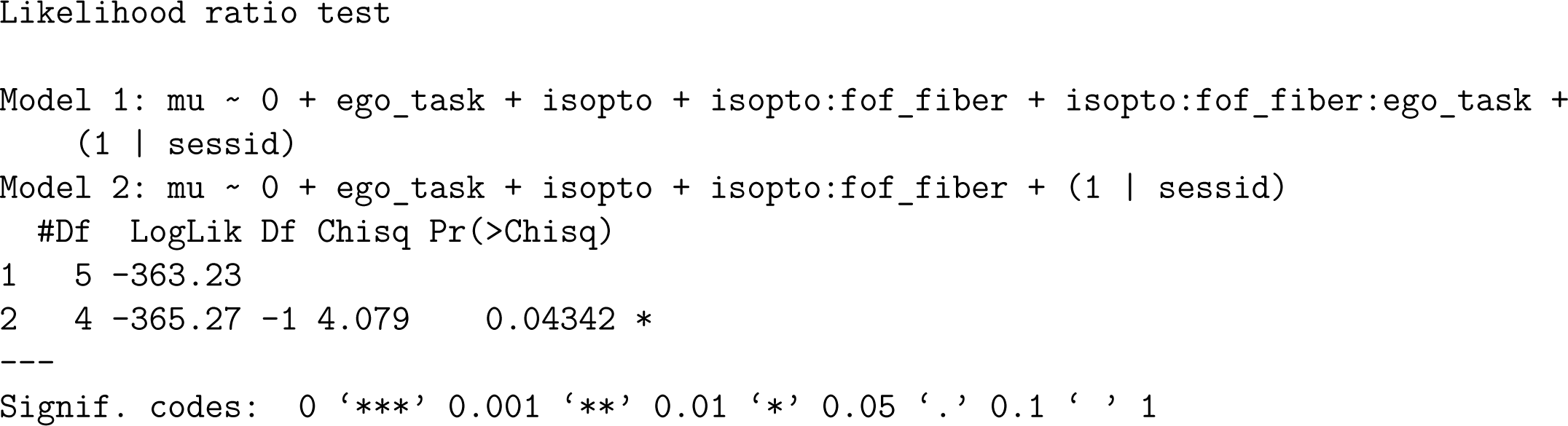

### 1.10 Supplementary figure 7

#### 1.10.1 Supplementary figure 7e, left Panel, Ego task, Sham/FOF comparison

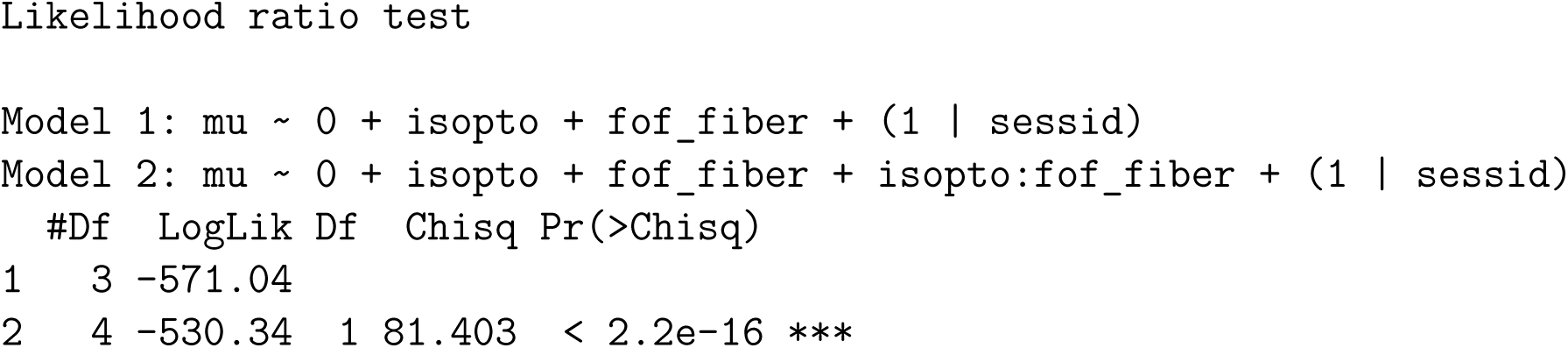

#### 1.10.2 Supplementary figure 7e, right Panel, Allo task, Sham/FOF comparison

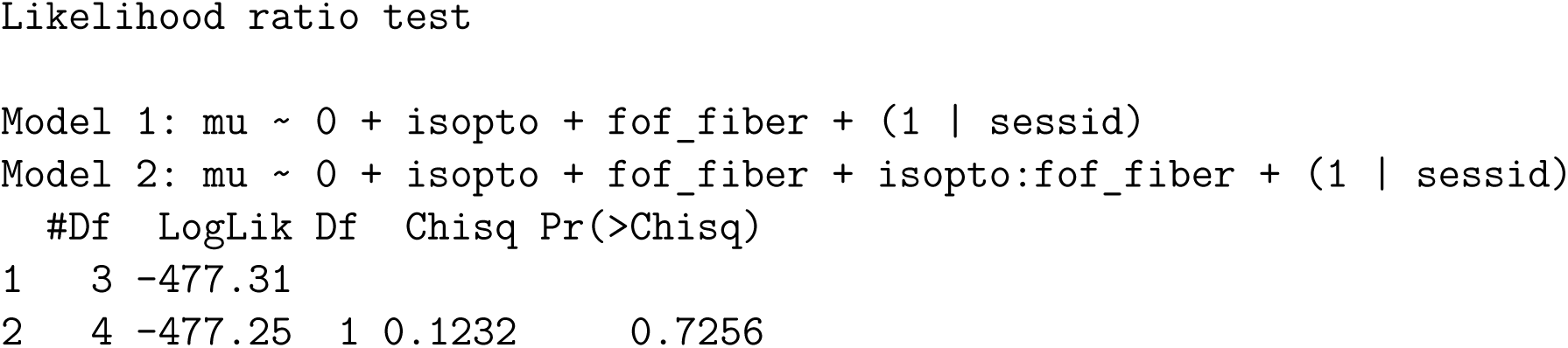

#### 1.10.3 Supplementary figure 7f, left Panel, Ego task, Sham/FOF comparison

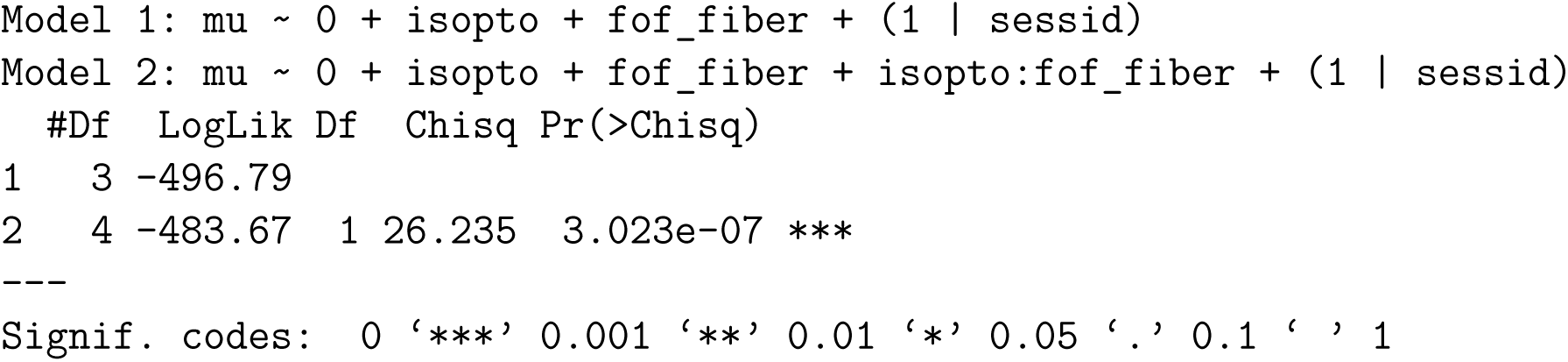

#### 1.10.4 Supplementary figure 7f, right Panel, Allo task, Sham/FOF comparison

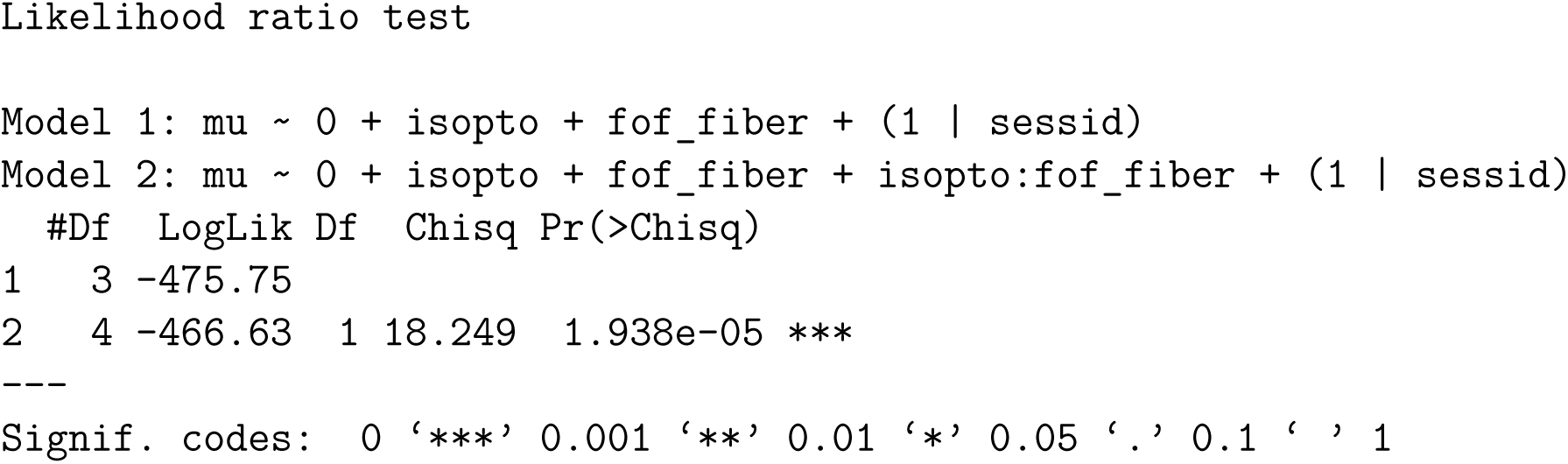

### 1.11 Supplementary figure 10

#### 1.11.1 Supplementary figure 10g hit trials

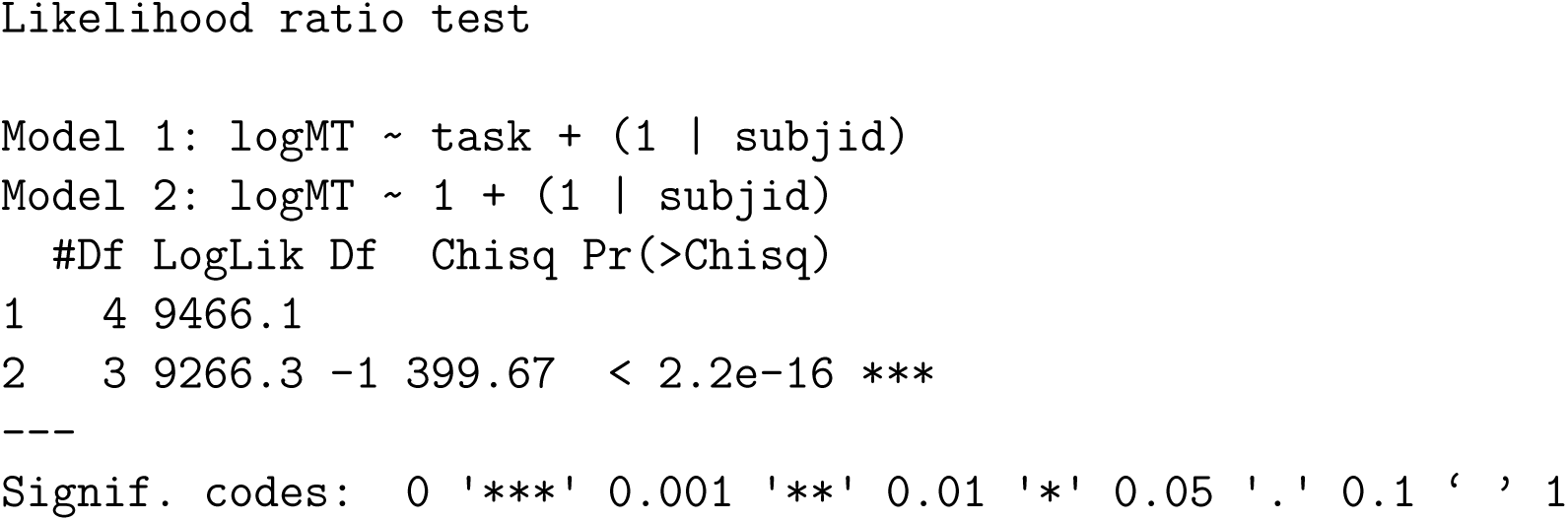

#### 1.11.2 Supplementary figure 10g error trials

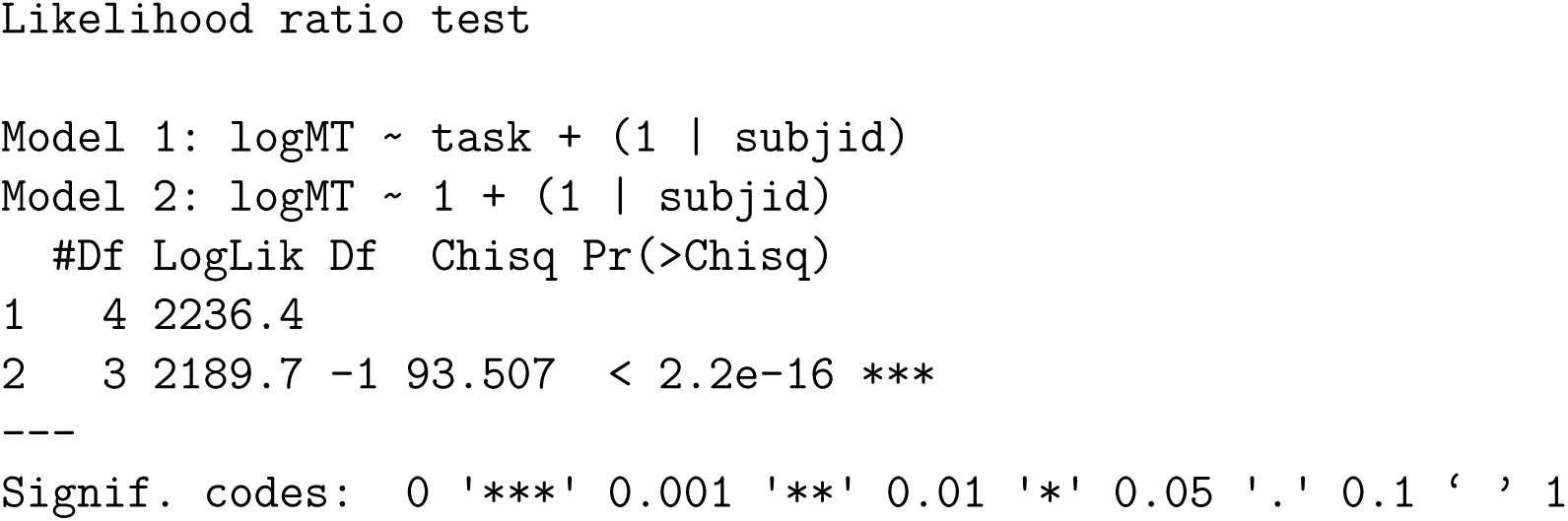

## References

[1] S. Funahashi, C. J. Bruce, and P. S. Goldman-Rakic. Mnemonic coding of visual space in the monkey’s dorsolateral prefrontal cortex. Journal of Neurophysiology, 61(2):331–349, February 1989. ISSN 0022-3077. doi: 10.1152/jn.1989.61.2.331.

[2] Joaquin M. Fuster and Garrett E. Alexander. Neuron Activity Related to ShortTerm Memory. Science, 173(3997):652–654, August 1971. ISSN 0036-8075, 1095-9203. doi: 10.1126/science.173.3997.652.

[3] Clayton E. Curtis and Mark D’Esposito. Persistent activity in the prefrontal cortex during working memory. Trends in Cognitive Sciences, 7(9):415–423, September 2003. ISSN 1364-6613. doi: 10.1016/s1364-6613(03)00197-9.

[4] Jeffrey C Erlich, Max Bialek, and Carlos D Brody. A cortical substrate for memory-guided orienting in the rat. Neuron, 72(2):330–343, October 2011. doi: 10.1016/j.neuron.2011.07.010.

[5] Nuo Li, Kayvon Daie, Karel Svoboda, and Shaul Druckmann. Robust neuronal dynamics in premotor cortex during motor planning. Nature, 532(7600):459–464, April 2016. ISSN 0028-0836. doi: 10.1038/nature17643.

[6] David A. Markowitz, Clayton E. Curtis, and Bijan Pesaran. Multiple component networks support working memory in prefrontal cortex. Proceedings of the National Academy of Sciences, 112(35):11084–11089, September 2015. ISSN 0027-8424, 1091–6490. doi: 10.1073/pnas.1504172112.

[7] Hidehiko K. Inagaki, Lorenzo Fontolan, Sandro Romani, and Karel Svoboda. Discrete attractor dynamics underlies persistent activity in the frontal cortex. Nature, 566(7743):212–217, February 2019. ISSN 1476-4687. doi: 10.1038/s41586-019-0919-7.

[8] Zengcai V. Guo, Hidehiko K. Inagaki, Kayvon Daie, Shaul Druckmann, Charles R. Gerfen, and Karel Svoboda. Maintenance of persistent activity in a frontal thalamocortical loop. Nature, 545(7653):181–186, May 2017. ISSN 0028-0836. doi: 10.1038/nature22324.

[9] Charles D Kopec, Jeffrey C Erlich, Bingni W Brunton, Karl Deisseroth, and Carlos D Brody. Cortical and Subcortical Contributions to Short-Term Memory for Orienting Movements. Neuron, 88(2):367–377, October 2015. doi: 10.1016/j.neuron.2015.08.033.

[10] Matthew V. Chafee and Patricia S. Goldman-Rakic. Inactivation of Parietal and Prefrontal Cortex Reveals Interdependence of Neural Activity During Memory-Guided Saccades. Journal of Neurophysiology, 83(3):1550–1566, March 2000. ISSN 0022-3077. doi: 10.1152/jn.2000.83.3.1550.

[11] Leah Acker, Erica N. Pino, Edward S. Boyden, and Robert Desimone. FEF inactivation with improved optogenetic methods. Proceedings of the National Academy of Sciences, 113(46):E7297–E7306, November 2016. ISSN 0027-8424, 1091-6490. doi: 10.1073/pnas.1610784113.

[12] Karel Svoboda and Nuo Li. Neural mechanisms of movement planning: Motor cortex and beyond. Current Opinion in Neurobiology, 49:33–41, April 2018. ISSN 09594388. doi: 10.1016/j.conb.2017.10.023.

[13] Hidehiko K. Inagaki, Susu Chen, Kayvon Daie, Arseny Finkelstein, Lorenzo Fontolan, Sandro Romani, and Karel Svoboda. Neural Algorithms and Circuits for Motor Planning. Annual Review of Neuroscience, 45(1):249–271, July 2022. ISSN 0147-006X, 1545-4126. doi: 10.1146/annurev-neuro-092021-121730.

[14] Ariel Gilad, Yasir Gallero-Salas, Dominik Groos, and Fritjof Helmchen. Behavioral Strategy Determines Frontal or Posterior Location of Short-Term Memory in Neocortex. Neuron, 99(4):814–828.e7, August 2018. ISSN 08966273. doi: 10.1016/j.neuron.2018.07.029.

[15] Liujunli Li, Timo Flesch, Ce Ma, Jingjie Li, Yizhou Chen, Hung-Tu Chen, and Jeffrey C. Erlich. Encoding of 2D self-centered plans and world-centered positions in the rat frontal orienting field. The Journal of Neuroscience, page e0018242024, August 2024. ISSN 0270-6474, 1529-2401. doi: 10.1523/JNEUROSCI.0018-24.2024.

[16] David J.-N. Maisson, Roberto Lopez Cervera, Benjamin Voloh, Indirah Conover, Mrunal Zambre, Jan Zimmermann, and Benjamin Y. Hayden. Widespread coding of navigational variables in prefrontal cortex. Current Biology, 33(16):3478–3488.e3, August 2023. ISSN 09609822. doi: 10.1016/j.cub.2023.07.024.

[17] Jacob M. Olson, Jamie K. Li, Sarah E. Montgomery, and Douglas A. Nitz. Secondary Motor Cortex Transforms Spatial Information into Planned Action during Navigation. Current Biology, page S0960982220303547, April 2020. ISSN 09609822. doi: 10.1016/j.cub.2020.03.016.

[18] Yasir Gallero-Salas, Shuting Han, Yaroslav Sych, Fabian F. Voigt, Balazs Laurenczy, Ariel Gilad, and Fritjof Helmchen. Sensory and Behavioral Components of Neocortical Signal Flow in Discrimination Tasks with Short-Term Memory. Neuron, 109(1):135–148.e6, January 2021. ISSN 08966273. doi: 10.1016/j.neuron.2020.10.017.

[19] Caterina Trentin, Chris Olivers, and Heleen A. Slagter. Action Planning Renders Objects in Working Memory More Attentionally Salient. Journal of Cognitive Neuroscience, 36(10):2166–2183, October 2024. ISSN 0898-929X, 1530-8898. doi: 10.1162/jocn_a_02235.

[20] Elise N. Mangin, Jian Chen, Jing Lin, and Nuo Li. Behavioral measurements of motor readiness in mice. Current Biology, 33(17):3610–3624.e4, September 2023. ISSN 09609822. doi: 10.1016/j.cub.2023.07.029.

[21] Alexa Riehle and Jean Requin. The predictive value for performance speed of preparatory changes in neuronal activity of the monkey motor and premotor cortex. Behavioural Brain Research, 53(1-2):35–49, February 1993. ISSN 01664328. doi: 10.1016/S0166-4328(05)80264-5.

[22] Ivan Voitov and Thomas D. Mrsic-Flogel. Cortical feedback loops bind distributed representations of working memory. Nature, 608(7922):381–389, August 2022. ISSN 0028-0836, 1476-4687. doi: 10.1038/s41586-022-05014-3.

[23] Chunyu A. Duan, Yuxin Pan, Guofen Ma, Taotao Zhou, Siyu Zhang, and Ninglong Xu. A cortico-collicular pathway for motor planning in a memory-dependent perceptual decision task. Nature Communications, 12(1):2727, May 2021. ISSN 2041-1723. doi: 10.1038/s41467-021-22547-9.

[24] M. Pare and D. P. Munoz. Saccadic reaction time in the monkey: Advanced preparation of oculomotor programs is primarily responsible for express saccade occurrence. Journal of Neurophysiology, 76(6):3666–3681, December 1996. ISSN 0022-3077, 1522-1598. doi: 10.1152/jn.1996.76.6.3666.

[25] Mark M. Churchland, Byron M. Yu, Stephen I. Ryu, Gopal Santhanam, and Krishna V. Shenoy. Neural Variability in Premotor Cortex Provides a Signature of Motor Preparation. The Journal of Neuroscience, 26(14):3697–3712, April 2006. ISSN 0270-6474, 1529-2401. doi: 10.1523/jneurosci.3762-05.2006.

[26] Munib A. Hasnain, Jaclyn E. Birnbaum, Juan Luis Ugarte Nunez, Emma K. Hartman, Chandramouli Chandrasekaran, and Michael N. Economo. Separating cognitive and motor processes in the behaving mouse. Nature Neuroscience, February 2025. ISSN 1097-6256, 1546-1726. doi: 10.1038/s41593-024-01859-1.

[27] Michael Krumin, Julie J Lee, Kenneth D Harris, and Matteo Carandini. Decision and navigation in mouse parietal cortex. eLife, 7:e42583, November 2018. ISSN 2050-084X. doi: 10.7554/eLife.42583.

[28] Diksha Gupta, Charles D. Kopec, Adrian G. Bondy, Thomas Z. Luo, Verity A. Elliott, and Carlos D. Brody. A multi-region recurrent circuit for evidence accumulation in rats, July 2024.

[29] R. Becket Ebitz and Benjamin Y. Hayden. The population doctrine in cognitive neuroscience. Neuron, 109(19):3055–3068, October 2021. ISSN 08966273. doi: 10.1016/j.neuron.2021.07.011.

[30] Hidehiko K. Inagaki, Susu Chen, Margreet C. Ridder, Pankaj Sah, Nuo Li, Zidan Yang, Hana Hasanbegovic, Zhenyu Gao, Charles R. Gerfen, and Karel Svoboda. A midbrain-thalamus-cortex circuit reorganizes cortical dynamics to initiate movement. Cell, 185(6):1065–1081.e23, March 2022. ISSN 00928674. doi: 10.1016/j.cell.2022.02.006.

[31] Stefano Panzeri, Monica Moroni, Houman Safaai, and Christopher D. Harvey. The structures and functions of correlations in neural population codes. Nature Reviews Neuroscience, 23(9):551–567, September 2022. ISSN 1471-003X, 1471-0048. doi: 10.1038/s41583-022-00606-4.

[32] Matthew L. Leavitt, Florian Pieper, Adam J. Sachs, and Julio C. Martinez-Trujillo. Correlated variability modifies working memory fidelity in primate prefrontal neuronal ensembles. Proceedings of the National Academy of Sciences, 114(12), March 2017. ISSN 0027-8424, 1091-6490. doi: 10.1073/pnas.1619949114.

[33] Valerio Mante, David Sussillo, Krishna V. Shenoy, and William T. Newsome. Context-dependent computation by recurrent dynamics in prefrontal cortex. Nature, 503(7474):78–84, November 2013. ISSN 0028-0836, 1476-4687. doi: 10.1038/nature12742.

[34] Mathias Mahn, Lihi Gibor, Pritish Patil, Katayun Cohen-Kashi Malina, Shir Oring, Yoav Printz, Rivka Levy, Ilan Lampl, and Ofer Yizhar. High-efficiency optogenetic silencing with soma-targeted anion-conducting channelrhodopsins. Nature Communications, 9(1):4125, October 2018. ISSN 2041-1723. doi: 10.1038/s41467-018-06511-8.

[35] Chunyu A Duan, Jeffrey C Erlich, and Carlos D Brody. Requirement of Prefrontal and Midbrain Regions for Rapid Executive Control of Behavior in the Rat. Neuron, 86(6):1491–1503, June 2015. doi: 10.1016/j.neuron.2015.05.042.

[36] Timothy D Hanks, Charles D Kopec, Bingni W Brunton, Chunyu A Duan, Jeffrey C Erlich, and Carlos D Brody. Distinct relationships of parietal and prefrontal cortices to evidence accumulation. Nature, 520(7546):220–223, April 2015. doi: 10.1038/nature14066.

[37] Nicholas A. Steinmetz, Peter Zatka-Haas, Matteo Carandini, and Kenneth D. Harris. Distributed coding of choice, action and engagement across the mouse brain. Nature, 576(7786):266–273, December 2019. ISSN 0028-0836, 1476-4687. doi: 10.1038/s41586-019-1787-x.

[38] Liping Yu, Jiawei Hu, Chenlin Shi, Li Zhou, Maozhi Tian, Jiping Zhang, and Jinghong Xu. The causal role of auditory cortex in auditory working memory. eLife, 10:e64457, April 2021. ISSN 2050-084X. doi: 10.7554/eLife.64457.

[39] Aman B. Saleem, E. Mika Diamanti, Julien Fournier, Kenneth D. Harris, and Matteo Carandini. Coherent encoding of subjective spatial position in visual cortex and hippocampus. Nature, 562(7725):124–127, October 2018. ISSN 0028-0836, 1476-4687. doi: 10.1038/s41586-018-0516-1.

[40] Jonathan R. Whitlock, Robert J. Sutherland, Menno P. Witter, May-Britt Moser, and Edvard I. Moser. Navigating from hippocampus to parietal cortex. Proceedings of the National Academy of Sciences, 105(39):14755–14762, September 2008. ISSN 0027-8424, 1091-6490. doi: 10.1073/pnas.0804216105.

[41] Andrew S. Alexander and Douglas A. Nitz. Retrosplenial cortex maps the conjunction of internal and external spaces. Nature Neuroscience, 18(8):1143–1151, August 2015. ISSN 1546-1726. doi: 10.1038/nn.4058.

[42] Andrew S. Alexander, Ryan Place, Michael J. Starrett, Elizabeth R. Chrastil, and Douglas A. Nitz. Rethinking retrosplenial cortex: Perspectives and predictions. Neuron, page S0896627322010273, December 2022. ISSN 08966273. doi: 10.1016/j.neuron.2022.11.006.

[43] Tsukasa Kamigaki and Yang Dan. Delay activity of specific prefrontal interneuron subtypes modulates memory-guided behavior. Nature Neuroscience, 20(6):854–863, June 2017. ISSN 1097-6256, 1546-1726. doi: 10.1038/nn.4554.

[44] David R. Euston, Aaron J. Gruber, and Bruce L. McNaughton. The Role of Medial Prefrontal Cortex in Memory and Decision Making. Neuron, 76(6):1057–1070, December 2012. ISSN 0896-6273. doi: 10.1016/j.neuron.2012.12.002.

[45] Dong-Hyun Lim, Young Ju Yoon, Eunsil Her, Suehee Huh, and Min Whan Jung. Active maintenance of eligibility trace in rodent prefrontal cortex. Scientific Reports, 10(1):18860, November 2020. ISSN 2045-2322. doi: 10.1038/s41598-020-75820-0.

[46] Veronika Samborska, James L. Butler, Mark E. Walton, Timothy E. J. Behrens, and Thomas Akam. Complementary task representations in hippocampus and prefrontal cortex for generalizing the structure of problems. Nature Neuroscience, 25(10):1314–1326, October 2022. ISSN 1097-6256, 1546-1726. doi: 10.1038/s41593-022-01149-8.

[47] Chunyu A. Duan, Marino Pagan, Alex T. Piet, Charles D. Kopec, Athena Akrami, Alexander J. Riordan, Jeffrey C. Erlich, and Carlos D. Brody. Collicular circuits for flexible sensorimotor routing. Nature Neuroscience, 24(8):1110–1120, August 2021. ISSN 1097-6256, 1546-1726. doi: 10.1038/s41593-021-00865-x.

[48] Xiang Gao and Michael Wehr. A Coding Transformation for Temporally Structured Sounds within Auditory Cortical Neurons. Neuron, 86(1):292–303, April 2015. ISSN 08966273. doi: 10.1016/j.neuron.2015.03.004.

[49] Raphael Steinfeld, André Tacão-Monteiro, and Alfonso Renart. Differential representation of sensory information and behavioral choice across layers of the mouse auditory cortex. Current Biology, 34(10):2200–2211.e6, May 2024. ISSN 09609822. doi: 10.1016/j.cub.2024.04.040.

[50] Alexandra Libby and Timothy J. Buschman. Rotational dynamics reduce interference between sensory and memory representations. Nature Neuroscience, 24(5): 715–726, May 2021. ISSN 1097-6256, 1546-1726. doi: 10.1038/s41593-021-00821-9.

[51] Florent Barthas and Alex C. Kwan. Secondary Motor Cortex: Where ‘Sensory’ Meets ‘Motor’ in the Rodent Frontal Cortex. Trends in Neurosciences, 40(3):181– 193, March 2017. ISSN 0166-2236. doi: 10.1016/j.tins.2016.11.006.

[52] Ranulfo Romo, Carlos D. Brody, Adrián Hernández, and Luis Lemus. Neuronal correlates of parametric working memory in the prefrontal cortex. Nature, 399 (6735):470–473, June 1999. ISSN 1476-4687. doi: 10.1038/20939.

[53] Vahid Esmaeili and Mathew E. Diamond. Neuronal Correlates of Tactile Working Memory in Prefrontal and Vibrissal Somatosensory Cortex. Cell Reports, 27(11): 3167–3181.e5, June 2019. ISSN 22111247. doi: 10.1016/j.celrep.2019.05.034.

[54] Joseph T. McGuire and Matthew M. Botvinick. Prefrontal cortex, cognitive control, and the registration of decision costs. Proceedings of the National Academy of Sciences, 107(17):7922–7926, April 2010. ISSN 0027-8424, 1091-6490. doi: 10.1073/pnas.0910662107.

[55] Mark K. Ho, David Abel, Carlos G. Correa, Michael L. Littman, Jonathan D. Cohen, and Thomas L. Griffiths. People construct simplified mental representations to plan. Nature, 606(7912):129–136, June 2022. ISSN 0028-0836, 1476-4687. doi: 10.1038/s41586-022-04743-9.

[56] Trinity B. Crapse and Marc A. Sommer. Corollary discharge across the animal kingdom. Nature Reviews Neuroscience, 9(8):587–600, August 2008. ISSN 1471-0048. doi: 10.1038/nrn2457.

[57] Sten Grillner. Evolution of the vertebrate motor system — from forebrain to spinal cord. Current Opinion in Neurobiology, 71:11–18, December 2021. ISSN 09594388. doi: 10.1016/j.conb.2021.07.016.

[58] Sten Grillner. The Brain in Motion: From Microcircuits to Global Brain Function. The MIT Press, Cambridge London, 2023. ISBN 978-0-262-37531-3 978-0-262-37530-6.

[59] Peter Carruthers. Evolution of working memory. Proceedings of the National Academy of Sciences, 110(supplement_2):10371–10378, June 2013. ISSN 00278424, 1091-6490. doi: 10.1073/pnas.1301195110.

[60] Rifqi O. Affan, Ian M. Bright, Luke N. Pemberton, Nathanael A. Cruzado, Benjamin B. Scott, and Marc W. Howard. Ramping dynamics in the frontal cortex unfold over multiple timescales during motor planning. Journal of Neurophysiology, 133(2):625–637, February 2025. ISSN 0022-3077, 1522-1598. doi: 10.1152/jn.00234.2024.

[61] C. K. Machens, R. Romo, and C. D. Brody. Functional, But Not Anatomical, Separation of “What” and “When” in Prefrontal Cortex. Journal of Neuroscience, 30(1): 350–360, January 2010. ISSN 0270-6474, 1529-2401. doi: 10.1523/JNEUROSCI.3276-09.2010.

[62] Aaron A. Wilber, Benjamin J. Clark, Tyler C. Forster, Masami Tatsuno, and Bruce L. McNaughton. Interaction of Egocentric and World-Centered Reference Frames in the Rat Posterior Parietal Cortex. Journal of Neuroscience, 34(16): 5431–5446, April 2014. ISSN 0270-6474, 1529-2401. doi: 10.1523/JNEUROSCI.0511-14.2014.

[63] Dario Campagner, Ruben Vale, Yu Lin Tan, Panagiota Iordanidou, Oriol Pavón Arocas, Federico Claudi, A. Vanessa Stempel, Sepiedeh Keshavarzi, Rasmus S. Petersen, Troy W. Margrie, and Tiago Branco. A cortico-collicular circuit for orienting to shelter during escape. Nature, 613(7942):111–119, January 2023. ISSN 0028-0836, 1476-4687. doi: 10.1038/s41586-022-05553-9.

[64] Joaquín M. Fuster. The Prefrontal Cortex—An Update. Neuron, 30(2):319–333, May 2001. ISSN 08966273. doi: 10.1016/S0896-6273(01)00285-9.

[65] Elad Avidan, Shlomo David Sherer, and Ariel Gilad. Interhemispheric transfer of sensory and working memory information is dictated by behavioral strategy, March 2026.

[66] Renana Oz Rokach, Odeya Marmor, Yoav Levy, and Ariel Gilad. Brain-wide pathways for object identity and location are dominated by internal strategies. Nature Communications, October 2026. ISSN 2041-1723. doi: 10.1038/s41467-026-78597-4.

[67] David Hocker, Christine M. Constantinople, and Cristina Savin. Compositional pretraining improves computational efficiency and matches animal behaviour on complex tasks. Nature Machine Intelligence, 7(5):689–702, May 2025. ISSN 25225839. doi: 10.1038/s42256-025-01029-3.

[68] Daniel R. Kepple, Rainer Engelken, and Kanaka Rajan. Curriculum learning as a tool to uncover learning principles in the brain. In International Conference on Learning Representations, 2022.

[69] Matthew T. Kaufman, Mark M. Churchland, Stephen I. Ryu, and Krishna V. Shenoy. Cortical activity in the null space: Permitting preparation without movement. Nature Neuroscience, 17(3):440–448, March 2014. ISSN 1546-1726. doi: 10.1038/nn.3643.

[70] Klaus Wimmer, Duane Q Nykamp, Christos Constantinidis, and Albert Compte. Bump attractor dynamics in prefrontal cortex explains behavioral precision in spatial working memory. Nature Neuroscience, 17(3):431–439, March 2014. ISSN 1097-6256, 1546-1726. doi: 10.1038/nn.3645.

[71] Elizabeth S. Lorenc, Remington Mallett, and Jarrod A. Lewis-Peacock. Distraction in Visual Working Memory: Resistance is Not Futile. Trends in Cognitive Sciences, 25(3):228–239, March 2021. ISSN 13646613. doi: 10.1016/j.tics.2020.12.004.

[72] Margaret M Henderson, Rosanne L Rademaker, and John T Serences. Flexible utilization of spatialand motor-based codes for the storage of visuo-spatial information. eLife, 11:e75688, May 2022. ISSN 2050-084X. doi: 10.7554/eLife.75688.

[73] Freek Van Ede. Visual working memory and action: Functional links and bidirectional influences. Visual Cognition, 28(5-8):401–413, September 2020. ISSN 1350-6285, 1464-0716. doi: 10.1080/13506285.2020.1759744.

[74] Nicholas E. Myers, Mark G. Stokes, and Anna C. Nobre. Prioritizing Information during Working Memory: Beyond Sustained Internal Attention. Trends in Cognitive Sciences, 21(6):449–461, June 2017. ISSN 13646613. doi: 10.1016/j.tics.2017.03.010.

[75] P. Znamenskiy and A. M. Zador. Corticostriatal neurons in auditory cortex drive decisions during auditory discrimination. Nature, 497:482–485, May 2013. ISSN 1476-4687 (Electronic)0028-0836 (Linking).

[76] Joao Barbosa, Heike Stein, Rebecca L. Martinez, Adrià Galan-Gadea, Sihai Li, Josep Dalmau, Kirsten C. S. Adam, Josep Valls-Solé, Christos Constantinidis, and Albert Compte. Interplay between persistent activity and activity-silent dynamics in the prefrontal cortex underlies serial biases in working memory. Nature Neuroscience, 23(8):1016–1024, August 2020. ISSN 1097-6256, 1546-1726. doi: 10.1038/s41593-020-0644-4.

[77] Mark G. Stokes. ‘Activity-silent’ working memory in prefrontal cortex: A dynamic coding framework. Trends in Cognitive Sciences, 19(7):394–405, July 2015. ISSN 13646613. doi: 10.1016/j.tics.2015.05.004.

[78] Falk Lieder and Thomas L. Griffiths. Resource-rational analysis: Understanding human cognition as the optimal use of limited computational resources. Behavioral and Brain Sciences, 43, 2020. ISSN 0140-525X, 1469-1825. doi: 10.1017/s0140525x1900061x.

[79] Samuel J. Gershman, Eric J. Horvitz, and Joshua B. Tenenbaum. Computational rationality: A converging paradigm for intelligence in brains, minds, and machines. Science, 349(6245):273–278, July 2015. ISSN 0036-8075, 1095-9203. doi: 10.1126/science.aac6076.

[80] Shuze Liu, Samuel J. Gershman, and Bilal A. Bari. Quantifying the Cost of Context Sensitivity in Decision-Making. Topics in Cognitive Science, page tops.70030, October 2025. ISSN 1756-8757, 1756-8765. doi: 10.1111/tops.70030.

[81] Stephen M. Fleming. Metacognition and Confidence: A Review and Synthesis. Annual Review of Psychology, 75(1):241–268, January 2024. ISSN 0066-4308, 1545-2085. doi: 10.1146/annurev-psych-022423-032425.

[82] Tobias U. Hauser, Eran Eldar, and Raymond J. Dolan. Separate mesocortical and mesolimbic pathways encode effort and reward learning signals. Proceedings of the National Academy of Sciences, 114(35), August 2017. ISSN 0027-8424, 1091-6490. doi: 10.1073/pnas.1705643114.

[83] Amitai Shenhav, Matthew M. Botvinick, and Jonathan D. Cohen. The Expected Value of Control: An Integrative Theory of Anterior Cingulate Cortex Function. Neuron, 79(2):217–240, July 2013. ISSN 08966273. doi: 10.1016/j.neuron.2013.07.007.

[84] Eliana Vassena, Clay B. Holroyd, and William H. Alexander. Computational Models of Anterior Cingulate Cortex: At the Crossroads between Prediction and Effort. Frontiers in Neuroscience, 11:316, June 2017. ISSN 1662-453X. doi: 10.3389/fnins.2017.00316.

[85] Asako Mitsuto, Rei Akaishi, Keiichi Onoda, Kenji Morita, Toshikazu Kawagoe, Tetsuya Yamamoto, Shuhei Yamaguchi, Ritsuko Hanajima, and Andrew Westbrook. Mental Effort Cost Learning is Retrospective, October 2025.

[86] Sarah L. Master, Shanshan Li, and Clayton E. Curtis. Trying Harder: How Cognitive Effort Sculpts Neural Representations during Working Memory. The Journal of Neuroscience, 44(28):e0060242024, July 2024. ISSN 0270-6474, 1529-2401. doi: 10.1523/JNEUROSCI.0060-24.2024.

[87] Ceyda Sayalı and David Badre. Neural systems underlying the learning of cognitive effort costs. Cognitive, Affective, & Behavioral Neuroscience, 21(4):698–716, August 2021. ISSN 1530-7026, 1531-135X. doi: 10.3758/s13415-021-00893-x.

[88] George Paxinos and Charles Watson. The Rat Brain in Stereotaxic Coordinates. Acad. Press, San Diego [u.a., 2004. ISBN 978-0-12-547617-1.

[89] Flora Bouchacourt and Timothy J. Buschman. A Flexible Model of Working Memory. Neuron, 103(1):147–160.e8, July 2019. ISSN 08966273. doi: 10.1016/j.neuron.2019.04.020.

[90] Alexander N. Ford, Jordyn E. Czarny, Meike M. Rogalla, Gunnar L. Quass, and Pierre F. Apostolides. Auditory Corticofugal Neurons Transmit Auditory and Non-auditory Information During Behavior. The Journal of Neuroscience, 44(7):e1190232023, February 2024. ISSN 0270-6474, 1529-2401. doi: 10.1523/JNEUROSCI.1190-23.2023.

[91] John D. Murray, Jorge Jaramillo, and Xiao-Jing Wang. Working Memory and Decision-Making in a Frontoparietal Circuit Model. The Journal of Neuroscience, 37(50):12167–12186, December 2017. ISSN 0270-6474, 1529-2401. doi: 10.1523/JNEUROSCI.0343-17.2017.

[92] Alex L. White, Martin Rolfs, and Marisa Carrasco. Stimulus competition mediates the joint effects of spatial and feature-based attention. Journal of Vision, 15(14): 7, October 2015. ISSN 1534-7362. doi: 10.1167/15.14.7.

[93] Bo Shen, Duc Nguyen, Jailyn Wilson, Paul W. Glimcher, and Kenway Louie. Early versus late noise differentially enhances or degrades context-dependent choice. Nature Communications, 16(1):3828, April 2025. ISSN 2041-1723. doi: 10.1038/s41467-025-59140-3.

[94] Chaofei Bao, Xiaoyue Zhu, Joshua Mōller-Mara, Jingjie Li, Sylvain Dubroqua, and Jeffrey C. Erlich. The rat frontal orienting field dynamically encodes value for economic decisions under risk. Nature Neuroscience, 26(11):1942–1952, November 2023. ISSN 1097-6256, 1546-1726. doi: 10.1038/s41593-023-01461-x.

[95] Mihály Vöröslakos, Peter C Petersen, Balázs Vöröslakos, and György Buzsáki. Metal microdrive and head cap system for silicon probe recovery in freely moving rodent. eLife, 10:e65859, May 2021. ISSN 2050-084X. doi: 10.7554/eLife.65859.

[96] Marius Pachitariu, Nicholas Steinmetz, Shabnam Kadir, Matteo Carandini, and Harris Kenneth D. Kilosort: Realtime spike-sorting for extracellular electrophysiology with hundreds of channels, June 2016.

[97] Marius Pachitariu, Shashwat Sridhar, Jacob Pennington, and Carsen Stringer. Spike sorting with Kilosort4. Nature Methods, 21(5):914–921, May 2024. ISSN 1548-7091, 1548-7105. doi: 10.1038/s41592-024-02232-7.

[98] Xiaoxuan Jia, Joshua H. Siegle, Corbett Bennett, Samuel D. Gale, Daniel J. Denman, Christof Koch, and Shawn R. Olsen. High-density extracellular probes reveal dendritic backpropagation and facilitate neuron classification. Journal of Neurophysiology, 121(5):1831–1847, May 2019. ISSN 0022-3077, 1522-1598. doi: 10.1152/jn.00680.2018.

[99] D. Mayerich, L. Abbott, and B. McCORMICK. Knife-edge scanning microscopy for imaging and reconstruction of three-dimensional anatomical structures of the mouse brain. Journal of Microscopy, 231(1):134–143, July 2008. ISSN 0022-2720, 1365-2818. doi: 10.1111/j.1365-2818.2008.02024.x.

[100] Timothy Ragan, Lolahon R Kadiri, Kannan Umadevi Venkataraju, Karsten Bahlmann, Jason Sutin, Julian Taranda, Ignacio Arganda-Carreras, Yongsoo Kim, H Sebastian Seung, and Pavel Osten. Serial two-photon tomography for automated ex vivo mouse brain imaging. Nature Methods, 9(3):255–258, March 2012. ISSN 1548-7091, 1548-7105. doi: 10.1038/nmeth.1854.

[101] Douglas Bates, Martin Mächler, Ben Bolker, and Steve Walker. Fitting Linear Mixed-Effects Models Using lme4. Journal of Statistical Software, 67(1):1–48, October 2015. ISSN 1548-7660. doi: 10.18637/jss.v067.i01.

[102] Talmo D. Pereira, Nathaniel Tabris, Arie Matsliah, David M. Turner, Junyu Li, Shruthi Ravindranath, Eleni S. Papadoyannis, Edna Normand, David S. Deutsch, Z. Yan Wang, Grace C. McKenzie-Smith, Catalin C. Mitelut, Marielisa Diez Castro, John D’Uva, Mikhail Kislin, Dan H. Sanes, Sarah D. Kocher, Samuel S.-H. Wang, Annegret L. Falkner, Joshua W. Shaevitz, and Mala Murthy. SLEAP: A deep learning system for multi-animal pose tracking. Nature Methods, 19(4):486–495, April 2022. ISSN 1548-7091, 1548-7105. doi: 10.1038/s41592-022-01426-1.

[103] Jeffrey Erlich, Josh Moller-Mara, and Jingjie Li. Erlichlab/elutils: First Zenodo Release. Zenodo, May 2021.

[104] Ethan M. Meyers, David J. Freedman, Gabriel Kreiman, Earl K. Miller, and Tomaso Poggio. Dynamic Population Coding of Category Information in Inferior Temporal and Prefrontal Cortex. Journal of Neurophysiology, 100(3):1407–1419, September 2008. ISSN 0022-3077, 1522-1598. doi: 10.1152/jn.90248.2008.

[105] J-R. King and S. Dehaene. Characterizing the dynamics of mental representations: The temporal generalization method. Trends in Cognitive Sciences, 18(4):203–210, April 2014. ISSN 13646613. doi: 10.1016/j.tics.2014.01.002.

[106] Mike X Cohen. Analyzing Neural Time Series Data: Theory and Practice. The MIT Press, January 2014. ISBN 978-0-262-31955-3. doi: 10.7551/mitpress/9609.001.0001.

[107] M. Pagan and N. C. Rust. Dynamic Target Match Signals in Perirhinal Cortex Can Be Explained by Instantaneous Computations That Act on Dynamic Input from Inferotemporal Cortex. Journal of Neuroscience, 34(33):11067–11084, August 2014. ISSN 0270-6474, 1529-2401. doi: 10.1523/JNEUROSCI.4040-13.2014.

[108] Hadley Wickham. Ggplot2: Elegant Graphics for Data Analysis. Use R! Springer International Publishing: Imprint: Springer, Cham, 2nd ed. 2016 edition, 2016. ISBN 978-3-319-24277-4. doi: 10.1007/978-3-319-24277-4.

[109] Achim Zeileis and Torsten Hothorn. Diagnostic checking in regression relationships. R News, 2(3):7–10, 2002.

[110] G. N. Wilkinson and C. E. Rogers. Symbolic Description of Factorial Models for Analysis of Variance. Applied Statistics, 22(3):392, 1973. ISSN 00359254. doi: 10.2307/2346786.

[111] Andrew Gelman and Hal Stern. The Difference Between “Significant” and “Not Significant” is not Itself Statistically Significant. The American Statistician, 60(4):328– 331, November 2006. ISSN 0003-1305, 1537-2731. doi: 10.1198/000313006X152649.

[112] Alex T. Piet, Jeffrey C. Erlich, Charles D. Kopec, and Carlos D. Brody. Rat Prefrontal Cortex Inactivations during Decision Making Are Explained by Bistable Attractor Dynamics. Neural Computation, 29(11):2861–2886, August 2017. ISSN 0899-7667. doi: 10.1162/neco_a_01005.

[113] H. R. Wilson and J. D. Cowan. A mathematical theory of the functional dynamics of cortical and thalamic nervous tissue. Kybernetik, 13(2):55–80, September 1973. ISSN 0023-5946. doi: 10.1007/BF00288786.

[114] Alain Destexhe and Terrence J. Sejnowski. The Wilson–Cowan model, 36 years later. Biological Cybernetics, 101(1):1–2, July 2009. ISSN 0340-1200, 1432-0770. doi: 10.1007/s00422-009-0328-3.

